# Extending Comparison Methods for Unsigned Networks to Signed Networks

**DOI:** 10.1101/2022.09.23.509251

**Authors:** William Krinsman

## Abstract

We can allow the edges of networks to have both negative and positive weights. For example, signed networks can describe the interactions of microbes. To evaluate the performance of estimators for signed networks, we need quantitative comparison methods for signed networks. Finding such comparison methods is done most easily by extending a comparison method for unsigned networks.

Almost all methods reported in the literature for quantitatively comparing networks implicitly assume that edge weights are non-negative. Naive attempts to modify these methods to be applicable to signed networks can lead to nonsensical conclusions. Herein I identify requirements that should be satisfied by reasonable methods for comparing signed networks, most importantly the “double penalization principle”. I extend several comparison methods for unsigned networks while satisfying these requirements. Finally, I give examples where these extensions behave reasonably but naive extensions do not.

## INTRODUCTION

This paper highlights the work presented in chapter 8 of the PhD Thesis (Krinsman,2022).

### Networks with Mixed-Sign Edge Weights in Microbial Ecology

Networks with both positive and negative edge weights, i.e. mixed-sign edge weights, are a straightforward extension of typical networks (whose edge weights are only positive). Being straightforward, such signed networks are attested in the literature at least as early as (Harary,1953). Networks with mixed-sign edge weights appear to be used ubiquitously in ecological models of population growth, but the usage is implicit. Therefore we now explain how signed networks are used in (microbial) ecology.

Given a microbial community with *S* strains, we can define a network with mixed-sign edge weights as follows. Connect an edge from strain *s*_1_ to strain *s*_2_ if strain *s*_1_ influences the growth of strain *s*_2_, otherwise don’t connect them. If strain *s*_1_ promotes the growth of strain *s*_2_, make the sign of the edge weight positive. If strain *s*_1_ inhibits the growth of strain *s*_2_, make the sign of the edge weight negative. Assign a magnitude to the edge weight corresponding to the strength of the interaction. Cf. figure 1. For example, if the population growth dynamics of a microbial community are modelled using a system of differential equations, cf. (Gonze et al.,2018) for a review, then the edge weights can correspond to the equations’ coefficients. A similar approach found in (Xiao et al.,2017) and (Angulo et al.,2019) treats the values of the Jacobian of the system of differential equations at an equilibrium point as the edge weights.

**Figure 1.**
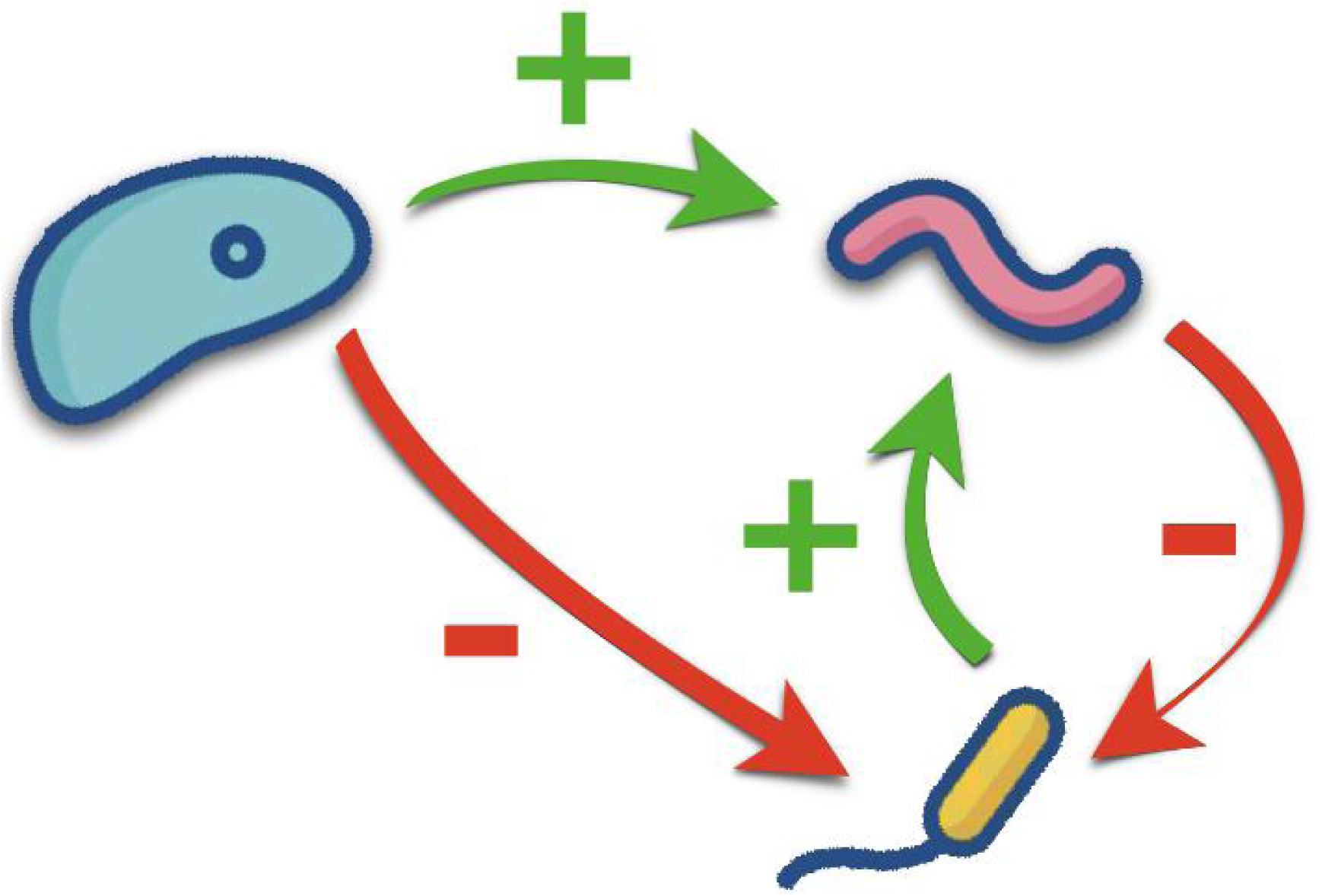
Schematic depiction of a microbial interaction network.

### Previous Work

Observe how such a framework has the flexibility to model all possible kinds of ecological interactions, not just trophic interactions^1^ where the growth of one organism necessarily comes at the expense of another organism. Previous work (López et al.,2019) has discussed comparing such trophic interaction networks^2^. In such a network signed edge weights are unnecessary because an edge from *s*_1_ to *s*_2_ indicates *s*_1_ promotes the growth of *s*_2_ and *s*_2_ inhibits the growth of *s*_1_ (necessarily because *s*_2_ consumes *s*_1_), while swapping the direction of the edge to be from *s*_2_ to *s*_1_ indicates the opposite relationship, that *s*_2_ promotes the growth of *s*_1_ and *s*_1_ inhibits the growth of *s*_2_ (due to *s*_1_ consuming *s*_2_). In the signed network framework, the former corresponds to 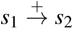 and 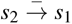, while the latter corresponds to 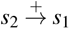 and 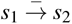. Note also that the signed network framework does not require the assumption that the cause of this asymmetric interaction is one organism consuming the other. For example, humans promote the growth of rats because human waste serves as a food source for rats, while rats can inhibit the growth of humans by serving as a vector for human diseases. The signed network framework also allows modelling mutualistic/mutually supporting interactions (pairs of interactions of the form 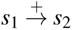 and 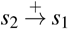), competitive/mutually antagonistic interactions (pairs of interactions of the form 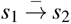 and 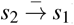), commensalistic interactions (pairs of interactions of the form 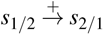 and 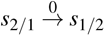), and amensalistic interactions (pairs of interactions of the form 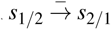 and 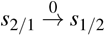). None of these other kinds of interactions can be described by the unsigned trophic network framework.

Previous work has looked at e.g. comparing “local” neighborhoods of nodes in (unweighted)^3^ signed networks (Zhu et al.,2017) or diffusion kernels in signed networks (Qi et al.,2008). However, overall the existing literature on signed networks seems underdeveloped. This is surprising because such networks could presumably model an extremely wide range of phenomena, not just ecological interaction networks. For example, the “gene interaction networks” defined implicitly in (Mani et al.,2008) could be considered (undirected) networks with mixed-sign edge weights if the edge weights were taken to be the differences (*ε* = *W*_*xy*_ − *E*(*W*_*xy*_) in the notation of the paper) between the observed values of double mutant fitnesses and those expected under a null model of no interactions, with “positive” (*ε >* 0) edges corresponding to “alleviating” interactions (*W*_*xy*_ *> E*(*W*_*xy*_)), and “negative” (*ε <* 0) edges corresponding to “synergistic” interactions (*W*_*xy*_ *< E*(*W*_*xy*_)). More generally, any dynamical system with a “multivariate state space” that has “incomplete connectivity” could potentially be summarized using a network with mixed-sign edge weights. In particular, it seems that no previous published work discusses what is called, in the framework of (Tantardini et al.,2019), the “known node correspondence” comparison problem for networks with mixed-sign edge weights. That is surprising because the problem seems like an obvious question to ask about a class of objects with extremely broad potential applicability. If prior literature exists then most likely I was unable to find it due to it using different terminology.

### Definition of Networks with Mixed-Sign Edge Weights

A “**network with mixed-sign edge weights**” *𝒢* with *S ∈* ℕ nodes can be identified with

(i) a “**node set**” 𝒩 _*𝒢*_ (which we can always assume equals [*S*] := {1, …, *S*} without loss of generality, cf. the discussion below),

(ii) an “**edge set**” ε _*𝒢*_ ⊆ 𝒩_*𝒢*_ × 𝒩_*𝒢*_ such that (*s*_1_, *s*_2_) ∈ *ε* _*𝒢*_ if and only if there is an edge directed from node *s*_1_ to node *s*_2_, and

(iii) an “**edge weight function**” *A*_*𝒢*_ : *𝒩*_*𝒢*_ *×𝒩*_*𝒢*_ *→* ℝ such that

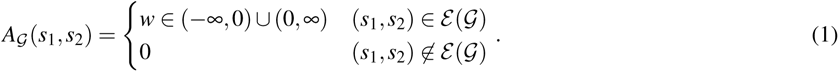

For each edge (*s*_1_, *s*_2_) ∈ *ε*_*𝒢*_, the value *A*_*𝒢*_ (*s*_1_, *s*_2_) is the “**weight**” of the edge, the value sign(*A*_*G*_(*s*_1_, *s*_2_)) = *±*1 is the “**sign**” of the edge, and the value |*A*_*𝒢*_ (*s*_1_, *s*_2_)| is the “**magnitude**” of the edge. Given an explicit identification between *𝒩*_*𝒢*_ and [*S*] (cf. below), there always exists a unique *S × S* matrix **A**_*𝒢*_, which is called the “**adjacency matrix**” of the network *𝒢*, that corresponds to the edge weight function *A*_*𝒢*_.

For every pair *s*_1_, *s*_2_ ∈ *ε*_*𝒢*_ such that *s*_1_ ≠ *s*_2_, the definition allows for at most one edge directed from *s*_1_ to *s*_2_, and for at most one edge directed from *s*_2_ to *s*_1_. Likewise, for every *s* ∈ *ε*_*𝒢*_, the definition allows for at most one edge (“self-loop”) directed from *s* to *s*. I.e. “multigraphs” are not considered.

### Monotonicity Principle

Because literature already exists on comparing unsigned networks, a practical paradigm for comparing signed networks focuses on how to extend comparison methods for unsigned networks to also apply to signed networks. Herein we will focus only on extending comparison methods for unsigned networks that satisfy a mild “well-behavedness” criterion, termed the “monotonicity principle”. Cf. figure 2.

**Figure 2.**
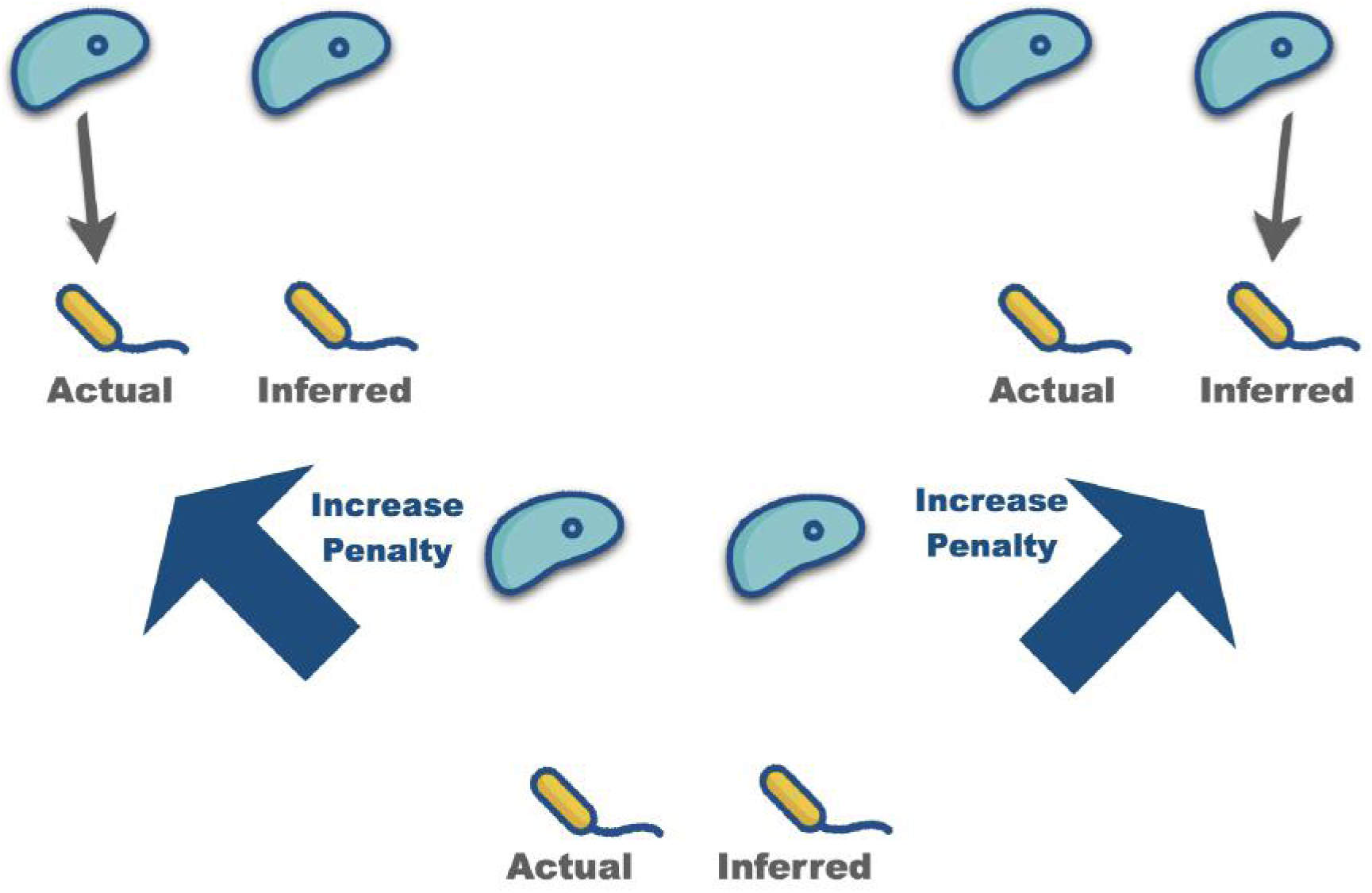
Schematic illustration of the monotonicity principle for well-behaved comparison functions of unsigned networks.

At a high level, the idea of the monotonicity principle is that “reasonable” comparison methods for unsigned networks should penalize existence errors no less than they penalize “true misses”, i.e. situations where an edge is missing in both networks. One can also ask for a “continuous” version of the monotonicity principle whereby the size of the penalty for an existence error must never decrease as the magnitude of the weight of the unmatched edge increases.

Because the monotonicity principle is a very intuitive property that one would most often tacitly assume to be true by default, we can consider comparison methods for unsigned networks that satisfy the monotonicity principle to be “reasonable”. Therefore, to implement our paradigm of extending such comparison methods to also apply to signed networks, we need to be able to check whether any proposed extension satisfies the analogous intuitive property for signed networks. This of course first requires us to identify what the analogous intuitive property for signed networks is, which we do below.

### The Double Penalization Principle

Confusing two (non-zero) edges with different signs is the only new kind of error that can occur when comparing signed networks that possesses no analogue when comparing unsigned networks. All other kinds of errors have analogues in the comparison of unsigned networks and hence can be treated using a “reasonable” comparison method for unsigned networks. Thus our main challenge, when trying to identify “reasonable” ways to extend comparison methods intended for unsigned networks to also apply to signed networks, is ensuring that errors confusing edges with different signs are treated in a “reasonable” way.

When interpreting positive and negative edge weights as corresponding to distinct (“equal but opposite”) phenomena, a mistake confusing edges with different signs amounts to the composite of two separate mistakes: both (i) inferring a phenomenon which does not exist, and (ii) failing to infer a phenomenon which does exist. Being worse than either of those two separate mistakes when considered individually, such a composite mistake should therefore receive a penalty that is no smaller than either of the penalties given to the two separate mistakes. This observation leads directly to the double penalization principle.

### Double penalization principle

*When the sign of an edge differs between two networks, the resulting penalty should be larger than the maximum of the two penalties that would occur if the edge was missing in either network*.

In other words, the penalty should equal the maximum penalty that could occur if the edge was missing from either graph, plus an additional (“second”) penalty. Both mistakes, of inferring the wrong sign and of failing to infer the correct sign, should be penalized. The “composite” mistake should therefore be “doubly penalized”. Cf. figure 3. The double penalization principle is equivalent to requiring that the monotonicity principle is simultaneously satisfied for both the positive parts and for the negative parts of the signed networks being compared. (Cf. supplemental section S2.4 for definitions of “positive part” and “negative part” of a signed network. Both are unsigned networks, with the positive part having weights equal to the magnitudes of edges with positive sign, and analogously for the negative part.)

**Figure 3.**
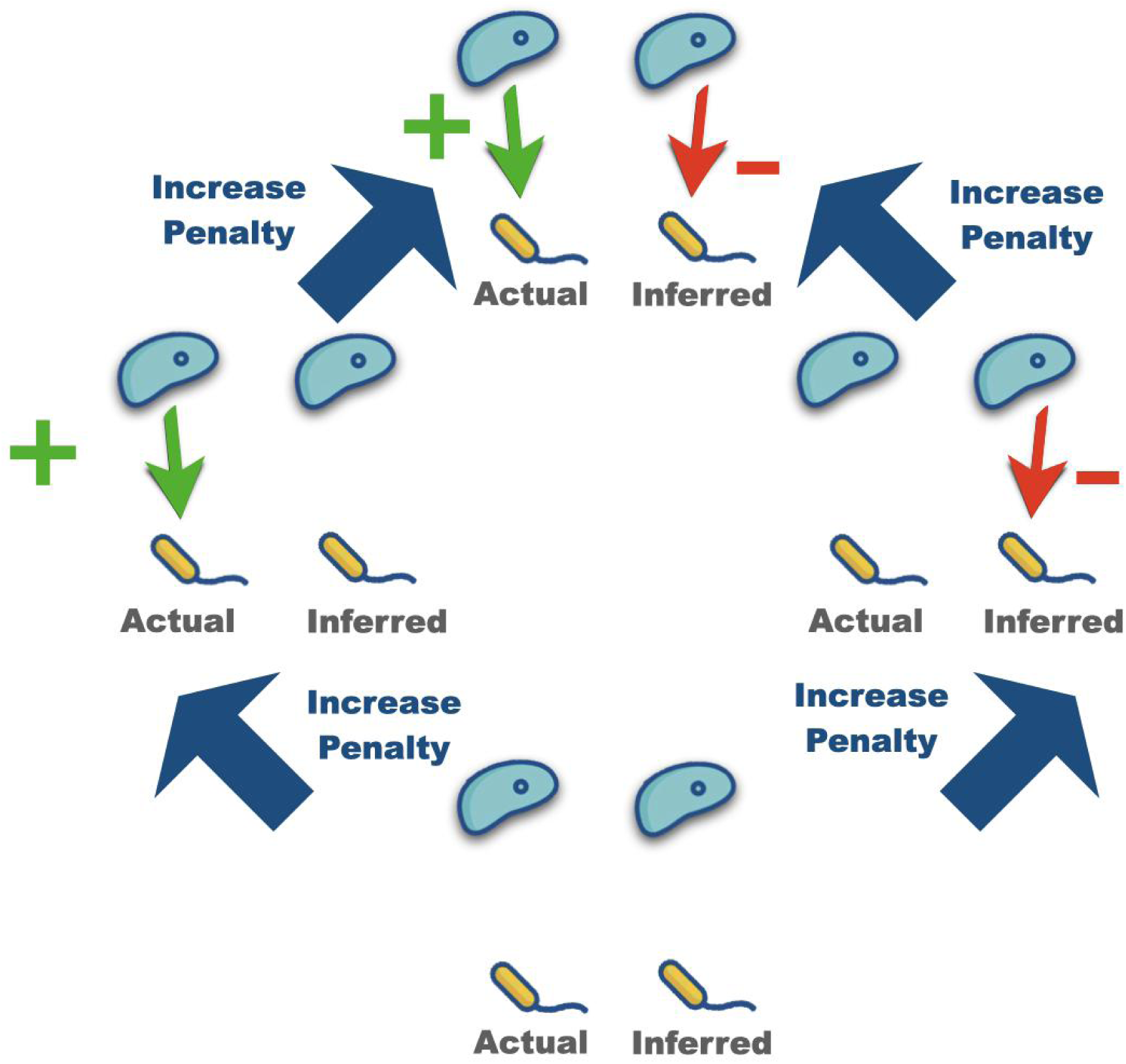
Double Penalization Principle

Note that the double penalization principle excludes naive comparison methods for signed networks that “project” the signed networks being compared onto a space of unsigned networks and then apply a comparison method for unsigned network to the “projections”. This means that the most straightforward idea for extending comparison methods for unsigned networks to also apply to signed networks fails to create comparison methods for signed networks that either are “well-behaved” or “behave intuitively”. This is true even when the original comparison method for unsigned networks is “well-behaved” and “behaves intuitively”, in the sense that it satisfies the monotonicity principle.

### Aspects of Networks to Compare

There are many aspects of networks we might seek to capture in our comparisons.

“Numerical” comparisons describe the correspondence of the numerical values of the edge weights. This could refer to only their magnitudes, or to their signed values. Relative error using a given norm (e.g. entrywise *L*_1_) is an example. Cf. supplementary section S3.1 for details.

There are also many ways we might seek to compare networks more “qualitatively”.

We might seek comparisons that describe the relative ordering of the edge weights (which are largest, which are smallest), either in magnitude, signed value, or both. (Sparsity-adjusted) Spearman correlation is an example. Cf. supplementary section S3.2 for details.

We might also seek “qualitative” comparisons that describe features of network topology. For example, (unweighted) Jaccard similarity describes the presence/absence of edges. Cf. supplementary section S3.3 for details. Similarly, (unweighted) DeltaCon distance (Koutra et al.,2016) describes the correspondences of paths. Cf. supplementary section S3.4 for details. One could also define comparisons to describe the correspondences of other subgraph motifs besides paths, although that is outside the scope of this paper.

We can also seek to capture both “numerical” and “qualitative” aspects of networks in a single comparison. Usually this is done by considering the correspondence of given qualitative features more or less important depending upon the weights of the edges in those features. Examples include weighted Jaccard similarity and weighted DeltaCon distance.

## METHODS

1, 000 Erdos-Renyi random 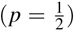 networks were generated, with edge weights determined by a hierarchical Uniform-Beta distribution (see supplementary section S4.1 for details). For each random network, whichever sign (+ or −) corresponded to more edges was designated “predominant”, the other “non-predominant” (see supplementary section S4.2 for details).

Adversarial attacks were then applied to each of the networks. The “magnitude swap attack” affected the magnitudes of the networks’ edges but not their signs, while the “sign flip attack” affected the signs if the networks’ edges but not their magnitudes. A third adversarial attack, the “shift attack”, affected both magnitudes and signs. See supplementary section S4.3 for details.

The values of the mixed-sign network comparison methods between the original networks and their attacked versions were compared with the corresponding values for variants which discard information or which do not obey the double penalization principle. “Magnitudes only” variants consider magnitude information but discard all sign information. “Signs only” or “mixed-sign unweighted” variants consider sign information but discard all magnitude information. “Presence/absence only” variants discard both magnitude and sign information. “Mixed-sign” or “mixed-sign weighted” variants (including the default relative error) consider both magnitude and sign information. See supplementary section S4.4 for details.

Side-by-side violin plots of the results were generated using Matplotlib (Hunter,2007) version 3.4.1 and Seaborn (Waskom,2021) version 0.11.1. Complete implementation details can be found in the code atthe relevant GitLab repository. See https://gitlab.com/krinsman/mixed-sign-networks.

## RESULTS

For all dissimilarity measures, variants that discard sign information (“magnitudes only” or “presence/absence only”) were completely oblivious to the sign flip attack, which affects signs but leaves magnitudes unchanged. Similarly, variants that discard magnitude information (“signs only”, “mixed-sign unweighted”, or “presence/absence only”) were completely oblivious to the magnitude swap, which affects magnitudes but leaves signs unchanged. The “presence/absence only” variants that discard sign and magnitude information were even oblivious to the shift attack, which affects both signs and magnitudes. Meanwhile, the “mixed-sign” or “mixed-sign weighted” variants were sensitive to all adversarial attacks.

### Relative Error

The versions of relative error used here takes values between 0 and 2, with smaller values corresponding to more similar networks. Cf. supplementary sections S3.1.4 and S3.1.2.

For all three adversarial attacks (figures 4, 5, and S5), we see that the relative error, which by default considers both magnitude and sign information, is always sensitive to the major changes that occur.

**Figure 4.**
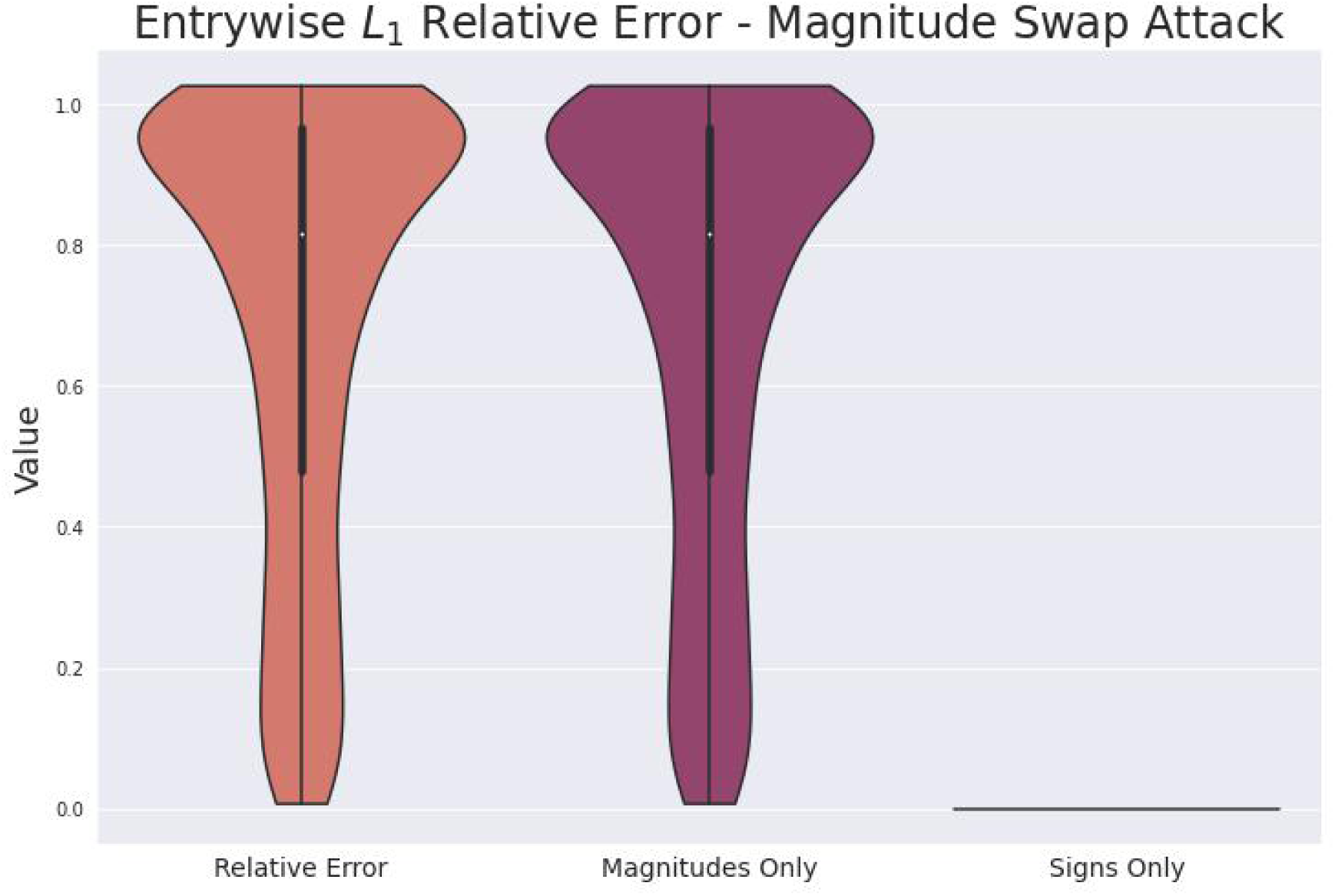

### Spearman Correlation

Spearman correlation takes values between −1 and 1, with higher values corresponding to more similar networks.

For all three adversarial attacks (figures 6, 7, and 8) we see that the mixed-sign Spearman correlation is always sensitive to the major changes that occur. At the same time, it is also often able to infer some similarity between the original networks and their attacked counterparts. In many cases this is more reasonable behavior than failing to infer any similarity, or inferring complete “anti-similarity”. The adversarial attacks were designed so that, depending on the structure of the original network (e.g. the percentage of edges belonging to the non-predominant sign), the attacked network could have substantial commonalities with the original network in addition to the blatant differences. Cf. again section S4.3.

The raw Spearman correlation responds similarly to the mixed-sign Spearman correlation for the magnitude swap attack (figure 6), although the mixed-sign Spearman correlation has a wider range of values. In contrast, the raw Spearman correlation is unable to never able to notice any similarity to the underlying network for the sign flip attack (figure 7), even for networks for which the edges with the non-predominant sign are a substantial minority, a problem the mixed-sign Spearman correlation does not have. Most problematically, the raw Spearman correlation is completely oblivious to the shift attack (figure 8). Although raw Spearman correlation arguably does consider both magnitude and sign information, it only does so indirectly. Raw Spearman correlation does not distinguish in any way between negative and positive edges as distinct phenomena, and thus usually does not satisfy the double penalization principle. This failure to distinguish negative and positive edges enables its obliviousness to the shift attack.

For those attacks to which they are not completely oblivious, the magnitudes only and signs only variants are arguably too sensitive, being unable to ever detect any similarities regardless of the particular network structure. This is most evident in the behavior of the magnitudes only variant in response to the the magnitude swap attack (figure6). Similarly for the shift attack (figure 8) the magnitudes only variant only ever reports “anti-similarity”. The signs only variant responds the same way to the sign flip (figure 7) and shift attacks, because for both adversarial attacks the sign vector of the attacked network is constant, cf. the note from section S3.2. For both adversarial attacks this makes the signs only variant is unable to report any useful information that would reflect the original network’s particular structure.

### Jaccard Similarity

Jaccard similarity takes values between 0 and 1, with larger values corresponding to more similar networks.

The wider flare at the bottom of the distribution of the mixed-sign weighted Jaccard similarities suggests that it is even more sensitive to the sign flip attack (figure 9) than the mixed-sign unweighted Jaccard similarity, even though only signs are affected. The magnitudes only variant is surprisingly insensitive to the shift attack (figure 10); even the mixed-sign unweighted Jaccard similarity is more sensitive. Both patterns suggest more importance for sign information than for magnitude information.

### DeltaCon Distance

DeltaCon distance takes non-negative values, with smaller values corresponding to more similar networks.

Only the mixed-sign weighted DeltaCon distance is sensitive to all three adversarial attacks (figures 11, 12, 13). The mixed-sign weighted DeltaCon distance is always more sensitive than the magnitudes only variant. This is true even for the magnitude swap attack (figure 11), even though that adversarial attack only affects magnitudes. This may result from DeltaCon distance considering “higher-order connectivities”, because the analogous phenomenon does not occur for entrywise *L*_1_ relative error (figure 4) nor for Jaccard similarity (figures 12). Cf. the discussion in supplementary section S5.4.3.

The mixed-sign weighted DeltaCon distance appears slightly less sensitive than the mixed-sign unweighted DeltaCon distance for the sign flip attack (figure 12) and the shift attack (figure13), but that may be an artifact (cf. supplementary section S5.4.1). The mixed-sign unweighted DeltaCon distance is definitely more sensitive to the shift attack (figure 13) than the magnitudes only variant.

## DISCUSSION

### Summary of Results

The results show that attempts to use methods for comparing networks with unsigned edges can lead to useless or even actively misleading results when applied to networks with mixed-sign edge weights. The results also show how methods satisfying the double penalization principle can avoid such results.

### Consequences of Discarding Information

The “whole” of simultaneously considering both the sign and magnitude information in a network with mixed-sign edge weights appears to be “greater than the sum of the parts” of using either the sign information or the magnitude information alone. One way to see this is by noting how the behaviors of variants considering both types of information are not “intermediate” between the behaviors of their respective signs only and magnitudes only variants. This is in contrast to how the behaviors of variants considering both types of information are often intermediate (in a sense that can be made precise, cf. supplementary sections S3.1.3, S3.2.6, S3.3.8, and S3.3.9) between the behaviors on the predominant sign and non-predominant sign edges, cf. supplementary sections S5.1.2, S5.2.1, and S5.3.2.

Several examples show how variants which consider both sign and magnitude information can perform better even when seemingly only one type of information should be relevant. In response to the sign flip attack, the mixed-sign variants of relative error and Jaccard similarity demonstrate wider flares at the extremes of their distributions indicating dissimilarity than the respective corresponding signs only variants. Cf. figure 5 for relative error and figure 9 for Jaccard similarity. These mixed-sign variants appear to be more sensitive to the sign flip attack than the respective corresponding signs only variants, despite that the sign flip attack only affects signs (and leaves magnitudes unaffected). Similarly, the mixed-sign weighted DeltaCon distance tends towards larger values than its corresponding magnitudes only variant in response to the magnitude swap attack, cf. figure 11. The mixed-sign weighted DeltaCon distance appears to be more sensitive to the magnitude swap attack than its corresponding magnitudes only variant, despite that the magnitude swap attack only affects magnitudes (and leaves signs unaffected).

**Figure 5.**
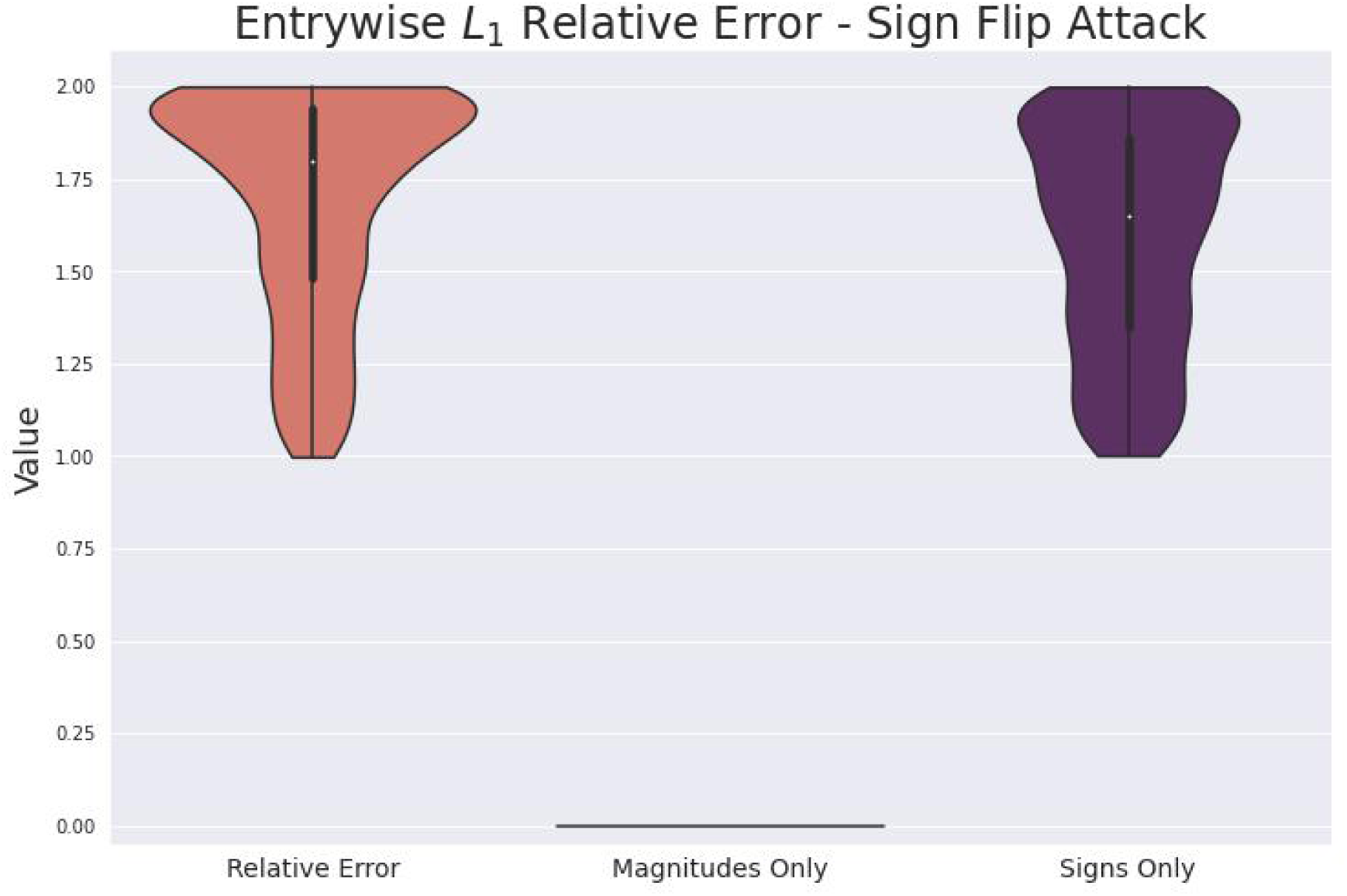

**Figure 6.**
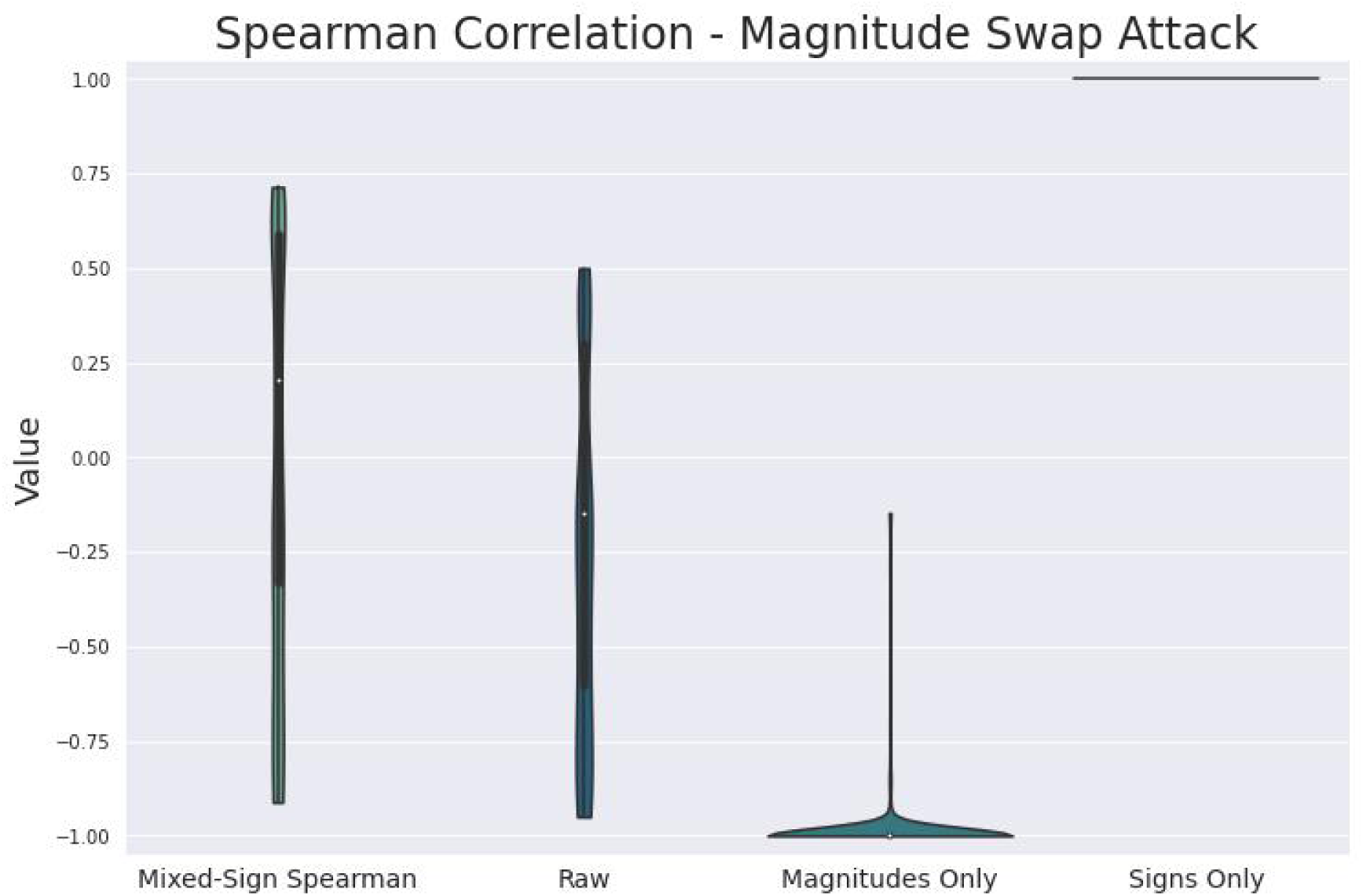

**Figure 7.**
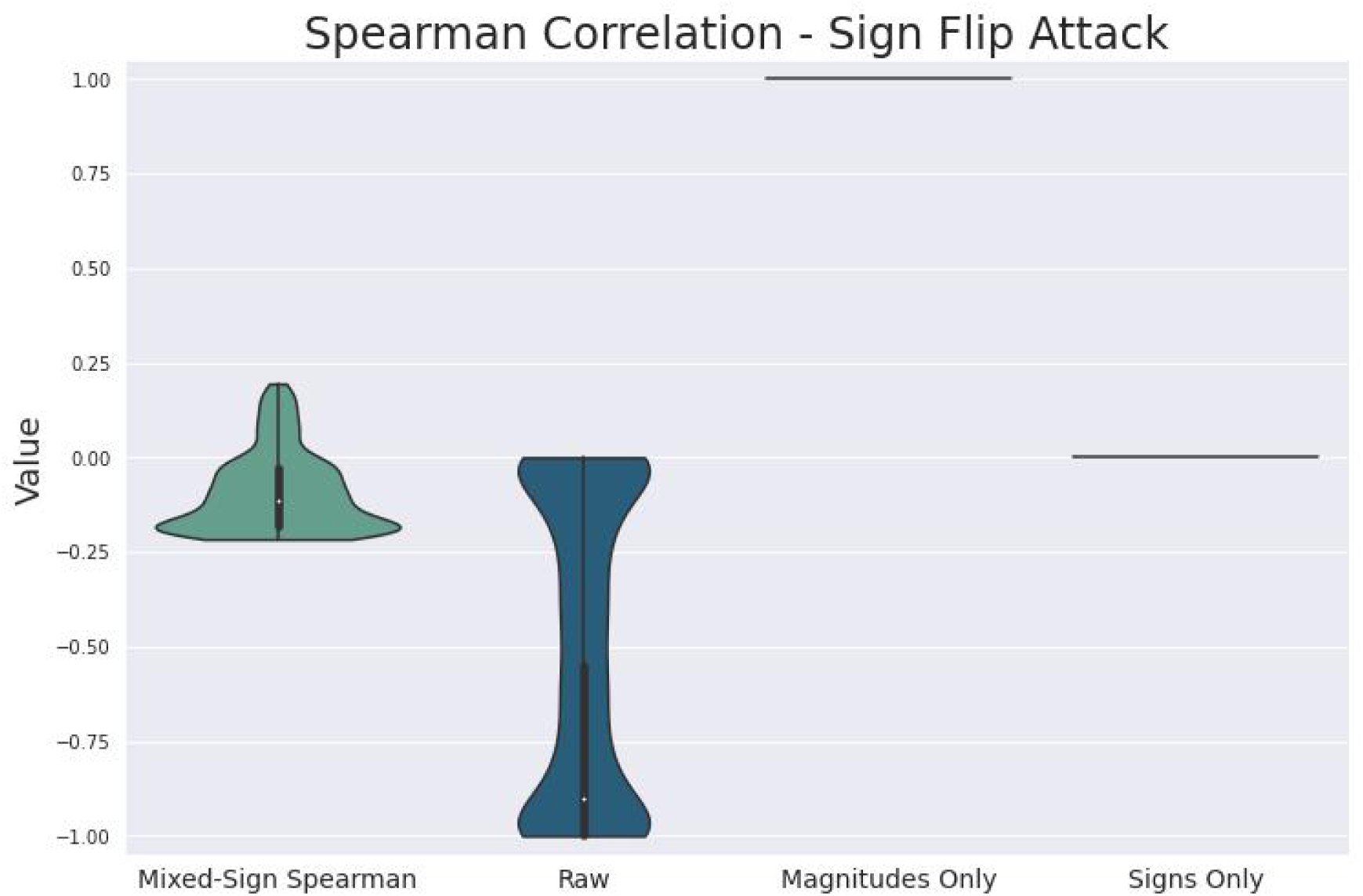

**Figure 8.**
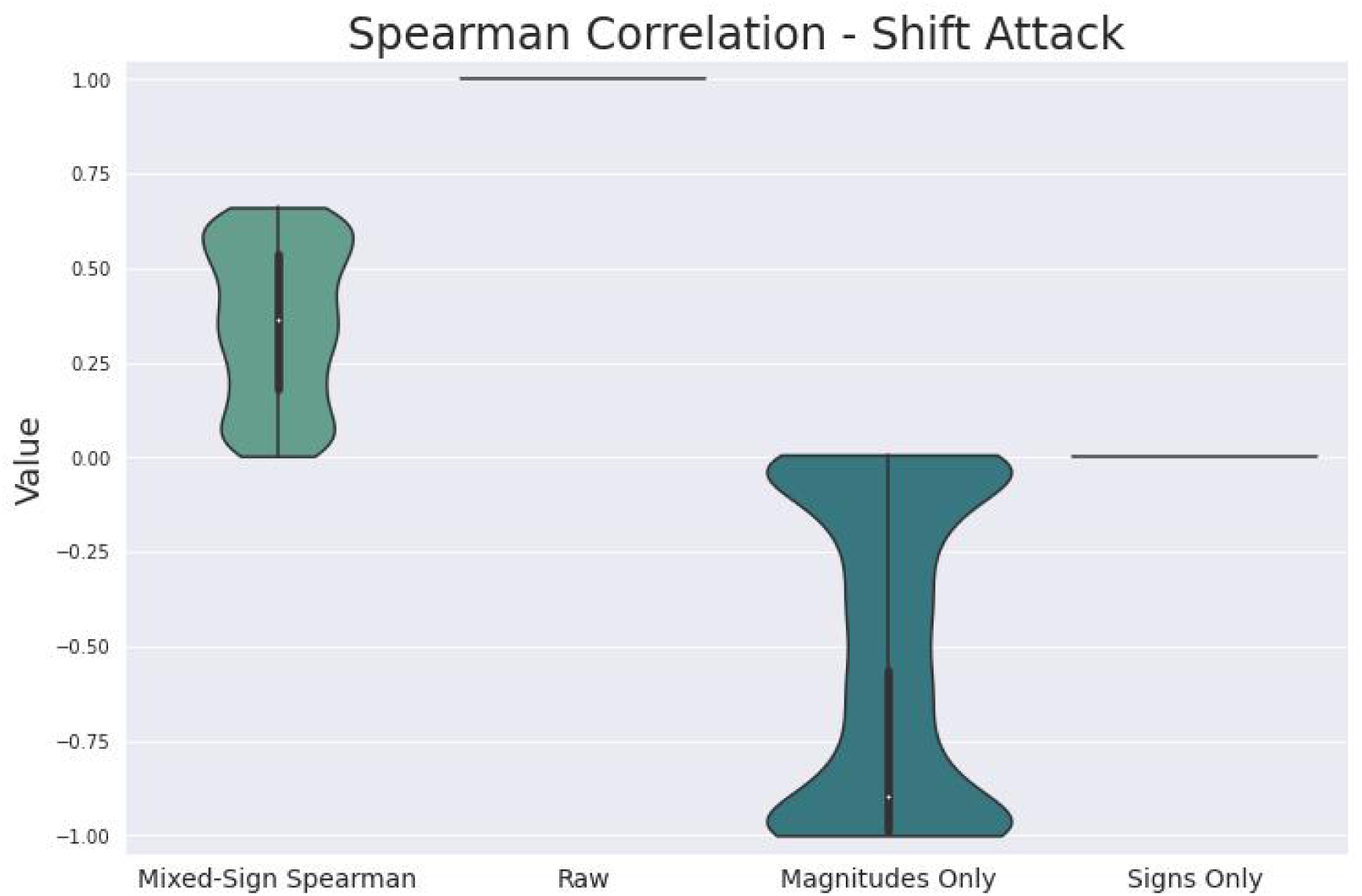

**Figure 9.**
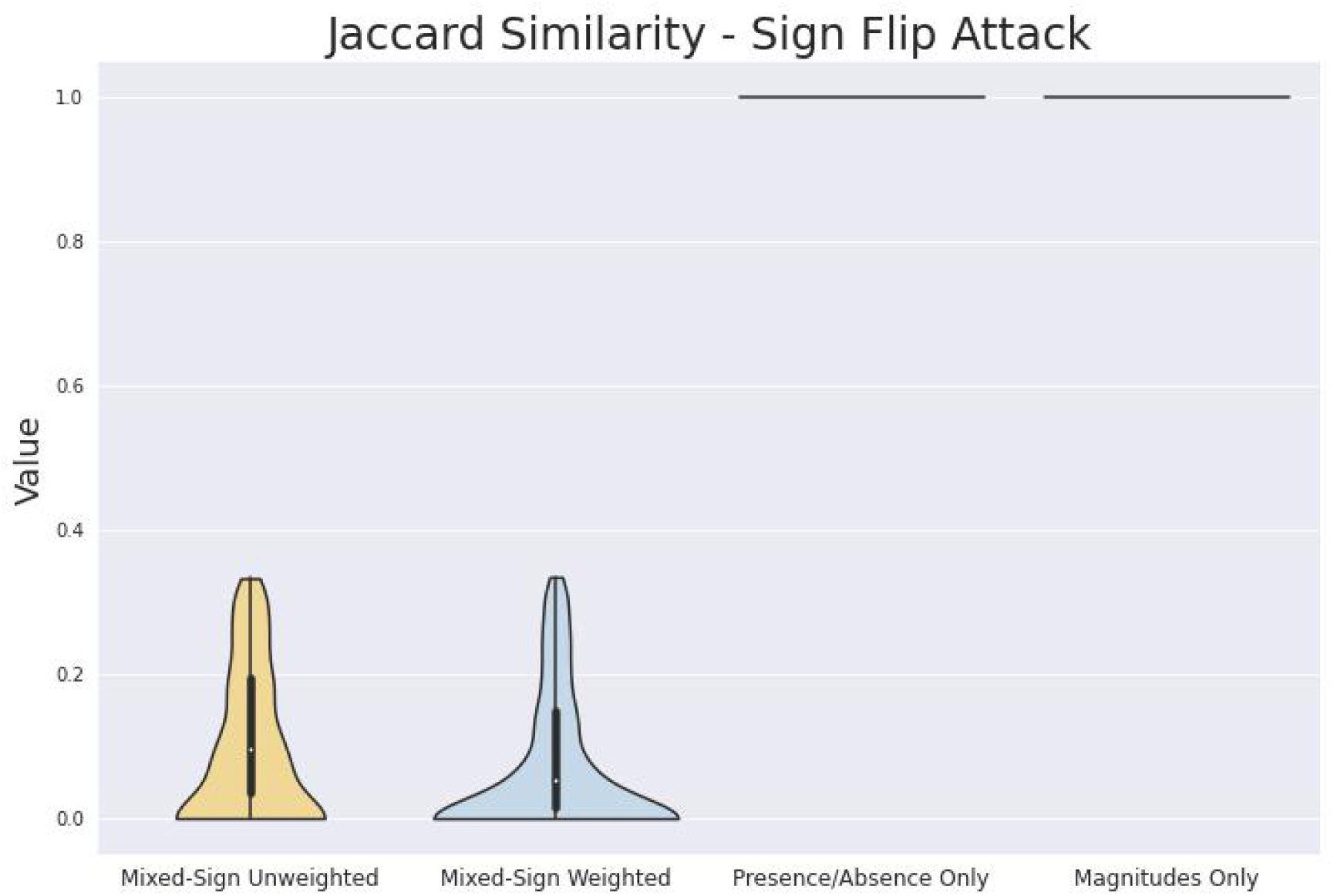

Blindly applying standard methods thar are intended for less structured networks to networks with mixed-sign edge weights can lead to drastically misleading conclusions. Doing so requires discarding information about the networks with mixed-sign edge weights that are being compared. In particular, the “presence/absence only” variants that might result from researchers trying to apply standard methods intended for unsigned and unweighted networks were completely oblivious to all of the adversarial attacks. The observed behavior suggests that even combining the results of discarding separate pieces of information, a “divide and conquer” approach, would be inadequate to avoid this. Instead, the better behavior of variants that consider both sign and magnitude information appears to be an emergent phenomenon of considering both kinds of information simultaneously. Thus we need to acknowledge and confront the problem by respecting the richer structure of networks with mixed-sign edge weights.

### Effectiveness of the Double Penalization Principle

For the shift attack, where both magnitudes and signs are changed, we might have expected signs only and magnitudes only variants to be equally sensitive. Both variants discard some but not all information. However, for those comparisons where the signs only variant satisfies the double penalization principle (relative error in figure S5, Jaccard similarity in figure 10, DeltaCon distance in figure 13), the signs only variant is more sensitive than the magnitudes only variant. (However this may also be largely an artifact of the magnitudes being bounded to be no greater than 1, while the “magnitudes” of the signed skeletons, on which the signs only variants act, are 1, and thus no smaller. Cf. supplementary section S5.4.1.)

**Figure 10.**
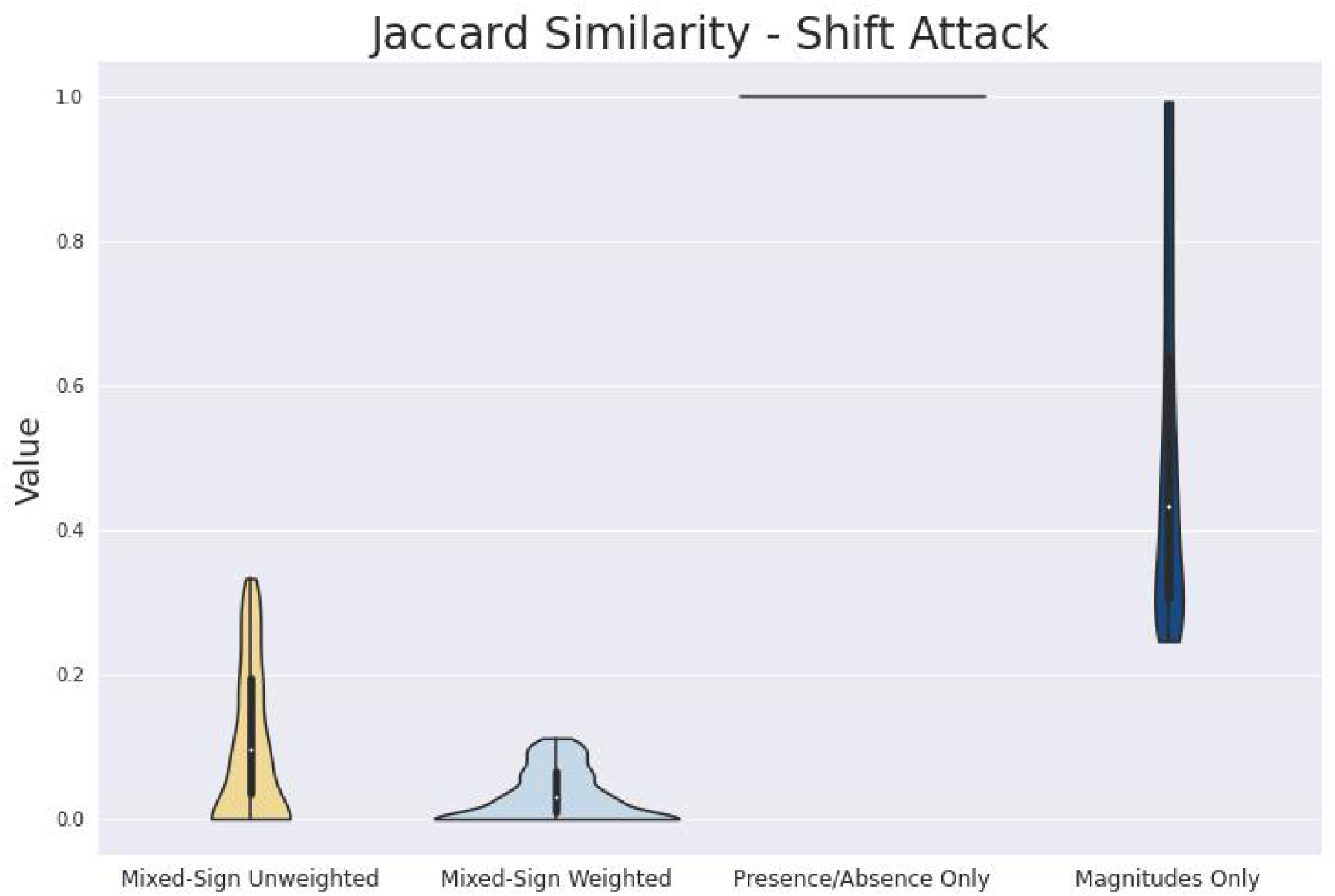

Because satisfying the double penalization principle requires sensitivity to signs, but not to magnitudes, this seems to suggest that sensitivity to signs is more important than sensitivity to magnitudes, *as long as the double penalization principle is satisfied*. Blind application of default methods for unsigned networks will usually be completely insensitive to signs, much less obey the double penalization principle.

Double penalization may explain why the the mixed-sign DeltaCon variants, weighted or unweighted, are generally more sensitive to all three adversarial attacks than the presence/absence only or magnitudes only variants. (Cf. again figures 11, 12, and 13.) For example, even though signs are unchanged, the mixed-sign weighted DeltaCon distance is more sensitive to the magnitude swap attack (figure 11) than the magnitudes only variant. This may imply that the double penalization principle is being applied to the corresponding similarity matrices, cf. equation (S45), whose signs may have changed.

**Figure 11.**
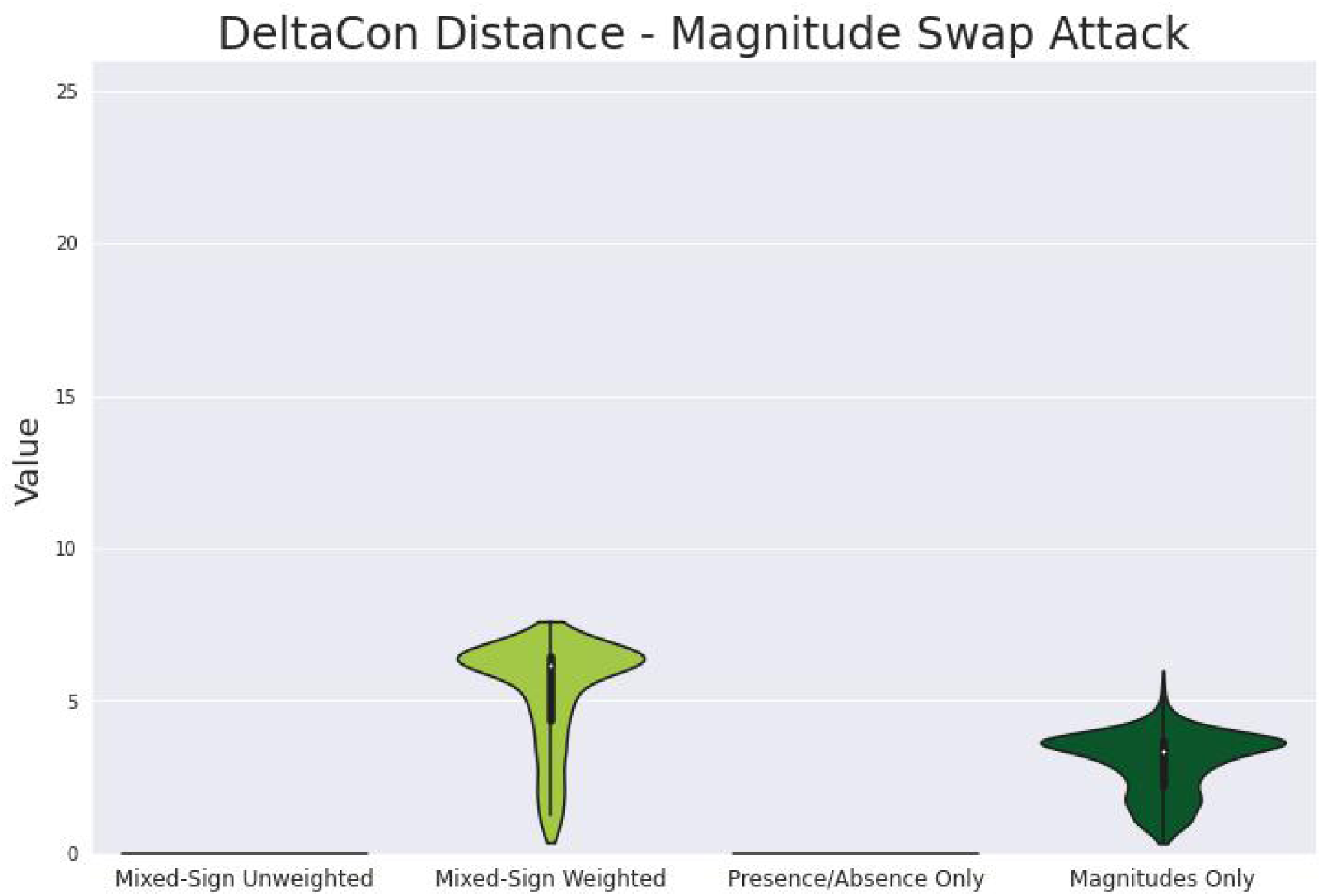

**Figure 12.**
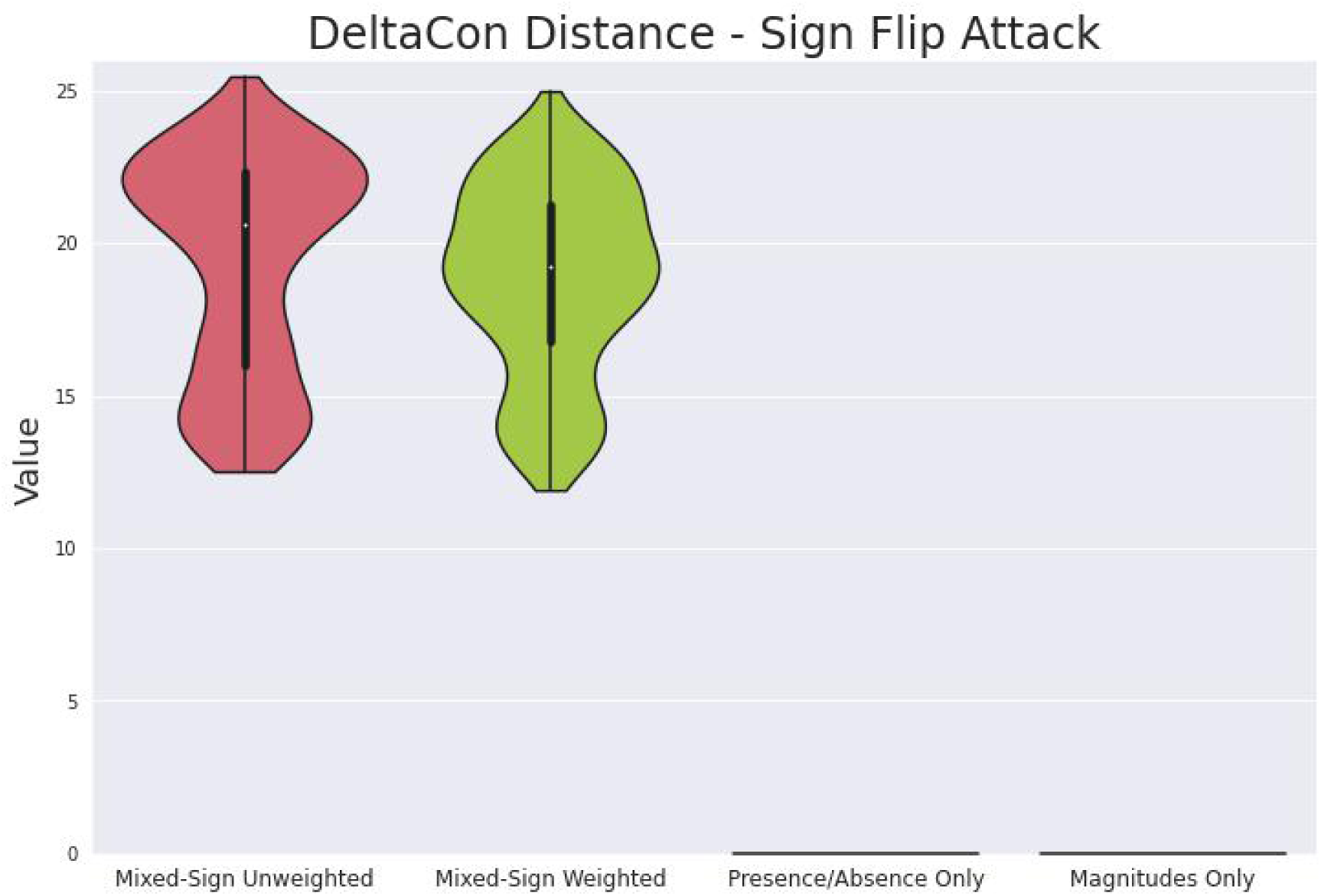

**Figure 13.**
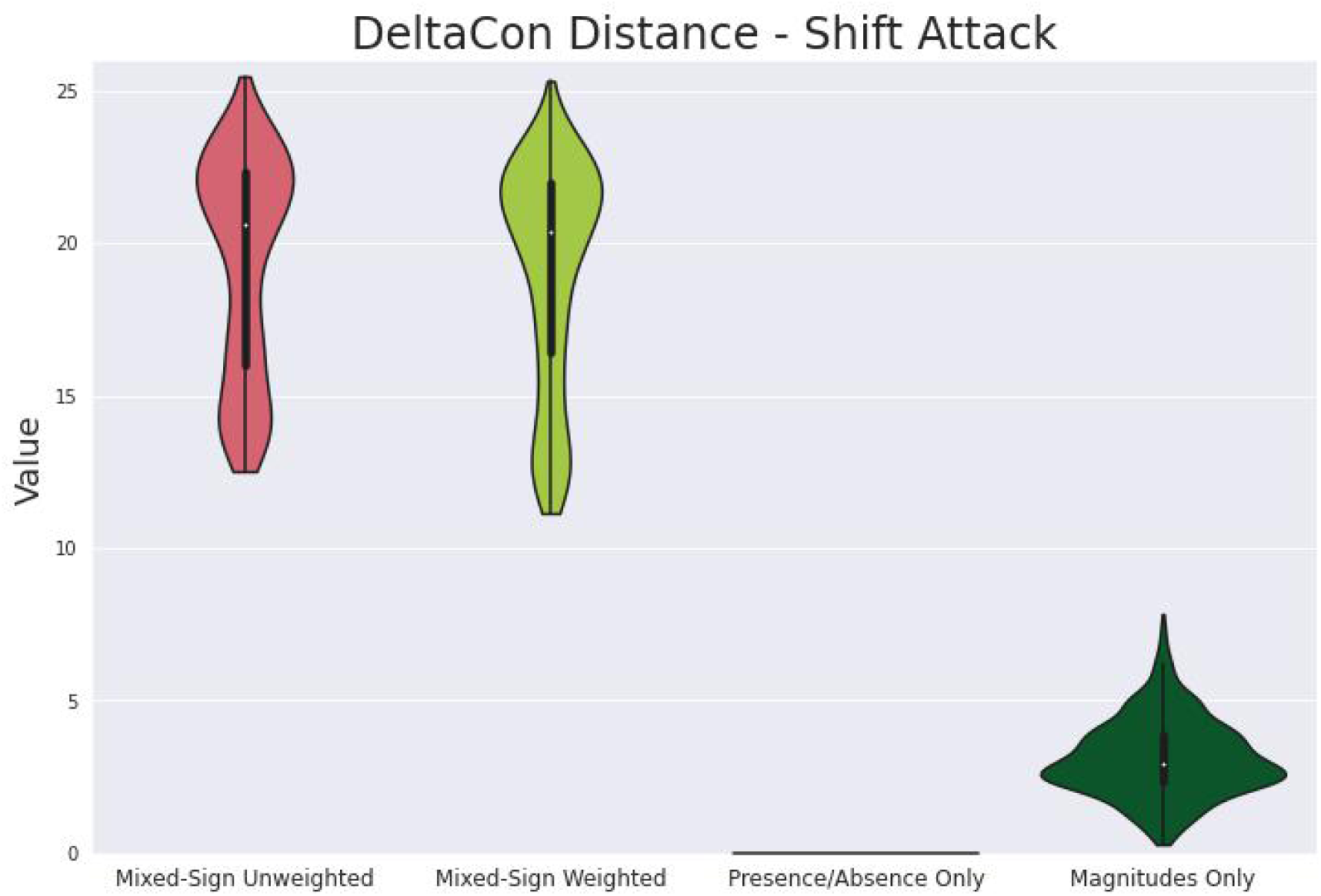

### Limitations

These results only considered a limited number of adversarial attacks. The goal of this initial work was to consider adversarial attacks which did not affect the overall connectivity of the network and which affected limited aspects of the structure in obvious ways. One can think of many more changes to structure of mixed-sign networks for which we would want (dis)similarity measures to behave well. Future work can investigate whether good responses are also observed for other kinds of adversarial attacks.

The goal was to consider a large number of networks with varying structure, such that altogether they might be enough to represent all “typical” networks. It could be argued that the chosen distribution of random networks is insufficient to accomplish that goal. See section S6.2.2 for details. For example, it would be interesting to use more edges with magnitudes very different from 1 (cf. e.g. section S5.4.1). Future work might investigate whether these results can be replicated using different kinds of networks.

This work provides no systematic procedure for extending arbitrary (dis)similarity measures to the case of networks with mixed-sign edge weights. The double penalization principle is merely a “design specification”. It seems unlikely that such a systematic procedure could exist. Nevertheless, the absence of such a systematic procedure could possibly still be considered to limit the usefulness of this work.

## CONCLUSIONS

Herein I identified that trying to compare networks with mixed-sign edge weights using standard network comparison methods can give useless or misleading results. I showed that the extra structure of networks with mixed-sign edge weights means that they need to be compared differently than “standard” networks. I identified a unifying principle, the double penalization principle, for identifying methods that are potentially useful for comparing networks with mixed-sign edge weights. I implemented this principle for several methods and showed that one simple method (relative error) already satisfies it “out of the box”. This work allows us to directly address the difficulties inherent in making meaningful comparisons between networks with mixed-sign edge weights, rather than ignore those difficulties.

## Supporting information

Supplementary Sections

## ACKNOWLEDGMENTS

Professor Adam P. Arkin for critical suggestions, essential scientific and technical advice, important introductions, and access to funding and computational resources without which this project would have been impossible. Dr. Fangchao Song for suggesting the topic of the dissertation (Krinsman,2022) and generous consultations, feedback, and advice throughout. Professor Mark van der Laan for critical suggestions, essential scientific and technical advice, and invaluable recommendations of strategies and resources for scientific writing. Dr. Lauren M. Lui for invaluable recommendations of strategies and resources for scientific writing.

