## Supplementary Sections for "Extending Comparison Methods for Unsigned Networks to Signed Networks"

**William Krinsman**[1]

[1]Graduate Group in Biostatistics; University of California, Berkeley
[1]Division of Computing, Data Science, and Society; University of California, Berkeley
[1]Center for Computational Biology; University of California, Berkeley
[1]BioSciences Area; Lawrence Berkeley National Laboratory

Section S1 explains why sensibly quantifying the (dis)similarity of two networks with mixed-sign edge weights helps to characterize microbial interactions, clarifies the problem, and identifies principles to judge whether a method sensibly quantifies the (dis)similarity of two networks with mixed-sign edge weights. Section S2 provides technical definitions which facilitate the analysis in the following sections. Section S3 proposes several examples of methods that satisfy the principles described in section S1. Section S4 explains details of the experiments shown in the main text. Section S5 provides additional empirical demonstrations of how methods that satisfy these principles behave compared to methods that do not satisfy these principles and also shows how they combine information from negative and positive edges. Finally section S6 interprets the results and explains what still requires further attention.

#### Supplementary Sections

**S1 Introduction** **3**

**Additional References** **43**

### S1 INTRODUCTION

Section S1.1 explains the motivation for considering this problem. Section S1.2 clarifies the type of problem considered herein. Section S1.3 provides a criterion we might like comparison methods to satisfy. Section S1.4 provides an additional criterion which is also relevant for unsigned networks.

#### S1.1 Statistical Motivation

We want to characterize microbial interactions using the data from microbial ecology experiments. This means estimating a network with mixed-sign edge weights from the data, the microbial interaction network. The positive edges of the network correspond to microbial interactions which promote the growth of the recipient. The negative edges of the network correspond to interactions which suppress the growth of the recipient. We want to estimate this network as accurately as possible.

To estimate the microbial interaction network as accurately as possible using the data from a given microbial ecology experiment, we need to choose a "best performing" estimator in a sensible way. To choose a "best performing" estimator in a sensible way, we need to be able to compare the performance of any two estimators in a sensible way. To compare the performance of any two estimators in a sensible way, we need to compare the (dis)similarity of two estimated networks with mixed-sign edge weights to a reference "ground truth" network with mixed-sign edge weights in a sensible way. Finally, to make such comparisons between networks with mixed-sign edge weights in a sensible way, we need to be able to quantify the (dis)similarity of two networks with mixed-sign edge weights in a sensible way.

#### S1.2 Known vs. Unknown Node Correspondence

There are at least two types of network comparison problem which we should distinguish. Following the framework from (Tantardini et al., 2019), we can consider

- the known node correspondence problem (”fixed nodes, variable edges”), or
- the unknown node correspondence problem (”variable everything”).

For the known node correspondence problem, the identity of the nodes is essential and fixed. The two compared networks must either have the same set of labeled nodes, or a known correspondence between their node sets must be given. The known-node-correspondence problem asks whether the two networks describe similar relationships for the fixed set of nodes.

An example of the known node correspondence problem would be to ask how much airline flight routes in Europe changed after the onset of the pandemic. This amounts to comparing two networks (one for before the pandemic and one for after) with the same labeled nodes (the airports), but for which the edges (the flight routes) between those nodes may be different.

For the unknown node correspondence problem, the identity of the nodes does not matter. The identity of the nodes may differ between the two networks. The unknown node correspondence asks essentially whether the relationships described by the two networks are "analogous".

An example of the unknown node correspondence problem would be to ask whether the structure of airports and flight routes is analogous between Europe and China. The set of nodes is obviously different, and a priori there's no reason to believe that e.g. the Shanghai airport should be identified with the London airport rather than the Paris airport or vice versa. Nevertheless one could still sensibly ask e.g. whether it's possible to identify analogous subsets of highly connected "central hub" airports in both networks.

The network comparison problem from section S1.1 clearly is known node correspondence. We want to quantitatively assess the ability of statistical estimators to recover the correct set of edges (interactions) for a given set of nodes (microbes). It makes no difference if the estimated network is "analogous" to the true network. If the ecological roles of the various microbes are misidentified by ascribing certain relationships to the incorrect pairs of microbes, the estimated network is still a terrible estimate of the truth. Only the known node correspondence problem is relevant for comparing the estimates produced by an estimator with the true value of an estimand.

Unfortunately, the fact that our network comparison problem is known node correspondence also means that most of the methods discussed in (Tantardini et al., 2019) would be inapplicable and irrelevant even if we were considering networks with all positive edges. While comparing distinct microbial ecosystems for analogous network structures is a valid scientific problem and would require extension of methods for the unknown node correspondence problem to networks with mixed-sign edge weights, it is outside of scope here and left to future work. Herein I try to modify and extend (as necessary) most of the computationally tractable methods for the known node correspondence problem discussed in (Tantardini et al., 2019) to apply also to networks with mixed-sign edge weights.

##### S1.3 Convex Combination Decomposition Property

A sufficient, *but not necessary*, condition for satisfying the double penalization principle is

- When the sign of an edge differs between two graphs, the resulting penalty equals the sum of the two penalties that would occur if the edge was set to zero (“removed”) in either graph.

An equivalent formulation of the double penalization principle is that the penalty for the edge should equal the maximum of the two penalties that occur when considering either the networks’ positive parts only or negative parts only (see section S2.4), plus an additional penalty. This leads to the following equivalent formulation of the above sufficient, *but not necessary*, criterion:

- When the sign of an edge differs between two graphs, the resulting penalty equals the sum of the penalty from comparing the positive parts of the graphs with the penalty from comparing the negative parts of the graphs.

When the penalties are additive across edges, the above criterion means the total penalty is the (weighted) sum of the penalties for the positive parts and negative parts of the graphs. Such (dis)similarity measures can be modified by suitable normalizations to produce (dis)similarity measures whose values equal a convex combination of (i) their value when applied to the positive parts of the networks only and (ii) their value when applied to the negative parts of the networks only. The resulting (dis)similarity measures are said to have the **convex combination decomposition property**. It follows from the above that the convex combination decomposition property is sufficient, *but not necessary*, to satisfy the double penalization principle.

The convex combination decomposition property is potentially desirable not only because it guarantees that the double penalization principle will be satisfied. (Dis)similarity measures with this property are easy to interpret because understanding how the networks’ positive edges vs. their negative edges contribute to the final value is straightforward.

##### S1.4 Sparsity Should Not Lead to Artifactual Similarity

This principle is applicable to general networks, not just those with mixed-sign edge weights. The fact that an edge is absent from two networks should not lead to an increase in their similarity score. In other words, “don’t reward true misses”. Otherwise, two networks which are very sparse (have a small percentage of all possible edges that *could* exist) could potentially be scored as very similar, even when the structure of the edges they *do* have is very different.

#### S2 GENERAL DEFINITIONS

This section gives precise definitions which will be used throughout the remaining sections for describing or modifying networks. Section S2.1 gives definitions and notation which will be used interchangeably to specify network structure. Section S2.2 defines a related matrix which is important for section S3.4 later. Section S2.3 defines standard ways of “forgetting” network structure. Section S2.4 defines a standard way of decomposing a network, extending well-known decompositions applicable to the functions and matrices from section S2.1 which can be used to represent the network. Finally section S2.5 defines a standard way to join two networks together to create a new (disconnected) network.

#### S2.1 Representations of Networks as Functions, Sets, and Matrices

The $(i, j)$th entry of any matrix $\mathbf{M}$ will be denoted $[\mathbf{M}]_{ij}$.

Recall that $S$ denotes the number of strains.

Any function $\varepsilon : [S] \times [S] \to \mathbb{R}$ is equivalent to specifying a (mixed-sign) weighted and directed graph $\mathcal{G}$ with $S$ nodes. The values of $\varepsilon$ are the edges weights. Given any pair $(s_1, s_2)$, $\varepsilon(s_1, s_2)$ gives the weight of the edge directed from $s_1$ to $s_2$, or equals 0 if no such edge exists. Cf. figure S1.

Any such **edge function** $\varepsilon : [S] \times [S] \to \mathbb{R}$ can also be identified with an $S \times S$ **adjacency matrix** $\mathbf{A}$, where entry $[\mathbf{A}]_{s_1 s_2}$ of the matrix equals $\varepsilon(s_1, s_2)$. Thus the $(s_1, s_2)$'th entry $[\mathbf{A}]_{s_1 s_2}$ of the adjacency matrix $\mathbf{A}$ gives the weight of the edge from $s_1$ to $s_2$, or is equal to 0 if no such edge exists.

The set of edges $\mathcal{E}$ of the graph $\mathcal{G}$ is then defined by the set

$$\mathcal{E} := \operatorname{supp}(\varepsilon) := \{(s_1, s_2) \in [S] \times [S] : \varepsilon(s_1, s_2) \neq 0\}. \tag{S1}$$

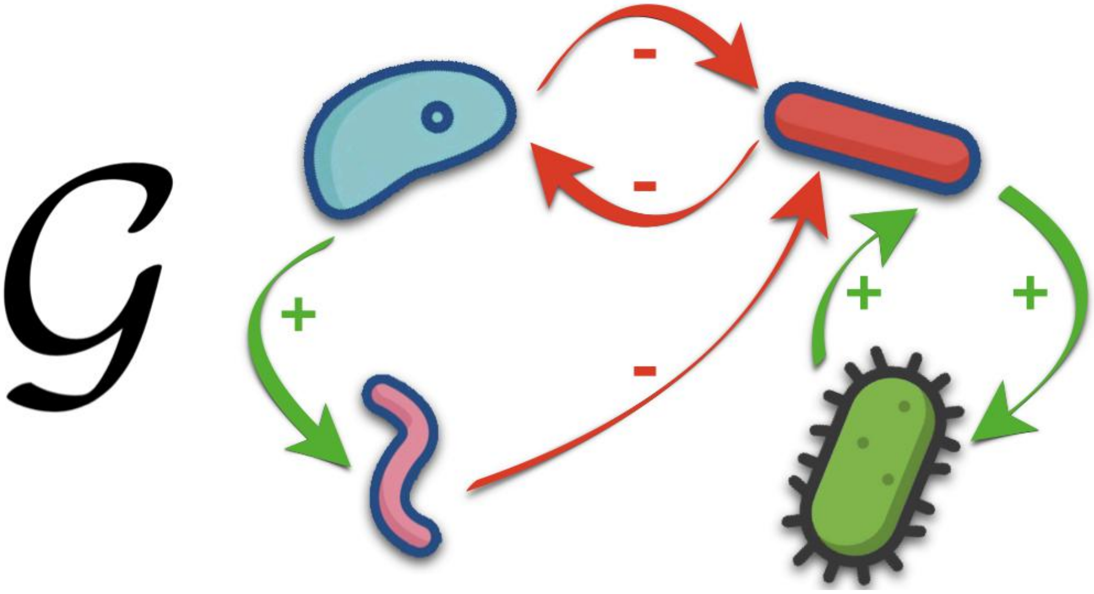

**Supplementary Figure S1.** Directed network with signed edges. Bacteria are the nodes and identified with the set $[S] = [4]$. The arrows are the edges, identified with $\operatorname{supp}(\varepsilon)$ where $\varepsilon : [4] \times [4] \to \mathbb{R}$. Green arrows are positively weighted edges, identified with the set $\{(s_1, s_2) \in [4] \times [4] : \varepsilon(s_1, s_2) > 0\}$. Red arrows are negatively weighted edges, identified with the set $\{(s_1, s_2) \in [4] \times [4] : \varepsilon(s_1, s_2) < 0\}$.

#### S2.2 The Degree Matrix

**Note:** the material in this section is only used later on in section S3.4.

The degree matrix $\mathbf{D}$ of a network $\mathcal{G}$ with adjacency matrix $\mathbf{A}$ is defined as

$$[\mathbf{D}]_{s_1 s_2} := \begin{cases} \sum_{\sigma_2 = 1}^{S} \mathbf{A}_{s_1 \sigma_2} & s_1 = s_2 \\ 0 & \text{else} \end{cases}. \tag{S2}$$

Intuitively speaking, the $s$'th entry on the diagonal of $\mathbf{D}$ encodes the "net influence" flowing out from node $s$. In the case of an unsigned and unweighted network, i.e. an adjacency matrix consisting entirely of 1's and 0's, this is quantified by the number of the number of edges originating from node $s$.

#### S2.3 Skeletons

Unsigned skeletons (section S2.3.1) preserve the least amount of structure of the original network. Signed skeletons (S2.3.2) and magnitude skeletons (S2.3.3) preserve more structure. By definition all notions of "graph skeleton" lead to networks with edge sets $\mathcal{E}$ that are the same as that of the original network.

###### *S2.3.1 (Unsigned) Skeletons*

Given a network $\mathcal{G}$ with adjacency matrix $\mathbf{A}$, the **unsigned skeleton** $\mathcal{S}(\mathcal{G})$ is defined to be the network with adjacency matrix $\mathbf{B}$ such that

$$[\mathbf{B}]_{ij} := \begin{cases} 1 & [\mathbf{A}]_{ij} \neq 0\,, \\ 0 & [\mathbf{A}]_{ij} = 0\,. \end{cases} \tag{S3}$$

The unsigned skeleton $\mathcal{S}(\mathcal{G})$ preserves only the underlying connectivity/topology of $\mathcal{G}$, while "deleting" all information about edge weights or signs.

###### *S2.3.2 Signed Skeletons*

Given a network $\mathcal{G}$ with adjacency matrix $\mathbf{A}$, the **signed skeleton** $\mathcal{S}^{\pm}(\mathcal{G})$ is defined to be the network with adjacency matrix $\mathbf{B}$ such that

$$[\mathbf{B}]_{ij} := \operatorname{sign}([\mathbf{A}]_{ij}) = \begin{cases} 1 & [\mathbf{A}]_{ij} > 0\,, \\ -1 & [\mathbf{A}]_{ij} < 0\,, \\ 0 & [\mathbf{A}]_{ij} = 0\,. \end{cases} \tag{S4}$$

In addition to the underlying connectivity of the original network $\mathcal{G}$, the signed skeleton $\mathcal{S}^{\pm}(\mathcal{G})$ also preserves the signs of the edge weights. The signed skeleton $\mathcal{S}^{\pm}(\mathcal{G})$ still "deletes" all information about edge magnitudes.

Note that for a network with non-negative edge weights, the unsigned and signed skeletons coincide. Thus the signed skeleton is only a distinct and new notion for the more general setting of networks with mixed-sign edge weights.

###### *S2.3.3 Magnitude Skeletons*

Given a network $\mathcal{G}$ with adjacency matrix $\mathbf{A}$, the **magnitude skeleton** $\mathcal{M}(\mathcal{G})$ is defined to be the network with adjacency matrix $\mathbf{B}$ such that

$$[\mathbf{B}]_{ij} := \operatorname{abs}([\mathbf{A}]_{ij}) := \left|[\mathbf{A}]_{ij}\right| = \begin{cases} [\mathbf{A}]_{ij} & [\mathbf{A}]_{ij} > 0\,, \\ -[\mathbf{A}]_{ij} & [\mathbf{A}]_{ij} < 0\,, \\ 0 & [\mathbf{A}]_{ij} = 0\,. \end{cases} \tag{S5}$$

In addition to the underlying connectivity of the original network $\mathcal{G}$, the magnitude skeleton $\mathcal{M}(\mathcal{G})$ also preserves the magnitudes of the edge weights. The magnitude skeleton $\mathcal{M}(\mathcal{G})$ still "deletes" all information about edge signs.

Note that for a network with non-negative edge weights, the magnitude skeleton coincides with the original network. Thus the magnitude skeleton is only a distinct and new notion for the more general setting of networks with mixed-sign edge weights. Relatedly, observe how $\mathcal{S}(\mathcal{G}) = \mathcal{M}(\mathcal{S}^{\pm}(\mathcal{G}))$.

##### S2.4 Positive and Negative Parts

Any adjacency matrix $\mathbf{A}$ admits the decomposition:

$$\mathbf{A} = \mathbf{A}^{+} - \mathbf{A}^{-}\,, \quad [\mathbf{A}^{+}]_{s_1 s_2} := \max\{A_{s_1 s_2}, 0\}\,, \quad [\mathbf{A}^{-}]_{s_1 s_2} := \max\{-A_{s_1 s_2}, 0\}\,,$$

which is equivalent to decomposing the edge function $\varepsilon$ in the standard way:

$$\varepsilon = \varepsilon^{+} - \varepsilon^{-}\,, \quad \varepsilon^{+} := \max\{\varepsilon, \mathbf{0}\}\,, \quad \varepsilon^{-} := \max\{-\varepsilon, \mathbf{0}\}\,.$$

$\mathbf{A}^{+}/\varepsilon^{+}$ corresponds to its own network, denoted $\mathcal{G}^{+}$, the **positive subnetwork**. Likewise, $\mathbf{A}^{-}/\varepsilon^{-}$ also corresponds to its own network, denoted $\mathcal{G}^{-}$, the **negative subnetwork**. Cf. figures S2 and S3.

The edge weights of both $\mathcal{G}^{+}$ and $\mathcal{G}^{-}$ are by definition all positive, so standard dissimilarity measures for graphs can be applied to each subnetwork separately. This facilitates analysis and corresponds to the point of view that positive and negative edge weights correspond to qualitatively distinct phenomena.

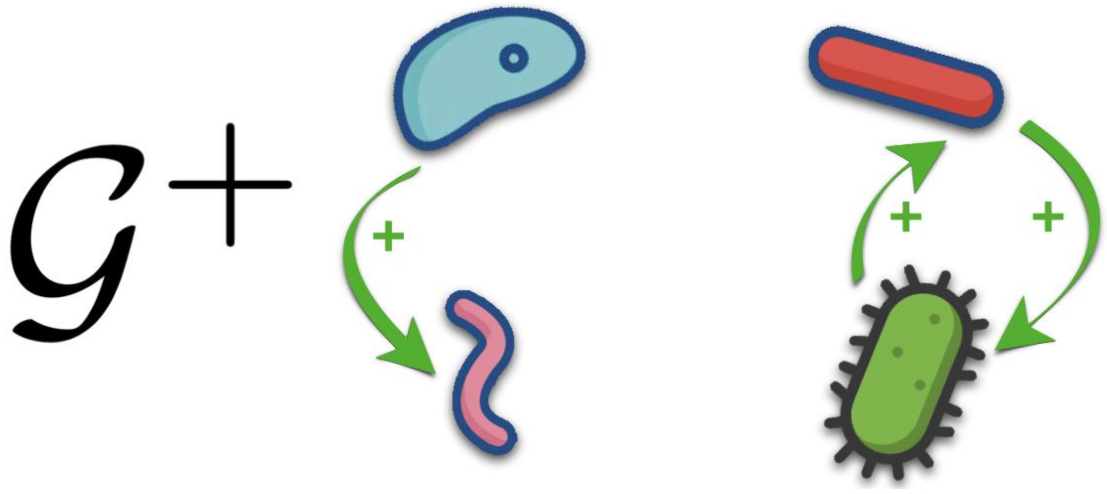

**Supplementary Figure S2.** Positive Subnetwork of the Network from Figure S1.

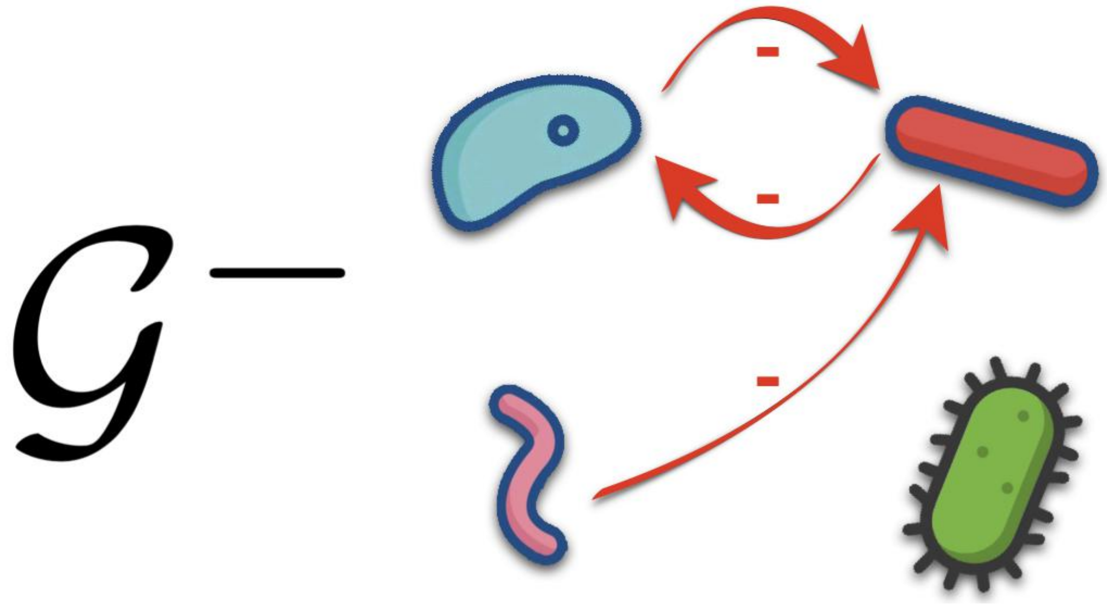

**Supplementary Figure S3.** Negative Subnetwork of the Network from Figure S1.

#### S2.5 Disjoint Union

Given two graphs, $\mathcal{G}_1$ with adjacency matrix $\mathbf{A}_1$ and $\mathcal{G}_2$ with adjacency matrix $\mathbf{A}_2$, define their "**disjoint union**" to be the graph $\mathcal{G}_1 \oplus \mathcal{G}_2$ corresponding to the adjacency matrix:

$$\mathbf{A}_1 \oplus \mathbf{A_2} = \begin{bmatrix} \mathbf{A}_1 & \mathbf{0} \\ \mathbf{0} & \mathbf{A}_2 \end{bmatrix},$$

i.e. the so-called direct sum of the matrices $\mathbf{A}_1$ and $\mathbf{A}_2$. The node set of this graph $\mathcal{G}_1 \oplus \mathcal{G}_2$ is the disjoint union of the node sets of $\mathcal{G}_1$ and $\mathcal{G}_2$, and likewise the edge set is the disjoint union of their edge sets $\mathcal{E}_1 \sqcup \mathcal{E}_2$. Cf. figure S4.

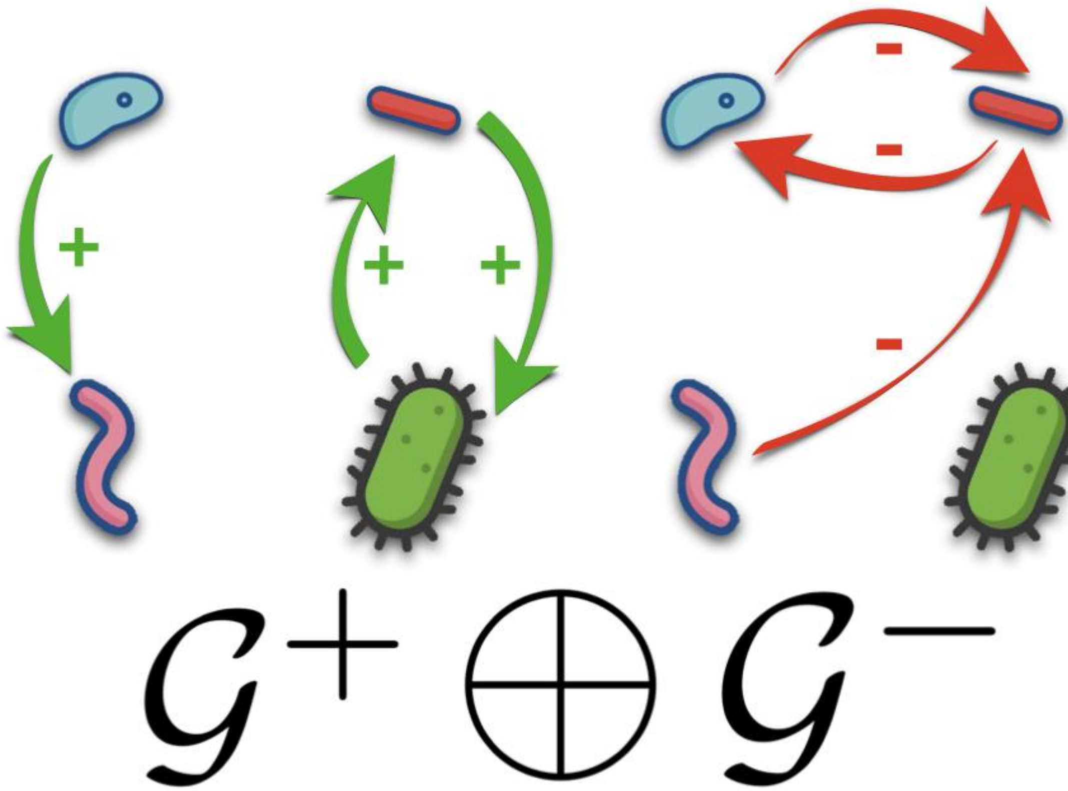

**Supplementary Figure S4.** Direct Sum of the Positive and Negative Subnetworks of the Network from Figure S1. Notice how the nodes are duplicated compared to Figure S1.

### S3 (DIS)SIMILARITY MEASURES

Below I define and discuss comparison methods applicable to networks with mixed-sign edge weights which span the full range of qualitative and quantitative comparisons explained in section **??**. When relevant, I explain how to modify methods for comparing unsigned networks to create new methods that give sensible results for networks with mixed-sign edge weights. I also explain the extent to which these proposed methods fall in line with the criteria discussed previously in sections **??**, S1.3, and S1.4.

Section S3.1 provides mostly “quantitative” comparisons of networks, section S3.2 provides mostly “qualitative” comparisons of relative orderings of edge weights, section S3.3 provides “qualitative” and “quantitative” comparisons of edge presence/absence, and finally section S3.4 provides “qualitative” and “quantitative” comparisons of path correspondence.

#### S3.1 (Entrywise $L_1$) Relative Error

Relative error is a “quantitative” comparison of networks. If we normalize the edge weights of both networks before computing their relative error, the comparison becomes somewhat more “qualitative” in the sense that the input edge-weight-normalized networks better capture the relative sizes of the edge weights. Herein I show that “out of the box” the entrywise $L_1$ relative error as defined below automatically considers both positive and negative edges separately and then sensibly recombines the results.

##### *S3.1.1 Entrywise $L_1$ Norm*

The entrywise $L_1$ norm of an $S \times S$ matrix $\mathbf{A}$ will herein be defined as

$$\|\mathbf{A}\|_{[1]} := \sum_{\sigma_1=1}^{S} \sum_{\sigma_2=1}^{S} |[\mathbf{A}]_{\sigma_1\sigma_2}| \,. \tag{S6}$$

In other words, this is the sum of the absolute values of the entries of $\mathbf{A}$, or the $L_1$ norm of a vector resulting from vectorizing $\mathbf{A}$. It can be considered the “$L_1$ equivalent” of the Frobenius norm.

It is *not* to be confused with $L_1$ *operator* norm of $\mathbf{A}$, which is typically denoted $\|\mathbf{A}\|_1$. The notation $\|\mathbf{A}\|_{[1]}$ serves not only to emphasize this distinction, but also to remind of the notation for the entries of a matrix.

An operator norm would be inappropriate to use here, given that the adjacency matrix $\mathbf{A}$ here is not meant to represent a linear function[4] (it is "entirely contravariant"). The Frobenius norm incorrectly "inflates" large-magnitude entries and "deflates" small-magnitude entries.

###### S3.1.2 Relative Error Definition

Given a "truth" matrix $\mathbf{A}_*$ and an "estimate" matrix $\hat{\mathbf{A}}$, the error of $\hat{\mathbf{A}}$ relative to $\mathbf{A}_*$, or just "relative error" if clear from context, is

$$\frac{\left\|\hat{\mathbf{A}} - \mathbf{A}_*\right\|_{[1]}}{\|\mathbf{A}_*\|_{[1]}} . \tag{S7}$$

The fact that relative error normalizes by the norm of one of the networks arguably helps to avoid considering two networks highly similar merely because they are both sparse. Cf. section S1.4. Normalization could help to ensure that a small number of differences is considered important if, due to sparsity, there are only a small number of edges that might be different to begin with.

###### S3.1.3 Convex Combination Decomposition Property

One can directly verify the identity:

$$\left\|\hat{\mathbf{A}} - \mathbf{A}_*\right\|_{[1]} = \left\|\hat{\mathbf{A}}^+ - \mathbf{A}_*^+\right\|_{[1]} + \left\|\hat{\mathbf{A}}^- - \mathbf{A}_*^-\right\|_{[1]} , \tag{S8}$$

which also happens to be a special case (for a counting measure) of the measure-theoretic identity $\int |f-g| \mathrm{d}\mu = \int |f^+ - g^+| \mathrm{d}\mu + \int |f^- - g^-| \mathrm{d}\mu$.

Therefore the overall relative error can be written as a convex combination/weighted average of the "positive relative error" and the "negative relative error". More specifically, the following is true:

$$\frac{\left\|\hat{\mathbf{A}} - \mathbf{A}_*\right\|_{[1]}}{\|\mathbf{A}_*\|_{[1]}} = c_1 \left( \frac{\left\|\hat{\mathbf{A}}^+ - \mathbf{A}_*^+\right\|_{[1]}}{\|\mathbf{A}_*^+\|_{[1]}} \right) + c_2 \left( \frac{\left\|\hat{\mathbf{A}}^- - \mathbf{A}_*^-\right\|_{[1]}}{\|\mathbf{A}_*^-\|_{[1]}} \right) , \tag{S9}$$

where the coefficients above are defined

$$c_1 := \frac{\|\mathbf{A}_*^+\|_{[1]}}{\|\mathbf{A}_*\|_{[1]}} , \quad c_2 := \frac{\|\mathbf{A}_*^-\|_{[1]}}{\|\mathbf{A}_*\|_{[1]}} . \tag{S10}$$

This also ensures that any positive interactions estimated as negative, or vice versa, will appear twice in the above expression. We can see how the convex combination property implies the double penalization principle is satisfied.

###### S3.1.4 Normalizing to get More Qualitative Comparisons

If a priori we don't believe that the edge weights of the two networks are on the same "length scale", e.g. they are measured in different units, but we still want to compare them in some qualitative, "scale-free" way, then we can normalize both networks' adjacency matrices before computing their relative error:

$$\frac{\left\| \frac{\hat{\mathbf{A}}}{\left\|\hat{\mathbf{A}}\right\|_{[1]}} - \frac{\mathbf{A}_*}{\|\mathbf{A}_*\|_{[1]}} \right\|_{[1]}}{\left\| \frac{\mathbf{A}_*}{\|\mathbf{A}_*\|_{[1]}} \right\|_{[1]}} = \frac{\left\| \frac{\hat{\mathbf{A}}}{\left\|\hat{\mathbf{A}}\right\|_{[1]}} - \frac{\mathbf{A}_*}{\|\mathbf{A}_*\|_{[1]}} \right\|_{[1]}}{1} = \left\| \frac{\hat{\mathbf{A}}}{\left\|\hat{\mathbf{A}}\right\|_{[1]}} - \frac{\mathbf{A}_*}{\|\mathbf{A}_*\|_{[1]}} \right\|_{[1]} . \tag{S11}$$

[4] Arguably except during the formation of the Neumann series in the definition of DeltaCon, cf. section S3.4.1.

For example, if we wanted to compare whether the relatively largest magnitude edge weights in both networks tended to have the same sign, or belong to edges connecting the same nodes, this could be useful. If an edge has the same sign, but three times the magnitude, in $\mathcal{G}_*$ compared to $\hat{\mathcal{G}}$, for example, that won't be penalized if the sum of the magnitudes of the edge weights is also three times as large in $\mathcal{G}_*$ compared to $\hat{\mathcal{G}}$ (hence why the comparison is "scale-free").

Note that the resulting values are necessarily always in the interval $[0,2]$. Also, unlike the relative error in general (S7), the resulting values are symmetric when interchanging $\hat{\mathbf{A}}$ and $\mathbf{A}_*$ in the formula (S11).

Normalizing both matrices first before computing their relative error arguably also helps to avoid considering two networks highly similar merely because they are both sparse. Cf. again section S1.4.

#### S3.2 Mixed-Sign Spearman Correlation

These methods based on Spearman correlation are designed to provide a qualitative comparison of two networks, by comparing the relative ordering of their edge weights. The formation of rank vectors destroys any direct quantitative information about exact edge weight values. The mixed-sign Spearman correlation defined below provides an advantage over "raw" Spearman correlation by separately comparing the relative ordering of positive and negative edges and then sensibly recombining the results. By considering this extra information contained in the signs of the edge weights, it is not fooled by drastic changes to the network structure to which the "raw" Spearman correlation or Spearman correlation of the weights' magnitudes are completely oblivious.

**Note:** by convention, correlations involving constant vectors are defined to be zero, rather than left undefined. I believe this convention makes sense because arguably constant vectors inherently have "zero predictive power".

##### S3.2.1 Sparsity Adjustment

Given two networks $\mathcal{G}_1$, $\mathcal{G}_2$ with adjacency matrices $\mathbf{A}_1$, $\mathbf{A}_2$ respectively, consider the subset of index pairs for which the edge corresponding to each index pair is present in at least one of the two networks:

$$\mathcal{I}_{\mathcal{G}_1,\mathcal{G}_2} := \{(s_1,s_2) \in [S]\times[S] : [\mathbf{A}_1]_{s_1 s_2} \neq 0 \text{ or } [\mathbf{A}_2]_{s_1 s_2} \neq 0\} . \tag{S12}$$

Define a modified vectorization operator $\text{vect}_{\mathcal{G}_1,\mathcal{G}_2}$ which, given $[S]\times[S]$ matrices, returns vectors of length $|\mathcal{I}_{\mathcal{G}_1,\mathcal{G}_2}|$ whose indices correspond to the index pairs belonging to $\mathcal{I}_{\mathcal{G}_1,\mathcal{G}_2}$. This is a "sparsity adjustment" because it removes values corresponding to index pairs for which no edge exists in either $\mathcal{G}_1$ or $\mathcal{G}_2$.

##### S3.2.2 Rank Vectors

Let $\mathcal{R}$ denote the "rank operator" which replaces each value of a vector with its relative ordering (e.g. the smallest element is assigned 1, the largest is assigned the length of the vector). Assume $\mathcal{R}$ is such that ties are replaced with the mean value of the tied positions (as is done by default e.g. in the 'rankdata' function of the SciPy stats library(Virtanen et al., 2020)).

Given a network $\mathcal{G}$ with adjacency matrix $\mathbf{A}$, define the following shorthand:

$$\underset{\mathcal{G}_1,\mathcal{G}_2}{\mathcal{R}}(\mathcal{G}) := \mathcal{R}\left(\text{vect}_{\mathcal{G}_1,\mathcal{G}_2}(\mathbf{A})\right) , \tag{S13}$$

where $\text{vect}_{\mathcal{G}_1,\mathcal{G}_2}$ is the same sparsity-adjusted vectorization operator that was defined in section S3.2.1.

##### S3.2.3 Positive and Negative Spearman Correlation

The "**positive Spearman correlation**" of $\mathcal{G}_1$ and $\mathcal{G}_2$ is defined to be the (sparsity-adjusted) Spearman correlation[5] of $\mathcal{G}_1^+$ and $\mathcal{G}_2^+$:

$$\rho_S^+(\mathcal{G}_1,\mathcal{G}_2) := \frac{\widehat{\text{cov}}\left[\underset{\mathcal{G}_1,\mathcal{G}_2}{\mathcal{R}}(\mathcal{G}_1^+), \underset{\mathcal{G}_1,\mathcal{G}_2}{\mathcal{R}}(\mathcal{G}_2^+)\right]}{\sqrt{\widehat{\text{var}}\left[\underset{\mathcal{G}_1,\mathcal{G}_2}{\mathcal{R}}(\mathcal{G}_1^+)\right]}\cdot\sqrt{\widehat{\text{var}}\left[\underset{\mathcal{G}_1,\mathcal{G}_2}{\mathcal{R}}(\mathcal{G}_2^+)\right]}} . \tag{S14}$$

[5]The "$S$" in the notation "$\rho_S$" can be interpreted to refer to either "Spearman", "sparsity-adjusted", or both. "$\rho$" of course is commonly-used notation for a correlation coefficient.

Notice that the sparsity adjustment is in terms of $\mathcal{G}_1$ and $\mathcal{G}_2$, not $\mathcal{G}_1^+$ and $\mathcal{G}_2^+$. This is noteworthy to the extent that the union of the edge sets of $\mathcal{G}_1^+$ and $\mathcal{G}_2^+$ could, in principle, be strictly smaller than the union of the edge sets of $\mathcal{G}_1$ and $\mathcal{G}_2$. Therefore one might sensibly argue that the strictest possible sparsity adjustment, that in terms of $\mathcal{G}_1^+$ and $\mathcal{G}_2^+$, should be applied when defining a notion of "positive Spearman correlation". Anything else might not be compatible with the principle outlined in section S1.4. On the other hand, one could also argue that a correlation which "preserves the context of the original networks" admits a more useful interpretation, at least when trying to compare or combine[6] the values of the positive and negative Spearman correlations.

Completely analogously, the "**negative Spearman correlation**" of $\mathcal{G}_1$ and $\mathcal{G}_2$ is defined to be the (sparsity-adjusted) Spearman correlation of $\mathcal{G}_1^-$ and $\mathcal{G}_2^-$:

$$\rho_S^-(\mathcal{G}_1,\mathcal{G}_2) := \frac{\widehat{\text{cov}}\left[\underset{\mathcal{G}_1,\mathcal{G}_2}{\mathcal{R}}(\mathcal{G}_1^-),\underset{\mathcal{G}_1,\mathcal{G}_2}{\mathcal{R}}(\mathcal{G}_2^-)\right]}{\sqrt{\widehat{\text{var}}\left[\underset{\mathcal{G}_1,\mathcal{G}_2}{\mathcal{R}}(\mathcal{G}_1^-)\right]}\cdot\sqrt{\widehat{\text{var}}\left[\underset{\mathcal{G}_1,\mathcal{G}_2}{\mathcal{R}}(\mathcal{G}_2^-)\right]}} \tag{S15}$$

Analogous comments about sparsity adjustments apply here too, of course.

###### *S3.2.4 Vector Concatenation*

The concatenation or "direct sum" $\vec{\mathbf{v}}_1 \oplus \vec{\mathbf{v}}_2 \in \mathbb{R}^{I_1+I_2}$ of two vectors $\vec{\mathbf{v}}_1 \in \mathbb{R}^{I_1}$ and $\vec{\mathbf{v}}_2 \in \mathbb{R}^{I_2}$ is defined as

$$[\vec{\mathbf{v}}_1 \oplus \vec{\mathbf{v}}_2]_i := \begin{cases} [\vec{\mathbf{v}}_1]_i & 1 \le i \le I_1 \\ [\vec{\mathbf{v}}_2]_{i-I_1} & I_1+1 \le i \le I_1+I_2\,. \end{cases} \tag{S16}$$

This has the intuitive interpretation of first listing the entries of $\vec{\mathbf{v}}_1$ and then appending to that list the entries of $\vec{\mathbf{v}}_2$.

Some properties are obvious consequences of the definition. For example:

$$(\vec{\mathbf{v}}_1+\vec{\mathbf{w}}_1)\oplus(\vec{\mathbf{v}}_2+\vec{\mathbf{w}}_2) = (\vec{\mathbf{v}}_1\oplus\vec{\mathbf{v}}_2)+(\vec{\mathbf{w}}_1\oplus\vec{\mathbf{w}}_2)\,. \tag{S17}$$

Here is a second example using the standard inner product, $\langle\vec{\mathbf{x}},\vec{\mathbf{y}}\rangle := \sum_{i=1}^{I}[\vec{\mathbf{x}}]_i\cdot[\vec{\mathbf{y}}]_i$:

$$\langle\vec{\mathbf{v}}_1\oplus\vec{\mathbf{v}}_2,\vec{\mathbf{w}}_1\oplus\vec{\mathbf{w}}_2\rangle = \langle\vec{\mathbf{v}}_1,\vec{\mathbf{w}}_1\rangle+\langle\vec{\mathbf{v}}_2,\vec{\mathbf{w}}_2\rangle\,. \tag{S18}$$

As seen above, for many operations on concatenated vectors in $\mathbb{R}^{I_1+I_2}$ we can treat what happens in $\mathbb{R}^{I_1}$ separately of what happens in $\mathbb{R}^{I_2}$.

###### *S3.2.5 Mixed-Sign Spearman Definition*

Without explaining or motivating anything, the definition may be written as

$$\rho_S^\pm(\mathcal{G}_1,\mathcal{G}_2) := \frac{\widehat{\text{cov}}\left[\underset{\mathcal{G}_1,\mathcal{G}_2}{\mathcal{R}}(\mathcal{G}_1^+)\oplus\underset{\mathcal{G}_1,\mathcal{G}_2}{\mathcal{R}}(\mathcal{G}_1^-),\underset{\mathcal{G}_1,\mathcal{G}_2}{\mathcal{R}}(\mathcal{G}_2^+)\oplus\underset{\mathcal{G}_1,\mathcal{G}_2}{\mathcal{R}}(\mathcal{G}_2^-)\right]}{\sqrt{\widehat{\text{var}}\left[\underset{\mathcal{G}_1,\mathcal{G}_2}{\mathcal{R}}(\mathcal{G}_1^+)\oplus\underset{\mathcal{G}_1,\mathcal{G}_2}{\mathcal{R}}(\mathcal{G}_1^-)\right]}\cdot\sqrt{\widehat{\text{var}}\left[\underset{\mathcal{G}_1,\mathcal{G}_2}{\mathcal{R}}(\mathcal{G}_2^+)\oplus\underset{\mathcal{G}_1,\mathcal{G}_2}{\mathcal{R}}(\mathcal{G}_2^-)\right]}}\,. \tag{S19}$$

Using the sparsity-adjusted rank operator, instead of the typical rank operator $\mathcal{R}$, prevents two networks from scoring high values merely because both are highly sparse. This is because the sparsity-adjusted rank operator only looks at the subset of edges which are nonzero in at least one of the networks, and does not consider edges missing from both networks. Thus the use of the sparsity-adjusted rank operator follows from the principle outlined in section S1.4.

[6] Admittedly my main motivation for defining the sparsity adjustments for the positive and negative Spearman correlations using the edge sets of the original networks was to facilitate the interpretation of the mixed-sign Spearman correlation. If the sparsity adjustment for the positive Spearman correlation was in terms of $\mathcal{G}_1^+$ and $\mathcal{G}_2^+$, and that for the negative Spearman correlation in terms of $\mathcal{G}_1^-$ and $\mathcal{G}_2^-$, then as far as I can tell there would in general be no straightforward way to write the mixed-sign Spearman correlation as a weighted sum of the positive and negative Spearman correlations. I came up with the notion of mixed-sign Spearman correlation before observing the need for sparsity adjustments, so it perhaps is not altogether surprising that the two notions do not seem to be easily compatible.

###### *S3.2.6 Subconvex Combination Decomposition Property*

The sense in which expression (S19) considers the positive and negative subnetworks separately and then combines the results is made precise below.

Because of Lemma S7.2, the hypotheses of Lemma S7.3 are satisfied. Applying Lemma S7.3 to expressions in both the numerator and denominator of (S19):

$$
\begin{aligned}
\rho_S^{\pm}(\mathcal{G}_1,\mathcal{G}_2) &= \frac{\widehat{\text{cov}}\left[\underset{\mathcal{G}_1,\mathcal{G}_2}{\mathcal{R}}(\mathcal{G}_1^+),\underset{\mathcal{G}_1,\mathcal{G}_2}{\mathcal{R}}(\mathcal{G}_2^+)\right]+\widehat{\text{cov}}\left[\underset{\mathcal{G}_1,\mathcal{G}_2}{\mathcal{R}}(\mathcal{G}_1^-),\underset{\mathcal{G}_1,\mathcal{G}_2}{\mathcal{R}}(\mathcal{G}_2^-)\right]}{\sqrt{\widehat{\text{var}}\left[\underset{\mathcal{G}_1,\mathcal{G}_2}{\mathcal{R}}(\mathcal{G}_1^+)\right]+\widehat{\text{var}}\left[\underset{\mathcal{G}_1,\mathcal{G}_2}{\mathcal{R}}(\mathcal{G}_1^-)\right]}\cdot\sqrt{\widehat{\text{var}}\left[\underset{\mathcal{G}_1,\mathcal{G}_2}{\mathcal{R}}(\mathcal{G}_2^+)\right]+\widehat{\text{var}}\left[\underset{\mathcal{G}_1,\mathcal{G}_2}{\mathcal{R}}(\mathcal{G}_2^-)\right]}} \\
&= c_+\frac{\widehat{\text{cov}}\left[\underset{\mathcal{G}_1,\mathcal{G}_2}{\mathcal{R}}(\mathcal{G}_1^+),\underset{\mathcal{G}_1,\mathcal{G}_2}{\mathcal{R}}(\mathcal{G}_2^+)\right]}{\sqrt{\widehat{\text{var}}\left[\underset{\mathcal{G}_1,\mathcal{G}_2}{\mathcal{R}}(\mathcal{G}_1^+)\right]}\cdot\sqrt{\widehat{\text{var}}\left[\underset{\mathcal{G}_1,\mathcal{G}_2}{\mathcal{R}}(\mathcal{G}_2^+)\right]}}+c_-\frac{\widehat{\text{cov}}\left[\underset{\mathcal{G}_1,\mathcal{G}_2}{\mathcal{R}}(\mathcal{G}_1^-),\underset{\mathcal{G}_1,\mathcal{G}_2}{\mathcal{R}}(\mathcal{G}_2^-)\right]}{\sqrt{\widehat{\text{var}}\left[\underset{\mathcal{G}_1,\mathcal{G}_2}{\mathcal{R}}(\mathcal{G}_1^-)\right]}\cdot\sqrt{\widehat{\text{var}}\left[\underset{\mathcal{G}_1,\mathcal{G}_2}{\mathcal{R}}(\mathcal{G}_2^-)\right]}} \\
&= c_+\cdot\rho_S^+(\mathcal{G}_1,\mathcal{G}_2)+c_-\cdot\rho_S^-(\mathcal{G}_1,\mathcal{G}_2),
\end{aligned}
\tag{S20}
$$

where the coefficients $c_+$, $c_-$ equal

$$
\begin{aligned}
c_+ &= \frac{\sqrt{\widehat{\text{var}}\left[\underset{\mathcal{G}_1,\mathcal{G}_2}{\mathcal{R}}(\mathcal{G}_1^+)\right]}\cdot\sqrt{\widehat{\text{var}}\left[\underset{\mathcal{G}_1,\mathcal{G}_2}{\mathcal{R}}(\mathcal{G}_2^+)\right]}}{\sqrt{\widehat{\text{var}}\left[\underset{\mathcal{G}_1,\mathcal{G}_2}{\mathcal{R}}(\mathcal{G}_1^+)\right]+\widehat{\text{var}}\left[\underset{\mathcal{G}_1,\mathcal{G}_2}{\mathcal{R}}(\mathcal{G}_1^-)\right]}\cdot\sqrt{\widehat{\text{var}}\left[\underset{\mathcal{G}_1,\mathcal{G}_2}{\mathcal{R}}(\mathcal{G}_2^+)\right]+\widehat{\text{var}}\left[\underset{\mathcal{G}_1,\mathcal{G}_2}{\mathcal{R}}(\mathcal{G}_2^-)\right]}}, \\
c_- &= \frac{\sqrt{\widehat{\text{var}}\left[\underset{\mathcal{G}_1,\mathcal{G}_2}{\mathcal{R}}(\mathcal{G}_1^-)\right]}\cdot\sqrt{\widehat{\text{var}}\left[\underset{\mathcal{G}_1,\mathcal{G}_2}{\mathcal{R}}(\mathcal{G}_2^-)\right]}}{\sqrt{\widehat{\text{var}}\left[\underset{\mathcal{G}_1,\mathcal{G}_2}{\mathcal{R}}(\mathcal{G}_1^+)\right]+\widehat{\text{var}}\left[\underset{\mathcal{G}_1,\mathcal{G}_2}{\mathcal{R}}(\mathcal{G}_1^-)\right]}\cdot\sqrt{\widehat{\text{var}}\left[\underset{\mathcal{G}_1,\mathcal{G}_2}{\mathcal{R}}(\mathcal{G}_2^+)\right]+\widehat{\text{var}}\left[\underset{\mathcal{G}_1,\mathcal{G}_2}{\mathcal{R}}(\mathcal{G}_2^-)\right]}}.
\end{aligned}
\tag{S21}
$$

Although in equation (S21) we have $c_+, c_- \geq 0$, in general we only have that $c_+ + c_- \leq 1$, cf. Lemma S7.4. In other words, in general the mixed-sign Spearman correlation is only guaranteed to be a **subconvex combination** of the positive Spearman correlation and the negative Spearman correlation. In special cases it may still also be a convex combination sensu stricto. Because this subconvex property can be derived using the Cauchy-Schwarz inequality, one way to interpret any deficit $1-(c_+ + c_-) > 0$ is as a measurement of how much[7] $\mathcal{G}_1$ and $\mathcal{G}_2$ "fail to align in almost the same direction".

Although the subconvex property does still correspond to a weighted sum, and does have some merit for possessing a useful interpretation, it nevertheless is still fair to argue that the mixed-sign Spearman correlation does not fully live up to the promised easy interpretation from section S1.3. The mixed-sign Spearman correlation does still satisfy the double penalization principle. Despite in general being a subconvex combination but not necessarily a convex combination, in practice the mixed-sign Spearman correlation also does not seem to necessarily give overly "conservative" results "biased" towards 0.

Perhaps one could argue that any such "shrinkage" might even be desirable, given that the sparsity adjustments defining the positive and negative Spearman correlations are not as aggressive as possible (cf. section S3.2.3). By possibly failing to adhere to the principle of section S1.4, one might be concerned that the positive and negative Spearman correlations artifactually overstate similarities. While this argument might be true to an extent, I would also caution that "two wrongs don't make a right". A priori we have no reason to believe that hypothetical "corrections" from the subconvex property would (or even could) exactly counterbalance hypothetical artifacts from the chosen sparsity adjustments for the positive and negative Spearman correlations.

[7] More precisely, it measures how much $\underset{\mathcal{G}_1,\mathcal{G}_2}{\mathcal{R}}(\mathcal{G}_1^+)\oplus\underset{\mathcal{G}_1,\mathcal{G}_2}{\mathcal{R}}(\mathcal{G}_1^-)$ and $\underset{\mathcal{G}_1,\mathcal{G}_2}{\mathcal{R}}(\mathcal{G}_2^+)\oplus\underset{\mathcal{G}_1,\mathcal{G}_2}{\mathcal{R}}(\mathcal{G}_2^-)$ "fail to almost align onto the same line". The Cauchy-Schwarz inequality is an equality only when two vectors are linearly dependent, so in the same (or exactly opposite) directions.

##### S3.3 Mixed-Sign Jaccard Similarity

Unweighted Jaccard similarity provides a qualitative comparison of networks by comparing the presence/absence of edges. Weighted Jaccard similarity provides a comparison that is a little quantitative and a little qualitative, by increasing the importance of comparisons for edge pairs where one of the edges has a large edge weight. Mixed-sign versions of both comparison methods provide an advantage over unsigned versions by separately considering presence/absence for positive and negative edges and then sensibly recombining the results. By considering this extra information contained in the signs of the edge weights, the proposed mixed-sign versions are not fooled by drastic changes to the network structure to which the unsigned versions are completely oblivious.

###### *S3.3.1 Binary Classification and Mixed Signs*

Considering positive and negative edges to correspond to distinct phenomena means we effectively have a ternary classification problem (present and positive, present and negative, absent). Our goal is to recast this ternary classification problem as two inter-related binary classification problems, one for the positive edges and one for the negative edges.

To avoid confusion with the edge signs, below I use the following alternative terminology to refer to the four possible binary classification outcomes:

- ”**True Hit**” (TH) := ”True Positive”,
- ”**False Hit**” (FH) := ”False Positive”,
- ”**True Miss**” (TM) := ”True Negative”,
- ”**False Miss**” (FM) := ”False Negative”.

###### *S3.3.2 Positive Edge Binary Classification Problem*

The positive edge binary classification problem has

- hits = $\{+\}$,
- misses = $\{0, -\}$.

So for the positive edge binary classification problem:

- a “**true hit**” occurs when both the truth and the estimate are positive,
- a “**false hit**” occurs when the truth is non-positive (0 or $-$) but the estimate is positive,
- a “**true miss**” occurs when the truth is non-positive and the estimate is non-positive, and
- a “**false miss**” occurs when the truth is positive but the estimate is non-positive.

###### *S3.3.3 Negative Edge Binary Classification Problem*

Analogously the negative edge binary classification problem has

- hits = $\{-\}$,
- misses = $\{0, +\}$.

So for the negative edge binary classification problem:

- a “**true hit**” occurs when both the truth and the estimate are negative,
- a “**false hit**” occurs when the truth is non-negative (0 or $+$) but the estimate is negative,
- a “**true miss**” occurs when the truth is non-negative and the estimate is non-negative, and
- a “**false miss**” occurs when the truth is negative but the estimate is non-negative.

###### S3.3.4 Origin of Double Penalization

From this perspective of having two inter-related binary classification problems, confusing a positive interaction with a negative interaction, or vice versa, is arguably both

- a **false hit**, inferring a type of interaction which does not exist, and
- a **false miss**, failing to infer a type of interaction which does exist.

Thus an ideal performance metric should "doubly penalize" such errors. This was the original insight which led to the more general double penalization principle from section **??**.

###### S3.3.5 (Unsigned) Unweighted Definition

The (unweighted) Jaccard similarity of two graphs $\mathcal{G}_1$ and $\mathcal{G}_2$ (with non-negative edge weights) is the Jaccard similarity of their edge sets $\mathcal{E}_1$ and $\mathcal{E}_2$:

$$\mathcal{S}_J(\mathcal{G}_1, \mathcal{G}_2) := \frac{|\mathcal{E}_1 \cap \mathcal{E}_2|}{|\mathcal{E}_1 \cup \mathcal{E}_2|} = \frac{|\mathcal{E}_1 \cap \mathcal{E}_2|}{|\mathcal{E}_1 \cap \mathcal{E}_2| + |\mathcal{E}_1 \setminus \mathcal{E}_2| + |\mathcal{E}_2 \setminus \mathcal{E}_1|}\,. \tag{S22}$$

Recall that

$$\mathcal{E}_1 := \text{supp}(\varepsilon_1)\,, \mathcal{E}_2 := \text{supp}(\varepsilon_2)$$

where $\varepsilon_1 \geq 0$ is the edge function corresponding to $\mathcal{G}_1$, and likewise for $\varepsilon_2 \geq 0$.

Notice that if $\mathcal{G}_1$ is an "estimate" $\hat{\mathcal{G}}$, and $\mathcal{G}_2$ is the "ground truth" $\mathcal{G}_*$ then their (unweighted) Jaccard similarity corresponds to

$$\mathcal{S}_J(\hat{\mathcal{G}}, \mathcal{G}_*) = \frac{TH}{TH + FH + FM}\,, \tag{S23}$$

where $TH$ is the number of true hits, $FH$ is the number of false hits, and $FM$ is the number of false misses. Thus Jaccard similarity is no less conservative than the minimum of precision and recall, since it has the same numerator and one additional non-negative term in the denominator.

It is beneficial that Jaccard similarity does not reward true misses[8], because not doing so helps to prevent considering two highly different networks as similar merely because both are highly sparse. Cf. again section S1.4.

As an aside, in meteorology Jaccard similarity (as used for binary classification) is often called the "critical success index" or "threat score"(Hogan et al., 2010). Many binary classification metrics considered improvements over Jaccard similarity (e.g. the "equitable threat score") are studied in meteorology(Hogan et al., 2010). Future work examining whether these other binary classification metrics also admit extensions to the comparison of networks of mixed-sign edge weights, and whether such extensions would be useful, would be interesting.

###### S3.3.6 (Unsigned) Weighted Definition

For two graphs $\mathcal{G}_1$ and $\mathcal{G}_2$ with non-negative edge weights and edge functions $\varepsilon_1$ and $\varepsilon_2$ as above, their (unsigned) weighted Jaccard similarity is

$$\mathcal{S}_J^W(\mathcal{G}_1, \mathcal{G}_2) := \frac{\sum_{i,j} \min\{\varepsilon_1(i,j), \varepsilon_2(i,j)\}}{\sum_{i,j} \max\{\varepsilon_1(i,j), \varepsilon_2(i,j)\}}\,. \tag{S24}$$

By considering the graphs' "skeletons", which have edge functions $(\text{sign} \circ \varepsilon_1)$ and $(\text{sign} \circ \varepsilon_2)$, one recovers the regular (unweighted) Jaccard similarity. Specifically

$$\mathcal{S}_J^W(\mathbb{S}(\mathcal{G}_1), \mathbb{S}(\mathcal{G}_2)) = \mathcal{S}_J(\mathcal{G}_1, \mathcal{G}_2)\,. \tag{S25}$$

Recall the definitions of graph (unsigned) skeletons from section S2.3.1.

[8] $TM$ would denote the number of true misses. Notably this is absent from the numerator of equation (S23). Thus higher values of $TM$ do not lead to higher values of Jaccard similarity.

##### S3.3.7 Positive and Negative Jaccard Similarities

From here on let $\mathcal{G}_1$, $\mathcal{G}_2$ denote two general networks with *mixed-sign* edge weights. (In the previous sections S3.3.5 and S3.3.6 it was assumed for the sake of simplicity that the networks $\mathcal{G}_1$, $\mathcal{G}_2$ had non-negative edge weights.)

One can define the "positive (unweighted) Jaccard similarity" of two general networks as:

$$\mathcal{S}_J^+(\mathcal{G}_1,\mathcal{G}_2) := \mathcal{S}_J(\mathcal{G}_1^+,\mathcal{G}_2^+)\,, \tag{S26}$$

as well as their "positive weighted Jaccard similarity",

$$(\mathcal{S}_J^W)^+(\mathcal{G}_1,\mathcal{G}_2) := \mathcal{S}_J^W(\mathcal{G}_1^+,\mathcal{G}_2^+)\,. \tag{S27}$$

Completely analogously, the "negative (unweighted) Jaccard similarity" of two general networks is:

$$\mathcal{S}_J^-(\mathcal{G}_1,\mathcal{G}_2) := \mathcal{S}_J(\mathcal{G}_1^-,\mathcal{G}_2^-)\,, \tag{S28}$$

and their "negative weighted Jaccard similarity",

$$(\mathcal{S}_J^W)^-(\mathcal{G}_1,\mathcal{G}_2) := \mathcal{S}_J^W(\mathcal{G}_1^-,\mathcal{G}_2^-)\,. \tag{S29}$$

The above definitions make sense because the positive $\mathcal{G}^+$ and negative subnetworks $\mathcal{G}^-$ have non-negative edge weights by definition. Therefore "standard" network dissimilarity methods may be applied to them.

Observe how the weighted versions $(\mathcal{S}_J^W)^+$, $(\mathcal{S}_J^W)^-$ reduce to their respective unweighted counterparts $\mathcal{S}_J^+$, $\mathcal{S}_J^-$ when applied to (signed) skeletons:

$$(\mathcal{S}_J^W)^+(\mathbb{S}^\pm(\mathcal{G}_1),\mathbb{S}^\pm(\mathcal{G}_2)) = \mathcal{S}_J^+(\mathcal{G}_1,\mathcal{G}_2)\,, \quad (\mathcal{S}_J^W)^-(\mathbb{S}^\pm(\mathcal{G}_1),\mathbb{S}^\pm(\mathcal{G}_2)) = \mathcal{S}_J^-(\mathcal{G}_1,\mathcal{G}_2)\,. \tag{S30}$$

##### S3.3.8 Mixed-Sign Unweighted Jaccard Similarity

The "mixed sign (unweighted) Jaccard similarity" of two networks $\mathcal{G}_1$, $\mathcal{G}_2$ with mixed-sign edge weights is defined to equal

$$\mathcal{S}_J^\pm(\mathcal{G}_1,\mathcal{G}_2) := \mathcal{S}_J(\mathcal{G}_1^+ \oplus \mathcal{G}_1^-, \mathcal{G}_2^+ \oplus \mathcal{G}_2^-) = \frac{|\mathcal{E}_1^+ \cap \mathcal{E}_2^+| + |\mathcal{E}_1^- \cap \mathcal{E}_2^-|}{|\mathcal{E}_1^+ \cup \mathcal{E}_2^+| + |\mathcal{E}_1^- \cup \mathcal{E}_2^-|}\,. \tag{S31}$$

($\mathcal{E}^+$ is the edge set of a graph's positive part, analogously for $\mathcal{E}^-$.)

Notice that if a positive edge is estimated as negative, it will show up once both as a false hit for the negative edges (contributing once to $|\mathcal{E}_1^- \cup \mathcal{E}_2^-|$) as well as a false miss for the positive edges (contributing once to $|\mathcal{E}_1^+ \cup \mathcal{E}_2^+|$). The analogous conclusions hold when a negative edge is estimated as positive.

Therefore, *mixed-sign (unweighted) Jaccard similarity satisfies the desired double penalization property for incorrectly estimated signs*. In fact:

$$\mathcal{S}_J^\pm(\hat{\mathcal{G}},\mathcal{G}_*) = \frac{TH^+ + TH^-}{TH^+ + TH^- + FH^+ + FH^- + FM^+ + FM^-}\,. \tag{S32}$$

The analogy between (S23) and (S32) strongly suggests that mixed-sign Jaccard similarity is "the correct" generalization of (unsigned) Jaccard similarity.

Notice how this definition again does not reward true misses[9], which could occur merely due to either the positive and/or negative subnetworks being highly sparse. This helps mixed-sign (unweighted) Jaccard similarity to avoid considering two highly different networks as similar merely because both are highly sparse, and/or both have few positive/negative edges. Cf. again section S1.4.

Moreover the mixed-sign (unweighted) Jaccard similarity can be described simply as a convex combination/weighted average of the positive (unweighted) Jaccard similarity and the negative (unweighted) Jaccard similarity:

$$\mathcal{S}_J^\pm(\mathcal{G}_1,\mathcal{G}_2) = c_+ \mathcal{S}_J^+(\mathcal{G}_1,\mathcal{G}_2) + c_- \mathcal{S}_J^-(\mathcal{G}_1,\mathcal{G}_2)\,, \tag{S33}$$

with the coefficients of the convex combination above being

$$c_+ = \frac{|\mathcal{E}_1^+ \cup \mathcal{E}_2^+|}{|\mathcal{E}_1^+ \cup \mathcal{E}_2^+| + |\mathcal{E}_1^- \cup \mathcal{E}_2^-|}\,, \quad c_- = \frac{|\mathcal{E}_1^- \cup \mathcal{E}_2^-|}{|\mathcal{E}_1^+ \cup \mathcal{E}_2^+| + |\mathcal{E}_1^- \cup \mathcal{E}_2^-|}\,. \tag{S34}$$

[9] $TM^+$ would denote the number of true misses for the positive subnetworks, $TM^-$ would denote the number of true misses for the negative subnetworks. Notably these are absent from the numerator of equation (S32). Thus neither higher values of $TM^+$, nor higher values of $TM^-$, lead to higher values of mixed-sign Jaccard similarity.

###### *S3.3.9 Mixed-Sign Weighted Jaccard Similarity*

Completely analogously the "mixed sign weighted Jaccard similarity" of two networks $\mathcal{G}_1$, $\mathcal{G}_2$ with mixed-sign edge weights is defined to equal

$$(\mathcal{S}_J^W)^{\pm}(\mathcal{G}_1,\mathcal{G}_2) := \mathcal{S}_J^W(\mathcal{G}_1^+ \oplus \mathcal{G}_1^-, \mathcal{G}_2^+ \oplus \mathcal{G}_2^-) = \frac{\sum_{i,j}\min\{\varepsilon_1^+(i,j),\varepsilon_2^+(i,j)\} + \sum_{i,j}\min\{\varepsilon_1^-(i,j),\varepsilon_2^-(i,j)\}}{\sum_{i,j}\max\{\varepsilon_1^+(i,j),\varepsilon_2^+(i,j)\} + \sum_{i,j}\max\{\varepsilon_1^-(i,j),\varepsilon_2^-(i,j)\}}. \tag{S35}$$

Like the unsigned version (cf. section S3.3.6), the mixed-sign weighted Jaccard similarity reduces to its unweighted counterpart when applied to skeletons:

$$(\mathcal{S}_J^W)^{\pm}(\mathcal{S}^{\pm}(\mathcal{G}_1),\mathcal{S}^{\pm}(\mathcal{G}_2)) = \mathcal{S}_J^{\pm}(\mathcal{G}_1,\mathcal{G}_2). \tag{S36}$$

Like its unweighted counterpart, the mixed-sign weighted Jaccard similarity also enjoys the convex combination decomposition property:

$$(\mathcal{S}_J^W)^{\pm}(\mathcal{G}_1,\mathcal{G}_2) = c_+(\mathcal{S}_J^W)^+(\mathcal{G}_1,\mathcal{G}_2) + c_-(\mathcal{S}_J^W)^-(\mathcal{G}_1,\mathcal{G}_2), \tag{S37}$$

with the coefficients $c_+$, $c_-$ of the convex combination being

$$\begin{aligned} c_+ &:= \frac{\sum_{i,j}\max\{\varepsilon_1^+(i,j),\varepsilon_2^+(i,j)\}}{\sum_{i,j}\max\{\varepsilon_1^+(i,j),\varepsilon_2^+(i,j)\} + \sum_{i,j}\max\{\varepsilon_1^-(i,j),\varepsilon_2^-(i,j)\}}, \\ c_- &:= \frac{\sum_{i,j}\max\{\varepsilon_1^-(i,j),\varepsilon_2^-(i,j)\}}{\sum_{i,j}\max\{\varepsilon_1^+(i,j),\varepsilon_2^+(i,j)\} + \sum_{i,j}\max\{\varepsilon_1^-(i,j),\varepsilon_2^-(i,j)\}}. \end{aligned} \tag{S38}$$

Therefore, like its unweighted counterpart, mixed-sign weighted Jaccard similarity also satisfies the double penalization principle.

###### *S3.3.10 Mixed-Sign False Discovery Rate and Mixed-Sign False Miss Rate*

Herein assume again that $\mathcal{G}_1$ is an "estimate" $\hat{\mathcal{G}}$, and $\mathcal{G}_2$ is the "ground truth", and both have non-negative edge weights. Similar to the "measure of goodness" defined by (S23), we can also define two corresponding "measures of badness":

- the **false discovery rate**:

$$\text{FDR}(\hat{\mathcal{G}},\mathcal{G}_*) := \frac{FH}{TH+FH}, \tag{S39}$$

- and the **false miss rate**[10]:

$$\text{FMR}(\hat{\mathcal{G}},\mathcal{G}_*) := \frac{FM}{TH+FM}. \tag{S40}$$

Note that, unlike the definition of Jaccard similarity, these definitions are not symmetric with respect to the roles of $\hat{\mathcal{G}}$ and $\mathcal{G}_*$.

Now allow $\hat{\mathcal{G}}$ and $\mathcal{G}_*$ to again have mixed-sign edge weights. Analogous to the relationship between definitions (S23) and (S32), we can also define mixed-sign versions of the false discovery rate (S39) and the false miss rate (S42):

- the **mixed-sign false discovery rate**:

$$\text{FDR}^{\pm}(\hat{\mathcal{G}},\mathcal{G}_*) := \frac{FH^+ + FH^-}{TH^+ + TH^- + FH^+ + FH^-}, \tag{S41}$$

[10] More typically called "false negative rate", but that would be confusing in this context.

- and the **mixed-sign false miss rate**:

$$\mathrm{FMR}^{\pm}(\hat{\mathcal{G}}, \mathcal{G}_*) := \frac{FM^+ + FM^-}{TH^+ + TH^- + FM^+ + FM^-}. \tag{S42}$$

Just like the mixed-sign Jaccard similarity, these can also be written straightforwardly as weighted averages of the corresponding single-sign performance metrics applied to the positive $\hat{\mathcal{G}}^+, \mathcal{G}_*^+$ and negative $\hat{\mathcal{G}}^-, \mathcal{G}_*^-$ subnetworks.

#### S3.4 Mixed-Sign DeltaCon Distance

Unweighted DeltaCon provides a qualitative comparison of networks by quantifying the correspondence of their paths. Weighted DeltaCon provides a comparison that is a little quantitative and a little qualitative, by increasing the importance of comparisons for paths where some of the edges have large edge weights. Herein mixed-sign versions of both unweighted and weighted DeltaCon are proposed which are not fooled by drastic changes to the network structure to which the unsigned versions from previous work are completely oblivious.

##### S3.4.1 Paths and Powers of Adjacency Matrices

Given an adjacency matrix as defined in section S2.1, for any $n \geq 1$, the matrix $\mathbf{A}^n$ gives information about the paths of length $n$ in the network. (The case where $n = 1$ corresponds to paths of length 1, i.e. the edges.) Specifically, given a pair $(s_1, s_2)$, the $(s_1, s_2)$'th entry $[\mathbf{A}^n]_{s_1, s_2}$ of the $n$th power $\mathbf{A}^n$ of the adjacency matrix $\mathbf{A}$ equals the sum over all paths from $s_1$ to $s_2$ with length exactly $n$ of the product of all of the weights of all of the edges in each path.

For example, if there were two paths of length exactly 3 from $s_1$ to $s_2$, and the weights of the edges in the first path were 1, $-4$, 2, and the weights of the edges in the second path were $-2$, 6, $-5$, then the $(s_1, s_2)$'th entry $[\mathbf{A}^3]_{s_1, s_2}$ of $\mathbf{A}^3$ would be $(1)(-4)(2) + (-2)(6)(-5) = -8 + 60 = 52$.

##### S3.4.2 Usefulness of this Formalism

The usefulness of encoding information about paths this way is uncontroversial in the case of an unsigned skeleton (i.e. an adjacency matrix consisting only of 1's and 0's), since then the entries count the number of paths of a given length in between any two nodes.

In the case of a non-negative weighted graph the usefulness of this notion is perhaps more controversial, e.g. one might wonder whether it makes more sense to take the sum of the weights along each path rather than the product. It is probably even more controversial in the case of networks with mixed-sign edge weights, since taking products over weights with different signs leads to difficult to predict "fluctuations" in the sign of the contribution from each path.

In particular, one might question whether the effect on signs given by multiplying the edges in the path is scientifically meaningful in a given context, or whether (instead of paths potentially cancelling one another out) edges within a path that have opposite signs should be allowed to cancel each other out.

##### S3.4.3 Usefulness for Microbial Ecology

I believe there is unlikely to be a universal answer. Instead, whether powers of adjacency matrices encode useful information about a mixed-sign network's paths likely depends on the context established by the specific scientific questions of interest. In the case of microbial ecology I believe that powers of adjacency matrices do make sense.

For example, consider a situation where strain $s_1$ strongly promotes the growth of strain $s_2$, and $s_2$ produces compounds toxic to strain $s_3$. Cf. figure S1, where $s_1$ could be the blue blob microbe, $s_2$ could be the pink spiral microbe, and $s_3$ could be the red rod microbe. I think it makes more sense for the resulting path from $s_1$ to $s_3$ to be given a strong negative weight, which corresponds only to the product of the weights of the individual edges, and *not* to the sum.

If one used the sum of the edge weights instead, one would instead "cancel" the effects of $s_1$ on $s_2$ and of $s_2$ on $s_3$. That makes no sense, because large numbers of $s_1$ would promote large numbers of $s_2$, which would in turn produce large amounts of the compound toxic to $s_3$. It seems clear that the net indirect effect of $s_1$ on $s_3$ is strong and negative, and that is reflected only through the product of the weights of the edges in the path, and not through the sum.

##### S3.4.4 Neumann Series

One might ask which length of path $n$ should be considered. A possible answer to this is "all of them", by using the Neumann series of the adjacency matrix $\mathbf{A}$. For a general matrix $\mathbf{M}$, its Neumann series is

$$\mathbf{M}+\mathbf{M}^2+\mathbf{M}^3+\cdots+\mathbf{M}^n+\cdots=(\mathbf{I}-\mathbf{M})^{-1}. \tag{S43}$$

This converges[11] if and only if the spectral radius of $\mathbf{M}$ is less than 1. Therefore it makes sense to think of the Neumann series as the matrix analogue of the geometric series for (real or complex) numbers.

The spectral radius lower bounds the Frobenius norm and any operator norm[12], so a sufficient (but not necessary) condition for the Neumann series of $\mathbf{M}$ to converge is if the value of one of these norms is less than 1 for $\mathbf{M}$. Therefore in general convergence of the corresponding Neumann series can be accomplished by rescaling $\mathbf{M}$ by a positive constant.

##### S3.4.5 DeltaCon Distance Overview

For both the unsigned and mixed-sign versions of DeltaCon distance, the general idea can be divided into two steps:

1. For each of the two networks, define a matrix which encodes the "path structure" of the network.
2. Define a distance between the two networks by evaluating a distance between the two matrices produced in the first step.

For the first step we form the Neumann series $\mathbf{S}$ of a suitable matrix $\mathbf{W}$. For the second step, we evaluate (a modified version of) the Matusita ("root Euclidean") distance of the two matrices $\mathbf{S}_1$, $\mathbf{S}_2$ constructed in the first step.

##### S3.4.6 DeltaCon Distance Definition

The definition of DeltaCon does not use the Neumann series of (a rescaled version of) the adjacency matrix $\mathbf{A}$, but instead uses the Neumann series of a matrix $\mathbf{W}$ based on $\mathbf{A}$. The formula (S44) for $\mathbf{W}$ in terms of $\mathbf{A}$ is derived from a linearized version of the belief propagation algorithm called FaBP(Koutra et al., 2011). The authors of (Koutra et al., 2016) state that nevertheless the intuition for using the Neumann series of the matrix $\mathbf{W}$ is still the same as the intuition described above for using the Neumann series of the adjacency matrix $\mathbf{A}$. The scaling constant $\varepsilon$ used below in the definition (S44) of this matrix $\mathbf{W}$ has the interpretation of encoding "the influence between neighboring nodes"(Koutra et al., 2016).

Combining the exact definition[13] from (Koutra et al., 2011) with the notation from (Koutra et al., 2016), I use the following for the definition of $\mathbf{W}$:

$$\mathbf{W} := \frac{1}{1-\varepsilon^2}(\varepsilon\mathbf{A}-\varepsilon^2\mathbf{D}). \tag{S44}$$

(The degree matrix $\mathbf{D}$ is a general notion defined in section S2.2.) The Neumann series $\mathbf{S}$ of $\mathbf{W}$, which encodes the "path structure" of the network, is called the "similarity matrix" in (Koutra et al., 2016) because each entry can be interpreted as measuring the "affinity" of two nodes in terms of the paths between them.

$$\mathbf{S} := (\mathbf{I}-\mathbf{W})^{-1} = \mathbf{I}+\mathbf{W}+\mathbf{W}^2+\dots. \tag{S45}$$

To ensure the definition of DeltaCon distance works we need to ensure the convergence of the Neumann series defining the similarity matrix $\mathbf{S}$. To guarantee the convergence of the Neumann series defining $\mathbf{S}$, we choose, for any given input $\mathbf{A}$, a corresponding value for $\varepsilon$ in (S44) that guarantees that the spectral radius of the resulting $\mathbf{W}$ defined in terms of $\mathbf{A}$ and $\varepsilon$ will be less than 1.

The normalization implicit in the definition of $\mathbf{W}$ (S44) and of the similarity matrix (S45) may help DeltaCon distance to avoid judging two highly different networks as being similar merely because both are sparse. Cf. again section S1.4, as well as the comments at the end of section S3.1.2. I am uncertain whether this is really true in practice, but it does seem plausible.

[11] This fact is not obvious, but appears to be a "folk theorem" such that it is difficult to find a "canonical" reference for the proof.
[12] See footnote 11, because the same comments apply here too.
[13] The definition of $\mathbf{W}$ from (Koutra et al., 2016) slightly differs and is not the original definition.

##### *S3.4.7 Unsigned Version*

The definition given in (Koutra et al., 2016) was intended to work for unsigned but weighted networks. Without being modified, it does not work for general networks with mixed-sign edge weights. I describe in detail the proposals made in (Koutra et al., 2016) for (1) the value of $\varepsilon$ in the definition (S44) of $\mathbf{W}$, and (2) the distance to apply to the similarity matrices $\mathbf{S_1}$ and $\mathbf{S}_2$. Along with using the original definition of $\mathbf{W}$ from (Koutra et al., 2011), these are the two aspects of the definition from (Koutra et al., 2016) that I changed in order to guarantee the new definition would work for all networks with mixed-sign edge weights.

**Step 1:** The authors of (Koutra et al., 2016) propose using the following for $\varepsilon$:

$$\varepsilon := \frac{1}{1+\max_i[\mathbf{D}]_{ii}}\,. \tag{S46}$$

However, they don't include an argument showing why this value (ostensibly) works. This is notable because the previous paper(Koutra et al., 2011) (i) only discussed unweighted (and unsigned) networks, (ii) used a different version of $\mathbf{W}$ (corresponding to (S44) above) than the definition used in (Koutra et al., 2016), and (iii) proposed using a slightly different value for $\varepsilon$ (for which a proof was provided).

**Step 2:** Given two adjacency matrices $\mathbf{A}_1$ and $\mathbf{A}_2$, and defining their corresponding similarity matrices $\mathbf{S}_1$ and $\mathbf{S}_2$ as in equation (S45), (Koutra et al., 2016) defines the (unsigned) DeltaCon distance to be the Matusita distance of $\mathbf{S}_1$ and $\mathbf{S}_2$:

$$d_{DC}(\mathcal{G}_1,\mathcal{G}_2) := \sqrt{\sum_{i,j}(\sqrt{[\mathbf{S}_1]_{ij}} - \sqrt{[\mathbf{S_2}]_{ij}})^2}\,. \tag{S47}$$

The authors of (Koutra et al., 2016) suggest several reasons for using Matusita distance instead of e.g. Frobenius distance, but one important reason they mention is "boosting" small entries in $[0,1]$ (given that $\sqrt{x} \geq x$ for all $x \in [0,1]$).

##### *S3.4.8 Mixed-Sign Version*

To modify the definition from (Koutra et al., 2016) to apply to networks with mixed-sign edge weights, I made the following two changes.

**Step 1:** I showed that a sufficient condition for a value of $\varepsilon$ to work with any network with mixed-sign edge weights is for $\varepsilon < \frac{1}{1+\|\mathbf{A}\|_\infty}$ (see Lemma S7.5 from section S7.2). I also showed that this bound is sharp (see Lemma S7.6 from section S7.2), in the sense that there exist adjacency matrices $\mathbf{A}$ of networks with mixed-sign edge weights for which choosing $\varepsilon := \frac{1}{1+\|\mathbf{A}\|_\infty}$ leads to a $\mathbf{W}$ with spectral radius exactly equal to 1. While such problematic adjacency matrices do not appear to be generic, for safety I chose to use instead:

$$\varepsilon := \frac{1}{2+\|\mathbf{A}\|_\infty}\,. \tag{S48}$$

In practice, even when $\varepsilon := \frac{1}{1+\|\mathbf{A}\|_\infty}$ technically works for a given network, if the spectral radius of the resulting $\mathbf{W}$ is still very close to 1, the DeltaCon distance appears to be poorly behaved, e.g. by producing unreasonably large values (data not shown). Using the more conservative definition (S48) of $\varepsilon$, the resulting $\mathbf{W}$ matrices had spectral radii bounded further away from 1, making the resulting values of the DeltaCon distance more stable than they were before.

**Step 2:** In equation (S47), I replaced $\sqrt{[\mathbf{S}]_{ij}}$ with $\operatorname{sign}([\mathbf{S}]_{ij}) \cdot \sqrt{|[\mathbf{S}]_{ij}|}$:

$$d^{\pm}_{DC}(\mathcal{G}_1,\mathcal{G}_2) := \sqrt{\sum_{i,j}\left(\operatorname{sign}([\mathbf{S}_1]_{ij}) \cdot \sqrt{|[\mathbf{S}_1]_{ij}|} - \operatorname{sign}([\mathbf{S}_2]_{ij}) \cdot \sqrt{|[\mathbf{S_2}]_{ij}|}\right)^2}\,. \tag{S49}$$

The choice to use $\operatorname{sign}([\mathbf{S}]_{ij}) \cdot \sqrt{|[\mathbf{S}]_{ij}|}$ instead of $\sqrt{[\mathbf{S}]_{ij}}$ appears to be necessary for at least two reasons. First, it ensures that the distance is defined (by not assuming that the entries of $\mathbf{S}$ are necessarily non-negative in the more general case of mixed-sign edge weights). Second, it also guarantees that the double penalization principle (cf. section **??**) is satisfied. At the same time, the new definition (S49) also retains the same "boosting" property that helped motivate definition (S47), because $\sqrt{|x|} \geq |x|$ for all $x \in [-1,1]$. So nothing important appears to be lost because of the change, as reflected perhaps in how (S49) reduces to (S47) in the case that the entries of $\mathbf{S}_1$ and $\mathbf{S}_2$ are non-negative.

### S4 METHODS

$1,000$ random networks were generated (section S4.1) and classified according to which sign had the larger number of edges (section S4.2). Three adversarial attacks were applied to each of the networks (section S4.3). The values of the mixed-sign network comparison methods between the original networks and their attacked versions were compared with the corresponding values for variants which do not use the entirety of the mixed-sign network structure or which do not obey the Double Penalization Principle (section S4.4). Side-by-side violin plots of the results were generated using Matplotlib (Hunter, 2007) version 3.4.1 and Seaborn Waskom (2021) version 0.11.1. Complete implementation details can be found in the code at the relevant GitLab repository. See https://gitlab.com/krinsman/mixed-sign-networks.

#### S4.1 Generating Random Networks

$1,000$ random networks were independently generated as follows:

- A hyperparameter $p \in (0,1)$ was selected from the uniform distribution.
- $100^2$ random entries of an adjacency matrix were independently generated according to a $\text{Beta}(2p, 2(1-p))$ distribution. (Considered as a Dirichlet distribution, this has the same concentration parameter as the uniform distribution.)
- 0.5 was subtracted from all entries, and then all entries were multiplied by 2, with the effect that their support changed from $(0,1)$ to $(-1,1)$.
- Each of the entries in the adjacency matrix was set to zero independently and with probability $\frac{1}{2}$ (this means the random networks are Erdos-Renyi).

#### S4.2 Predominant and Non-Predominant Sign Edges

For each network, if a majority of the nonzero entries were positive, $+$ was the "predominant sign" and $-$ the "non-predominant sign". If a majority of the nonzero entries were negative, $-$ was the "predominant sign" and $+$ the "non-predominant sign". Results looking at "predominant sign edges" consider only the subnetworks (using the definitions from section S2.4) corresponding to the "predominant sign" for any given networks. Similarly for results looking at "non-predominant sign edges". Results were split according to the "predominant sign" and "non-predominant sign" for each network, rather than always grouping the positive subnetworks together and the negative subnetworks together, because the corresponding distributions usually depended only on whether the subnetwork corresponded to the numerically predominant sign for that network, and not to what the particular sign was.

#### S4.3 Definitions of the Adversarial Attacks

Three types of adversarial attack were applied to each of the random networks. The corresponding values of similiarity/dissimilarity were then computed between each original random network and their attacked counterparts.

The three types were as follows.

##### *S4.3.1 Shift Attack*

This adversarial attack affected both the signs and magnitudes of the edges of the random network. The effect on the signs was the same as the sign flip attack. First, the predominant sign of the network was identified. If the predominant sign was positive, then twice the maximum value of any edge in the network was subtracted from all nonzero edges, forcing all edges to be negative (the non-predominant sign). If the predominant sign was negative, then twice the absolute value of the minimum value of any edge in the network was added to all nonzero edges, forcing all edges to be positive (the non-predominant sign). In both cases the values of all nonzero edges were shifted by a constant (whence the name), leaving their relative ordering unaffected.

##### *S4.3.2 Magnitude Swap Attack*

This adversarial attack affected the magnitudes of the edges of the random network, but not the signs. The relative ordering of all magnitudes of all nonzero edges was computed. Then the edge with the largest magnitude had its magnitude swapped with the magnitude of the edge with the smallest magnitude, the edge with the second largest magnitude had its magnitude swapped with the magnitude of the edge with the second smallest magnitude, and so on. For all $n$ (less than or equal to the number of nonzero edges), the edge with the $n$th largest magnitude had its magnitude swapped with the magnitude of the edge with the $n$th smallest magnitude.

###### S4.3.3 Sign Flip Attack

This adversarial attack affected the signs of the edges of the random network, but not the magnitudes. First, the predominant sign of the network was identified. Then all of the edges corresponding to the predominant sign had their sign flipped (i.e. multiplied by $-1$), while all edges corresponding to the non-predominant sign were unchanged.

##### S4.4 Definitions of Variants

Mixed-sign network comparison methods were compared against related comparison methods that only considered subsets of the features of the networks. Section S4.4.1 explains the definitions of the variants used for (entrywise $L_1$) relative error. Section S4.4.2 explains the definitions of the variants used for Spearman correlation. Section S4.4.3 explains the definitions of the variants used for Jaccard similarity. Section S4.4.4 explains the definitions of the variants used for DeltaCon distance.

###### S4.4.1 Relative Error

First I compared the values of relative error with those values resulting from considering (i) only the magnitudes of the original networks or (ii) only the signs of the original networks. I then compared the distribution of relative error values for the whole network with the corresponding values for the subnetworks corresponding to (i) the predominant sign edges only or (ii) the non-predominant sign edges only.

Whenever computing the relative error of two (sub)networks, the magnitudes of the edges were first normalized by the value of the entrywise $L_1$ norm of the (sub)networks. Cf. section S3.1.4. This served two purposes. First, it ensured that the distribution of values was confined to $[0,2]$ and thus made the results easier to visualize. Second, it corresponds to what it was done in preliminary work, where only the relative but not absolute sizes of estimates are of interest. The convex combination relationship mentioned in section S3.1.3 is therefore no longer directly relevant. Thus the most accurate interpretation of results is somewhat obscured.

"Magnitudes only" refers to the relative error between the "magnitude skeletons" $\mathcal{M}(\mathcal{G}_1)$, $\mathcal{M}(\mathcal{G}_2)$ of the original networks $\mathcal{G}_1, \mathcal{G}_2$. "Signs only" refers to the relative error between the "signed skeletons" $\mathcal{S}^{\pm}(\mathcal{G}_1), \mathcal{S}^{\pm}(\mathcal{G}_2)$ of the original networks $\mathcal{G}_1, \mathcal{G}_2$. (See section S2.3 for skeleton definitions.)

###### S4.4.2 Spearman Correlation

First I compared the values of mixed-sign Spearman correlation with those values resulting from (i) using the raw Spearman correlation, considering (ii) only the magnitudes of the original networks, or (iii) only the signs of the original networks. I then compared the distribution of mixed-sign Spearman correlation values for the whole network with the corresponding (raw) Spearman correlation values for the subnetworks corresponding to (i) the predominant sign edges only or (ii) the non-predominant sign edges only.

"Raw" Spearman correlation refers to the sparsity-adjusted Spearman correlation of the (unaltered) original networks $\mathcal{G}_1, \mathcal{G}_2$. "Magnitudes only" refers to the sparsity-adjusted Spearman correlation of the "magnitude skeletons" $\mathcal{M}(\mathcal{G}_1), \mathcal{M}(\mathcal{G}_2)$ of the original networks $\mathcal{G}_1, \mathcal{G}_2$. Similarly "signs only" refers to the sparsity-adjusted Spearman correlation of the "signed skeletons" $\mathcal{S}^{\pm}(\mathcal{G}_1), \mathcal{S}^{\pm}(\mathcal{G}_2)$ of the original networks $\mathcal{G}_1, \mathcal{G}_2$. (See section S2.3 for skeleton definitions.)

As mentioned already in section S3.2.5, only looking at the subset of edges which are nonzero in at least one of the networks (i.e. not considering edges missing from both networks) avoids considering two very different networks as similar due only to them both being highly sparse.

Also note that when one of the compared vectors of edge values was constant (and thus its rank vector had 0 variance), the Spearman correlation herein is defined by convention to be 0 (since the ranks from one vector have zero predictive value for predicting the ranks of the other vector).

###### S4.4.3 Jaccard Similarity

First I compared the values of mixed-sign unweighted Jaccard similarity and mixed-sign weighted Jaccard similarity with the values resulting from considering only (i) the presence/absence of edges or (ii) the magnitudes of edges. I then compared the mixed-sign unweighted Jaccard similarity values with the

Jaccard similarity values for the subnetworks corresponding to (i) the predominant sign edges only or (ii) the non-predominant sign edges only, and then analogously also for the mixed-sign weighted Jaccard similarity.

"Presence/Absence Only" refers to the weighted Jaccard similarity of the "unsigned skeletons" $\mathcal{S}(\mathcal{G}_1), \mathcal{S}(\mathcal{G}_2)$ of the original networks $\mathcal{G}_1, \mathcal{G}_2$ (or equivalently the unweighted Jaccard similarity of the "magnitude skeletons" $\mathcal{M}(\mathcal{G}_1), \mathcal{M}(\mathcal{G}_2)$ of the original networks $\mathcal{G}_1, \mathcal{G}_2$). "Magnitudes only" refers to the weighted Jaccard similarity of the "magnitude skeletons" $\mathcal{M}(\mathcal{G}_1), \mathcal{M}(\mathcal{G}_2)$ of the original networks $\mathcal{G}_1, \mathcal{G}_2$. (See section S2.3 for skeleton definitions.)

Variants called "Signs Only" for relative error or Spearman correlation effectively correspond to the mixed-sign unweighted Jaccard similarity (which again is equivalent to the mixed-sign weighted Jaccard similarity of the "signed skeletons" of the original networks). More variants are considered than for relative error or Spearman correlation because the unweighted versions of those (their "Signs only" variants) are not as interesting as they are for Jaccard similarity.

###### *S4.4.4 DeltaCon Distance*

First I compared the values of mixed-sign unweighted DeltaCon distance and mixed-sign weighted DeltaCon distance with the values resulting from considering only (i) the presence/absence of edges or (ii) the magnitudes of edges. I then compared the mixed-sign unweighted DeltaCon distance values with the DeltaCon distance values for the subnetworks corresponding to (i) the predominant sign edges only or (ii) the non-predominant sign edges only, and then analogously also for the mixed-sign weighted DeltaCon distance.

"Presence/Absence Only" refers to the DeltaCon distance of the "unsigned skeletons" $\mathcal{S}(\mathcal{G}_1), \mathcal{S}(\mathcal{G}_2)$ of the original networks $\mathcal{G}_1, \mathcal{G}_2$, analogous to Jaccard similarity. "Magnitudes Only" refers to the DeltaCon distance of the "magnitude skeletons" $\mathcal{M}(\mathcal{G}_1), \mathcal{M}(\mathcal{G}_2)$ of the original networks $\mathcal{G}_1, \mathcal{G}_2$. Variants called "Signs Only" elsewhere effectively correspond to the mixed-sign *unweighted* DeltaCon distance, which by definition equals the mixed-sign weighted DeltaCon distance of the "signed skeletons" $\mathcal{S}^{\pm}(\mathcal{G}_1), \mathcal{S}^{\pm}(\mathcal{G}_2)$ of the original networks $\mathcal{G}_1, \mathcal{G}_2$. (See section S2.3 for skeleton definitions.)

### S5 RESULTS

Section S5.1 discusses the behavior of variants of relative error for all three adversarial attacks. Section S5.2 discusses the behavior of variants of Spearman correlation for all three adversarial attacks. Section S5.3 discusses the behavior of variants of Jaccard similarity for all three adversarial attacks. Section S5.4 discusses the behavior of variants of DeltaCon distance for all three adversarial attacks.

#### S5.1 Relative Error

Section S5.1.1 overviews the results for the shift attack, explaining which variants were sensitive to which adversarial attacks. Section S5.1.2 explains the indirect evidence for the convex combination property seen in the results. (Note that nevertheless the convex combination property is proven and doesn't actually require empirical evidence to be substantiated.)

###### *S5.1.1 Sensitivity to Adversarial Attacks*

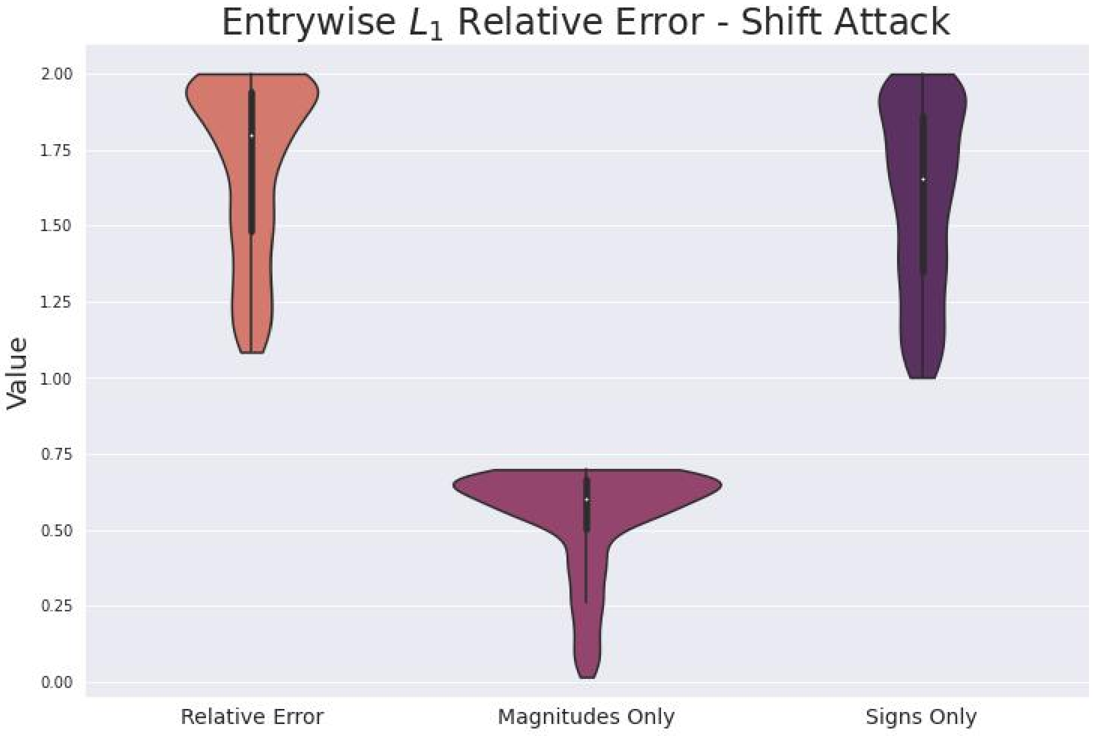

**Supplementary Figure S5**

For the shift attack in figure S5, the wider flare at the top of the distribution for relative error compared to signs only suggests that relative error is able to combine both sign and magnitude information to be more sensitive to adversarial attacks than considering either magnitudes only or signs only. Despite the changes in magnitude the accompany the shift attack, in figure S5 we see that considering magnitudes only still leads to substantially less sensitivity.

###### *S5.1.2 Effects of Convex Combination Decomposition Property*

As mentioned previously in section S4.4.1, that subnetworks were normalized before computing the relative error (cf. section S3.1.4) means the interpretation as a convex combination as in equation (S9) does not apply. Nevertheless, we still do see in figures S8, S6, and S7 results for the relative error which are weakly intermediate between the relative errors for the predominant sign edges and the non-predominant sign edges. Note that for the shift attack in figure S8 and the sign flip attack in figure S7, the fact that the relative errors are all exactly 1 is because the corresponding subnetwork has 0 edges in the attacked versions of the networks.

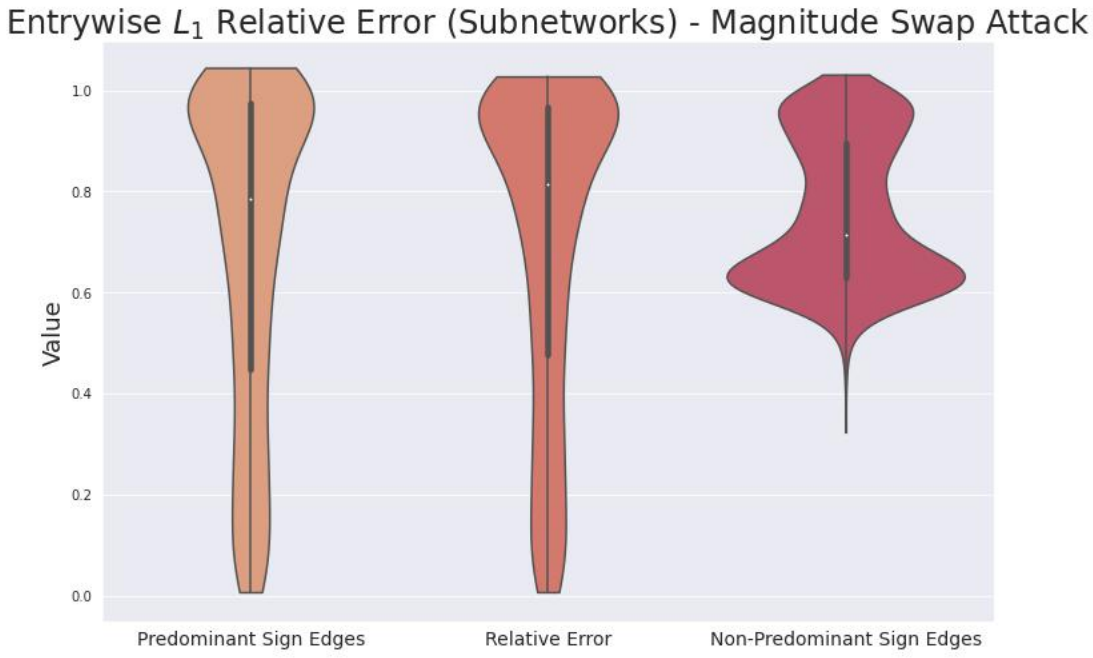

**Supplementary Figure S6**

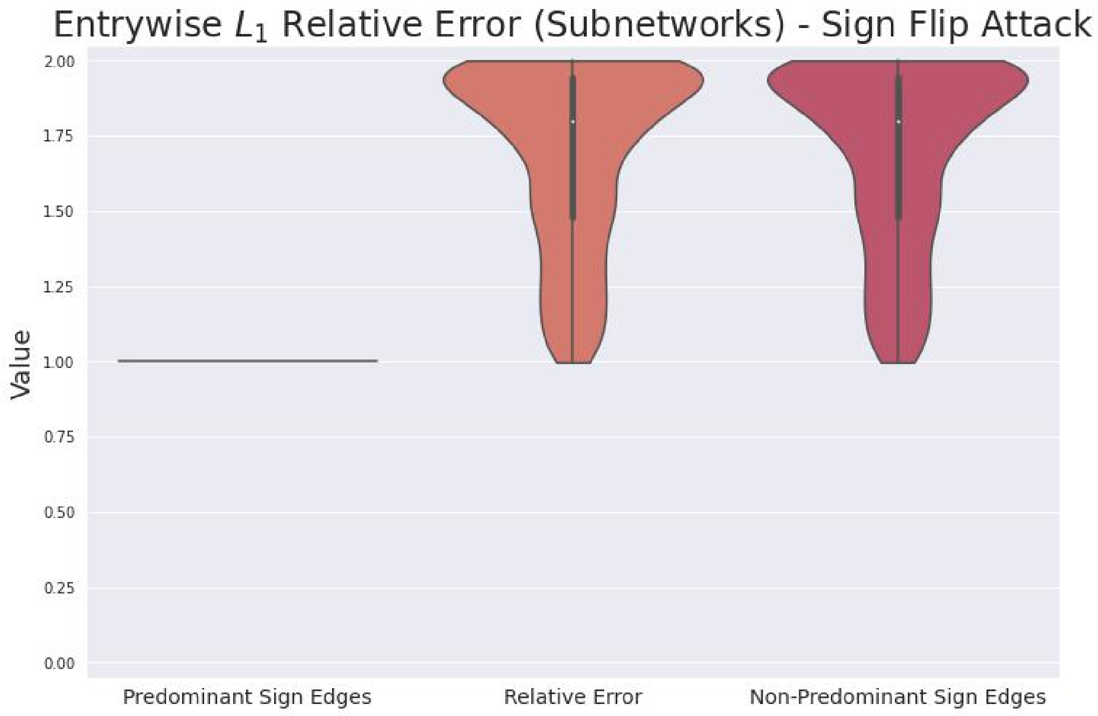

**Supplementary Figure S7**

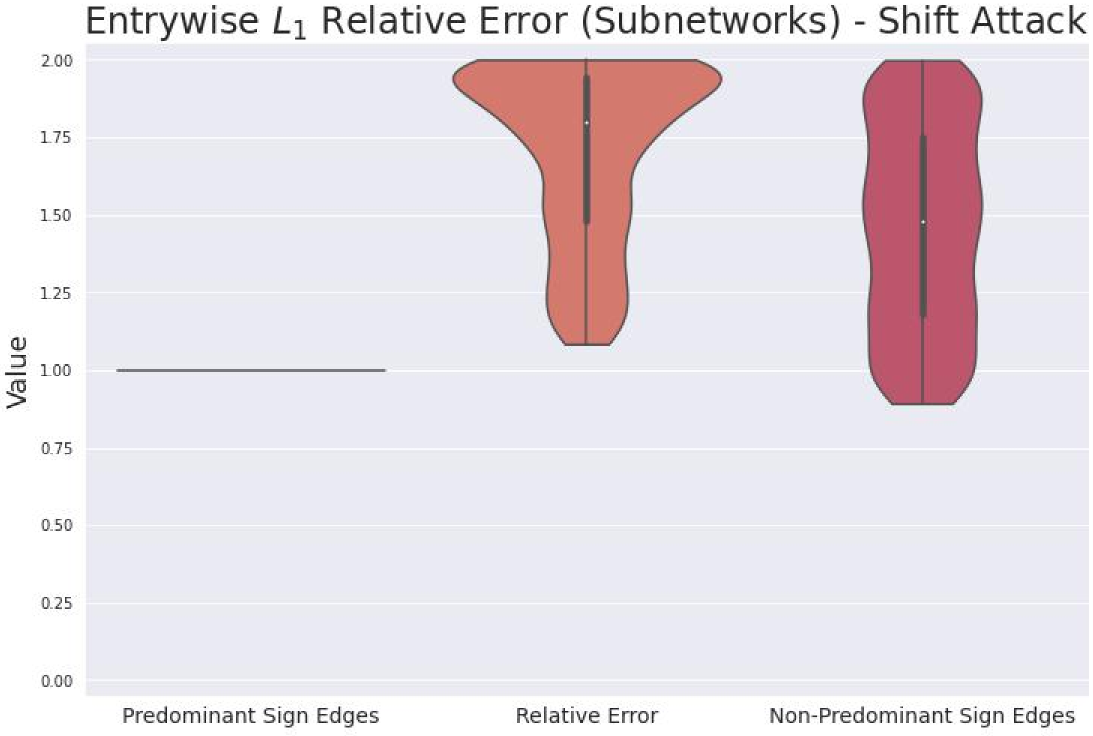

**Supplementary Figure S8**

#### S5.2 Spearman Correlation

Section S5.2.1 explains the evidence from the results that the mixed-sign Spearman not only directly combines the information from the positive and negative edges.

##### *S5.2.1 Combining Information*

The mixed-sign Spearman correlation is the only variant that is not oblivious to any of the adversarial attacks, and which always reports values depending on the particular structure of the underlying network. We also see from figures S11, S9, and S10 that the mixed-sign Spearman reports values intermediate between those for the predominant sign and non-predominant sign subnetworks, corresponding to the subconvex combination property mentioned in section S3.2.6 and demonstrating double penalization. The mixed-sign Spearman directly combines information from the positive and negative edges.

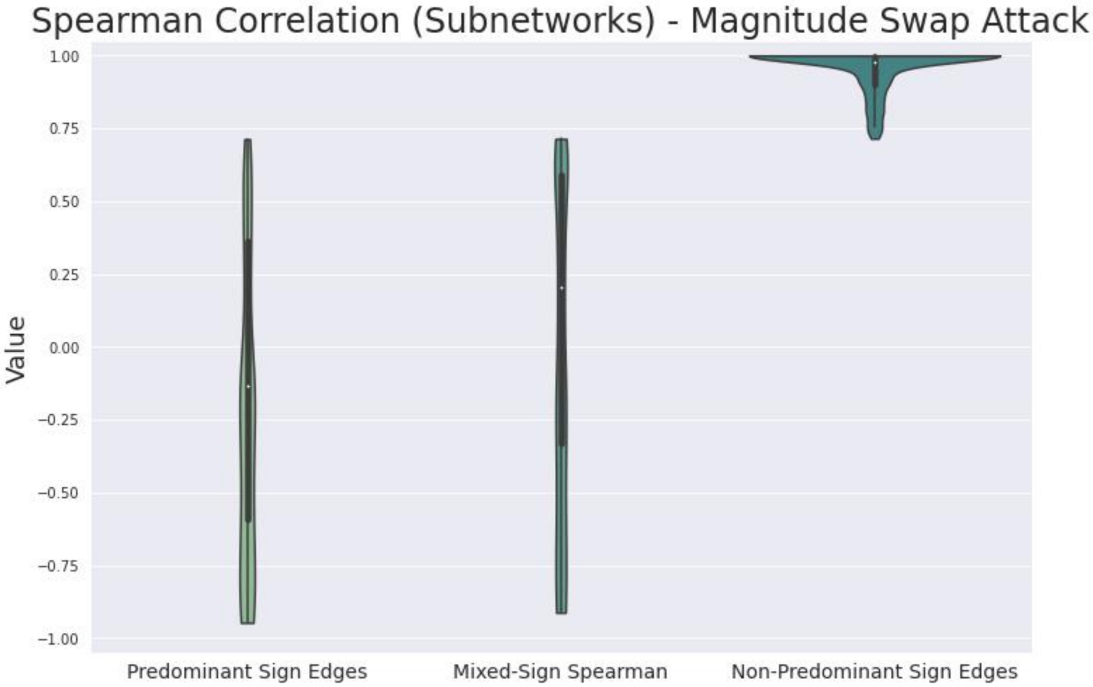

**Supplementary Figure S9**

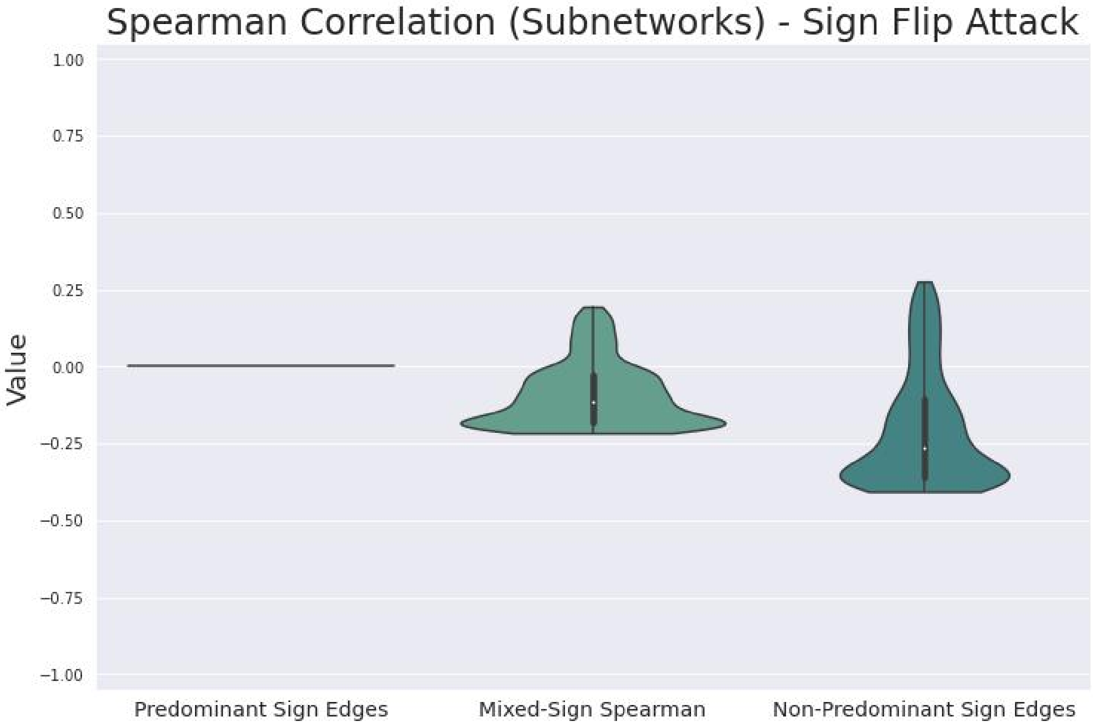

**Supplementary Figure S10**

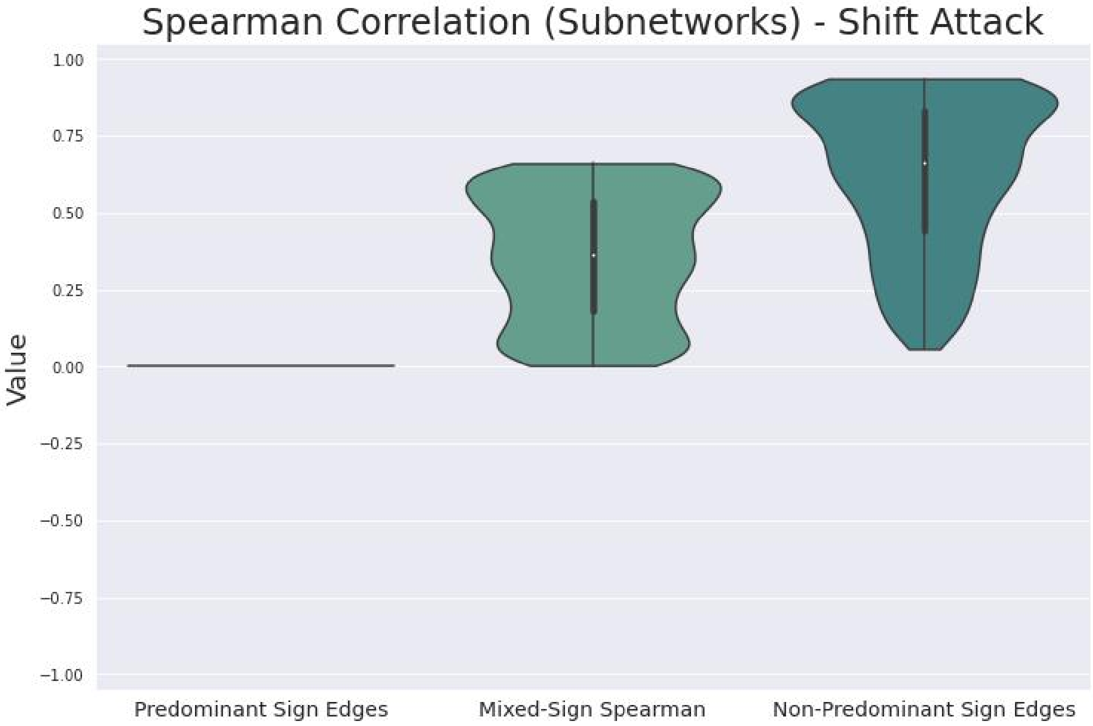

**Supplementary Figure S11**

#### S5.3 Jaccard Similarity

Section S5.3.2 explains how the convex combination property satisfied by both the mixed-sign unweighted and mixed-sign weighted Jaccard similarity manifests itself in the results.

##### *S5.3.1 Mixed-Sign Weighted and Magnitudes Only Coincide for Magnitude Swap*

The distribution of values appears largely identical between the mixed-sign weighted Jaccard similarity and the magnitudes only variant for the magnitude swap attack (figure S12). I am not entirely sure why that is the case; it does not seem to obviously follow as a consequence of the definition (S35). However, that the distribution of values for the Jaccard similarity for the predominant sign edges also has largely the same shape for the magnitude swap attack (cf. figure S16) is probably an important clue.

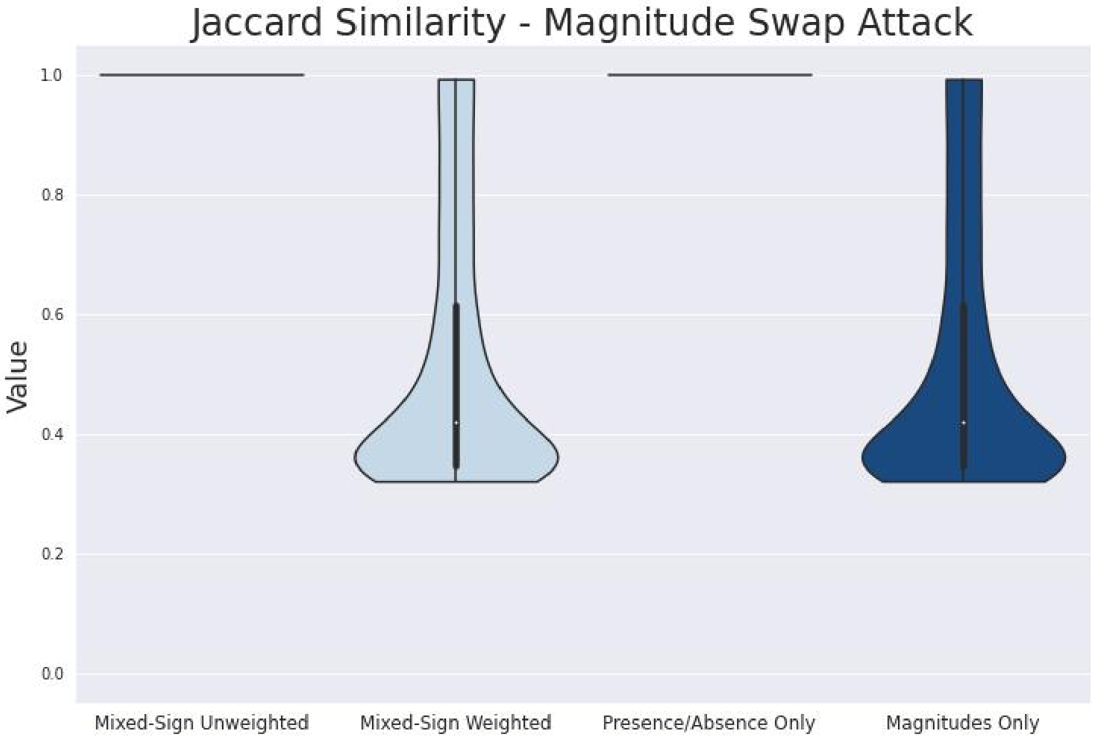

**Supplementary Figure S12**

###### *S5.3.2 Effects of Convex Combination Decomposition Property*

Figures S15, S13, and S14 correctly suggest how the mixed-sign unweighted Jaccard similarity is a convex combination of the unweighted Jaccard similarities for the positive and negative subnetworks.

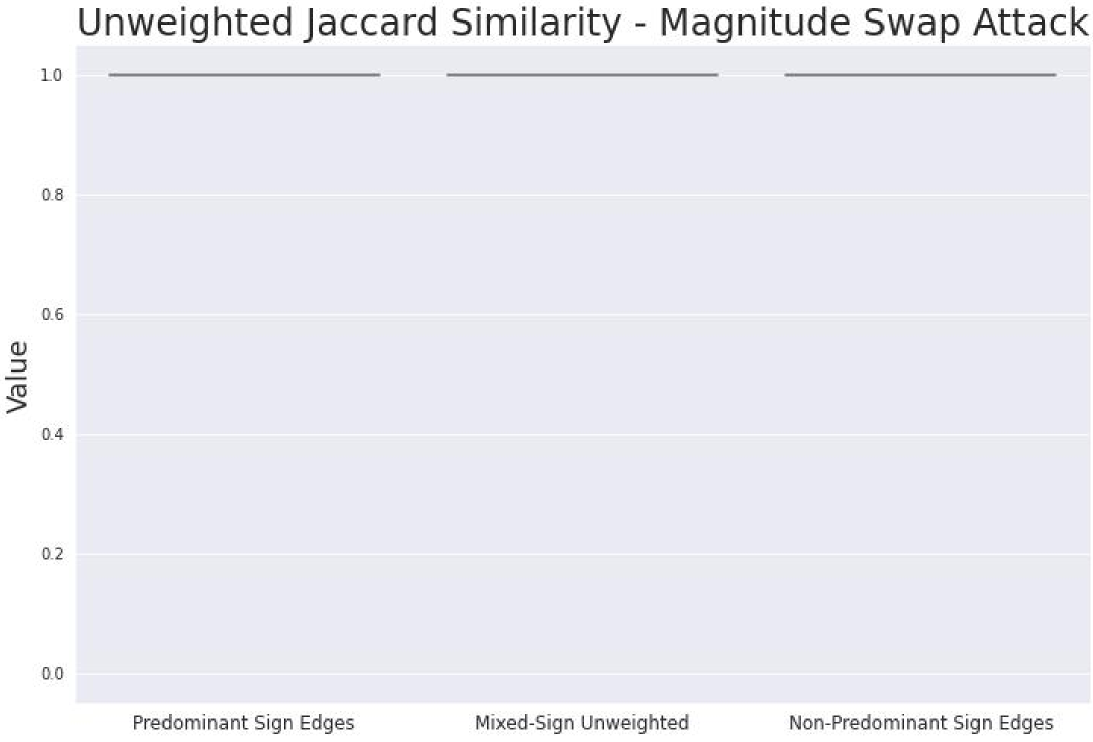

**Supplementary Figure S13**

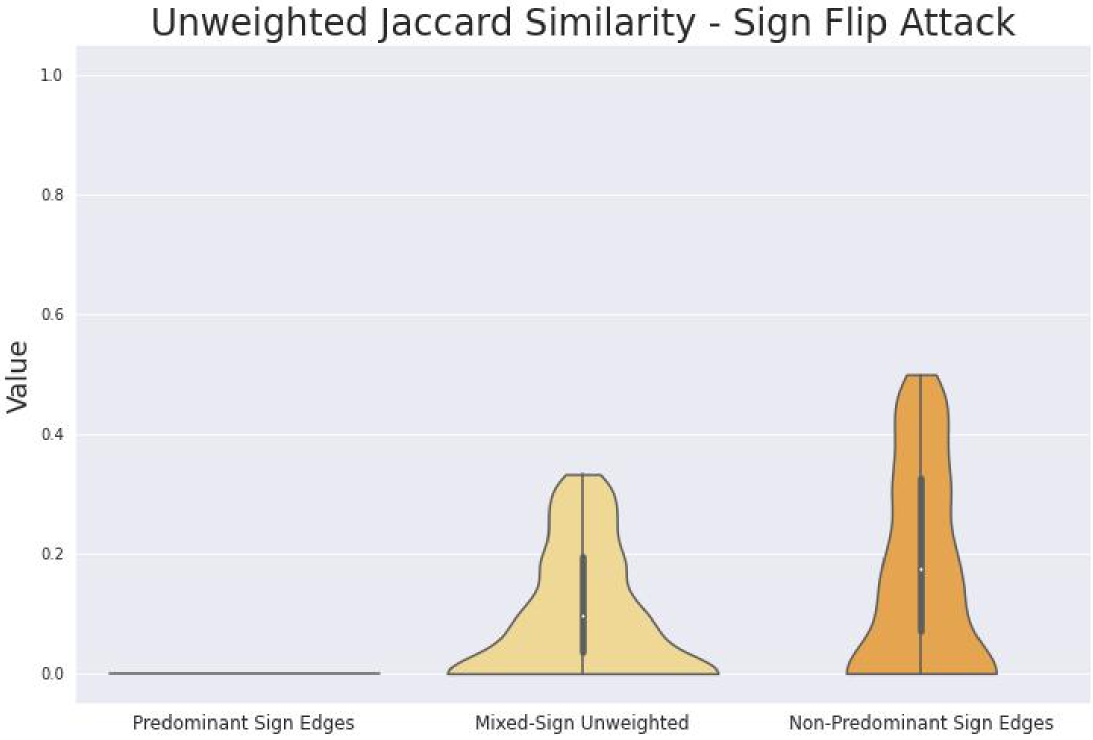

**Supplementary Figure S14**

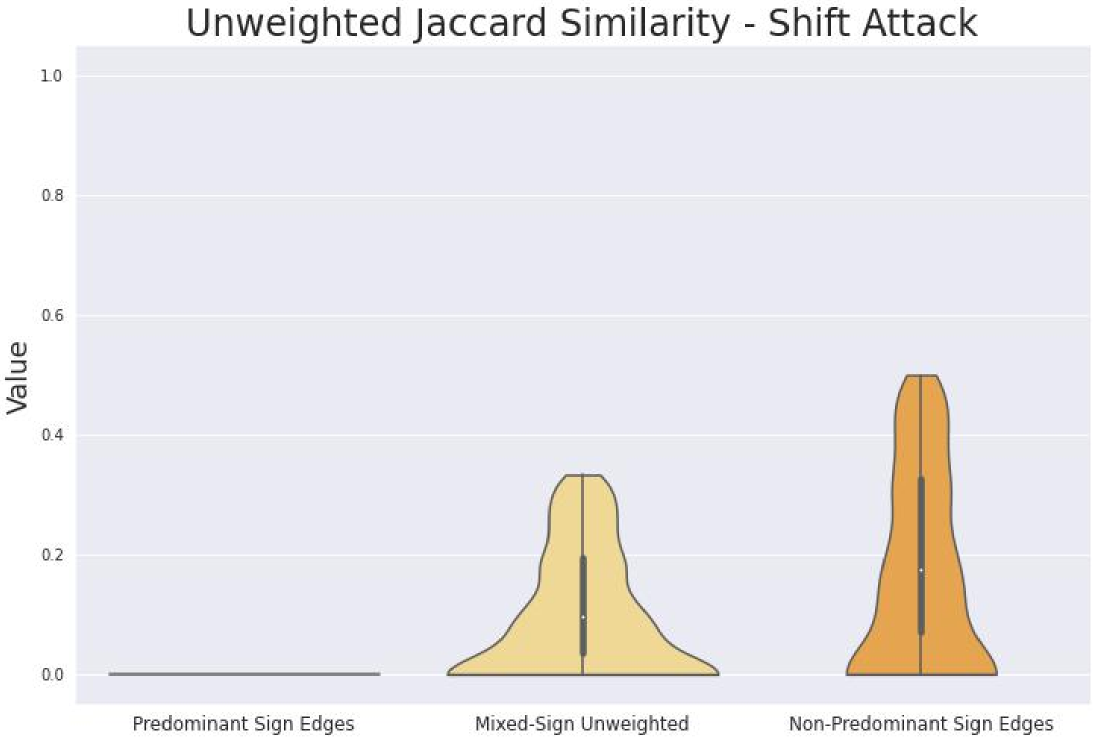

**Supplementary Figure S15**

Similarly, figures S18, S16, and S17 also correctly suggest how the mixed-sign weighted Jaccard similarity is a convex combination of the weighted Jaccard similarities for the positive and negative subnetworks.

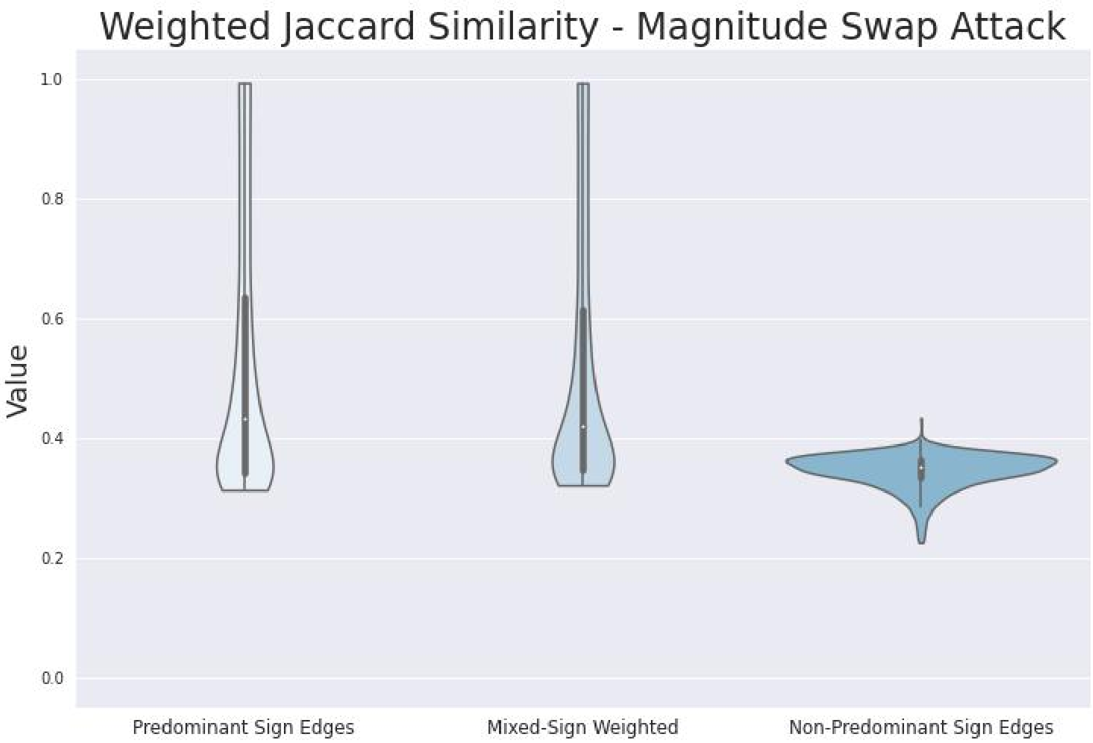

**Supplementary Figure S16**

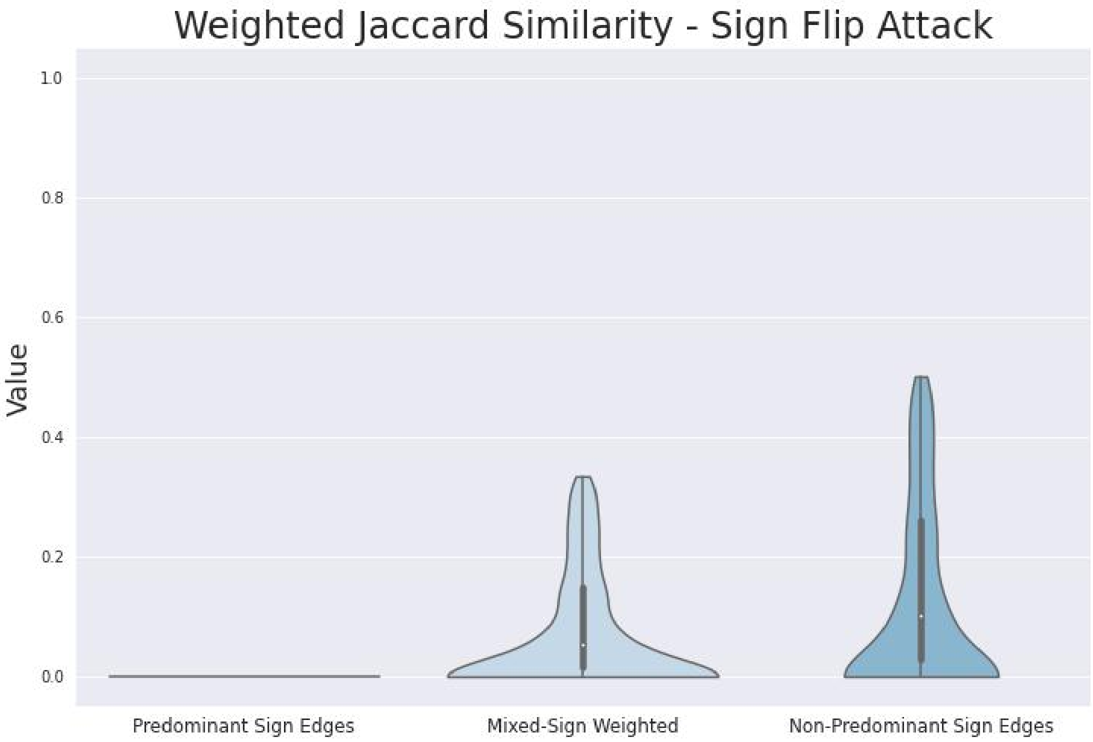

**Supplementary Figure S17**

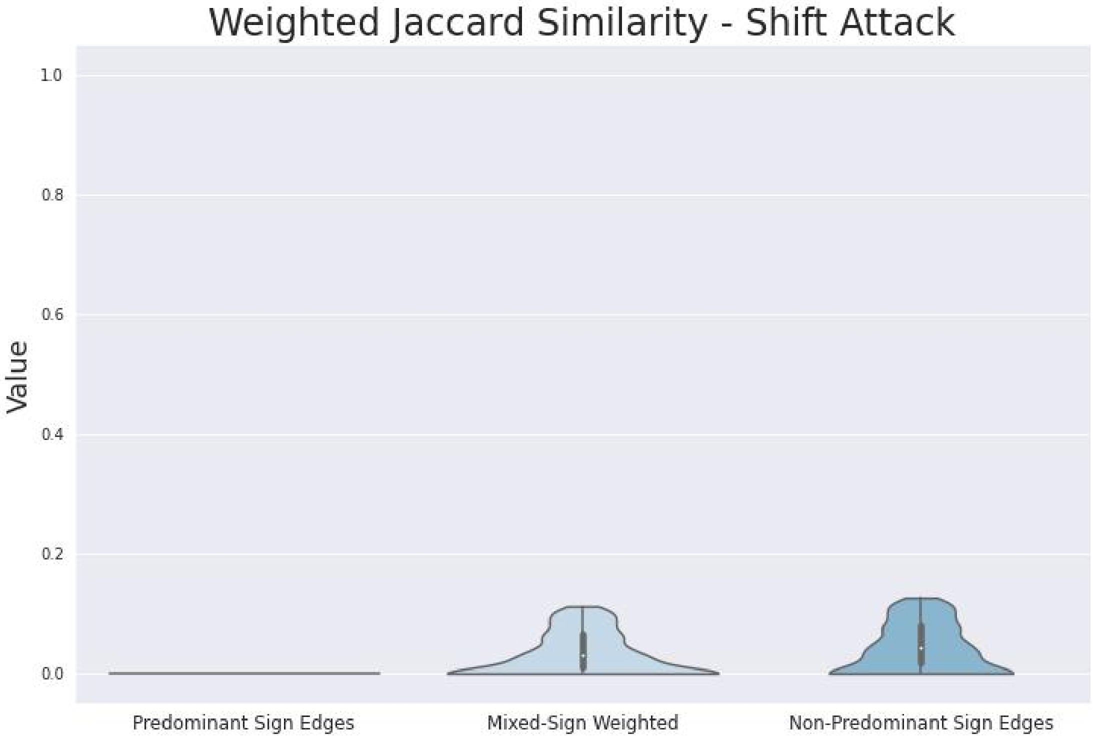

**Supplementary Figure S18**

Thus both mixed-sign versions of Jaccard similarity incorporate information from both subnetworks in a reasonable fashion.

#### S5.4 DeltaCon Distance

Section S5.4.1 explains an important caveat when comparing the results for mixed-sign unweighted and mixed-sign weighted DeltaCon distance. Section S5.4.2 gives a "sanity check" that demonstrates the results are sensible and in line with expectations. Section S5.4.3 argues that the subnetwork results show that the double penalization principle is important of DeltaCon in spite of it not satisfying the convex combination property (section S1.3).

##### *S5.4.1 Mixed-Sign Weighted May Actually Not Be Less Sensitive than Mixed-Sign Unweighted*

That the values of the mixed-sign unweighted DeltaCon distance trend slightly larger than those for the mixed-sign weighted DeltaCon distance in figures 13 and 12 appears to be an artifact of the magnitudes all being less than or equal to 1. (Thus the "magnitudes" of the signed skeleton are in general greater, and mixed-sign unweighted DeltaCon is equivalent to mixed-sign weighted DeltaCon applied to the signed skeletons of the original networks.)

##### *S5.4.2 Mixed-Sign Weighted Picks up Changes in Magnitude*

Note that the distributions of values look the same between figures S21 and S20, reflecting the fact that the shift and sign flip attacks have the same effect on the signs of the edges, and that the mixed-sign unweighted DeltaCon is only sensitive to changes in sign.

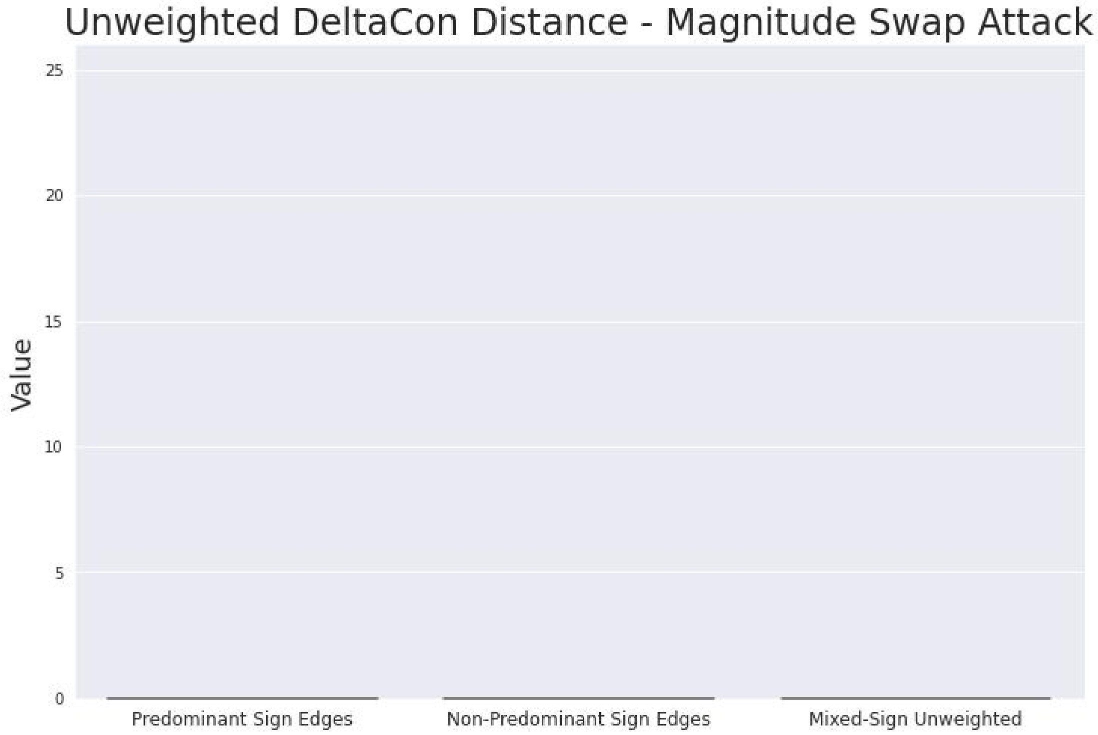

**Supplementary Figure S19**

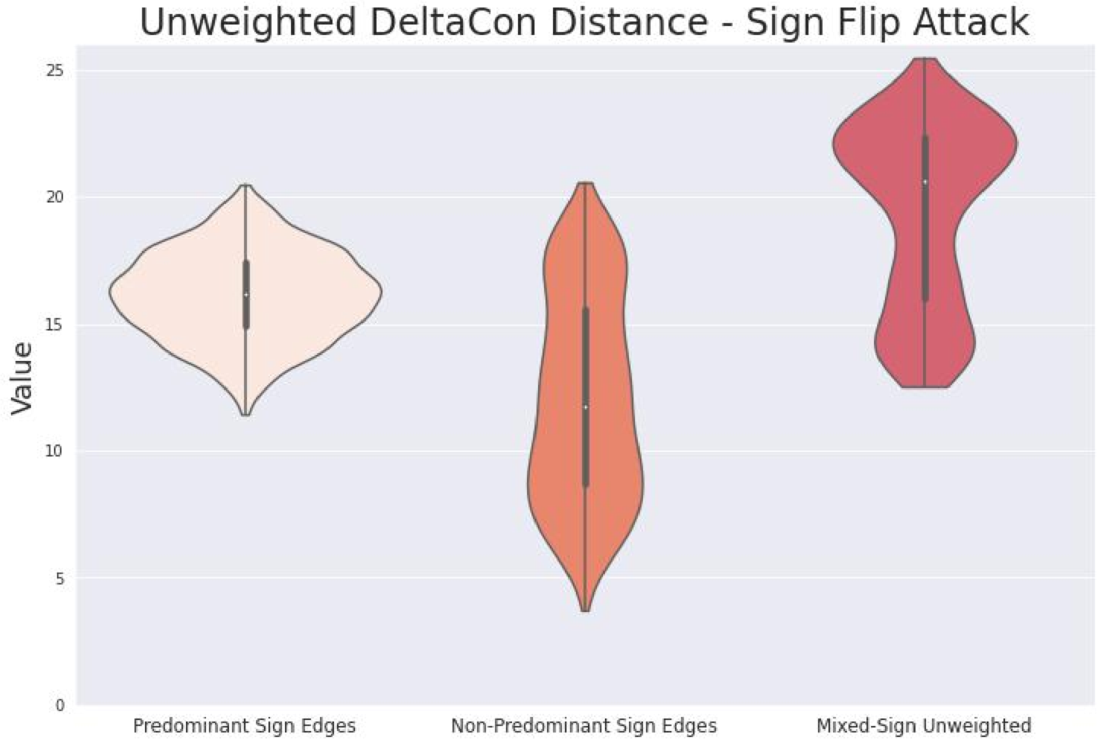

**Supplementary Figure S20**

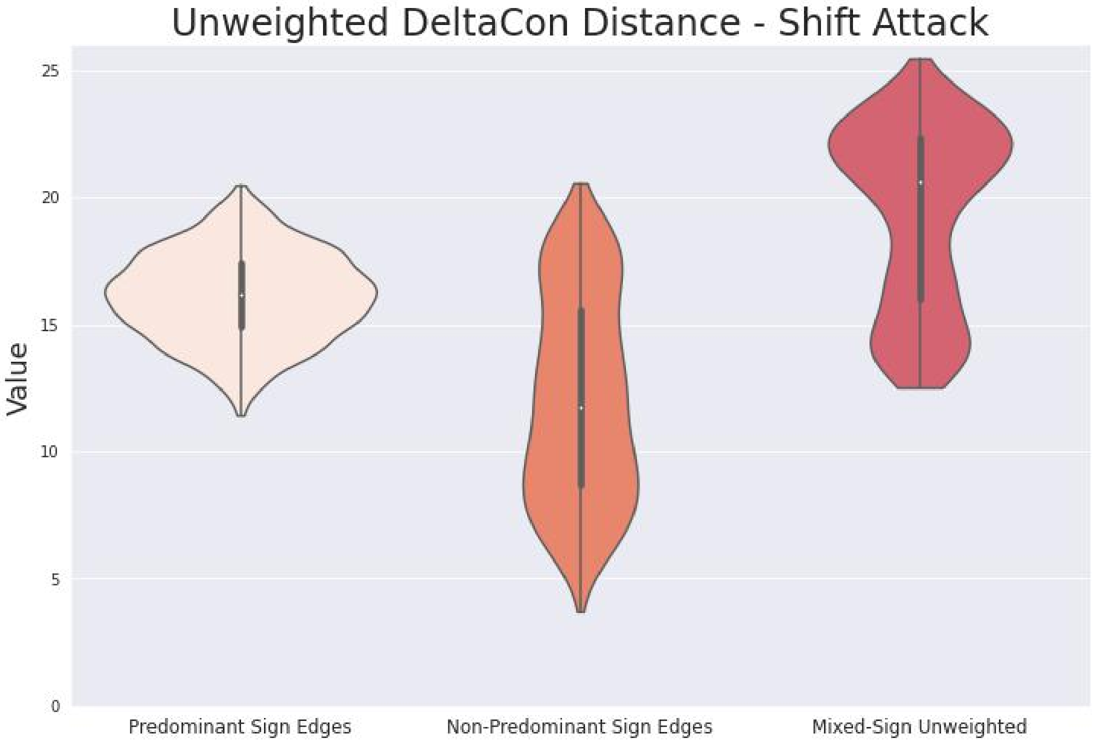

**Supplementary Figure S21**

In contrast, the distributions of values look different between figures S24 and S23, reflecting that the shift attack affects magnitudes (and the sign flip attack does not), and also demonstrating that the mixed-sign weighted version of DeltaCon is truly sensitive to changes in both sign and magnitude.

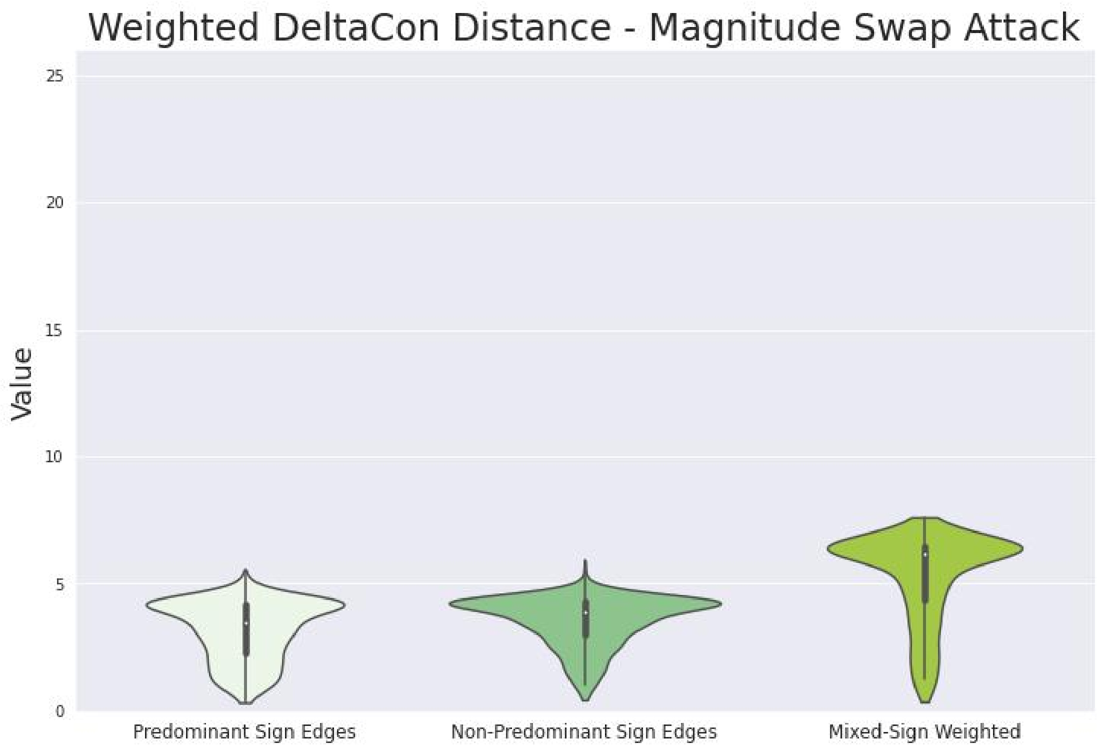

**Supplementary Figure S22**

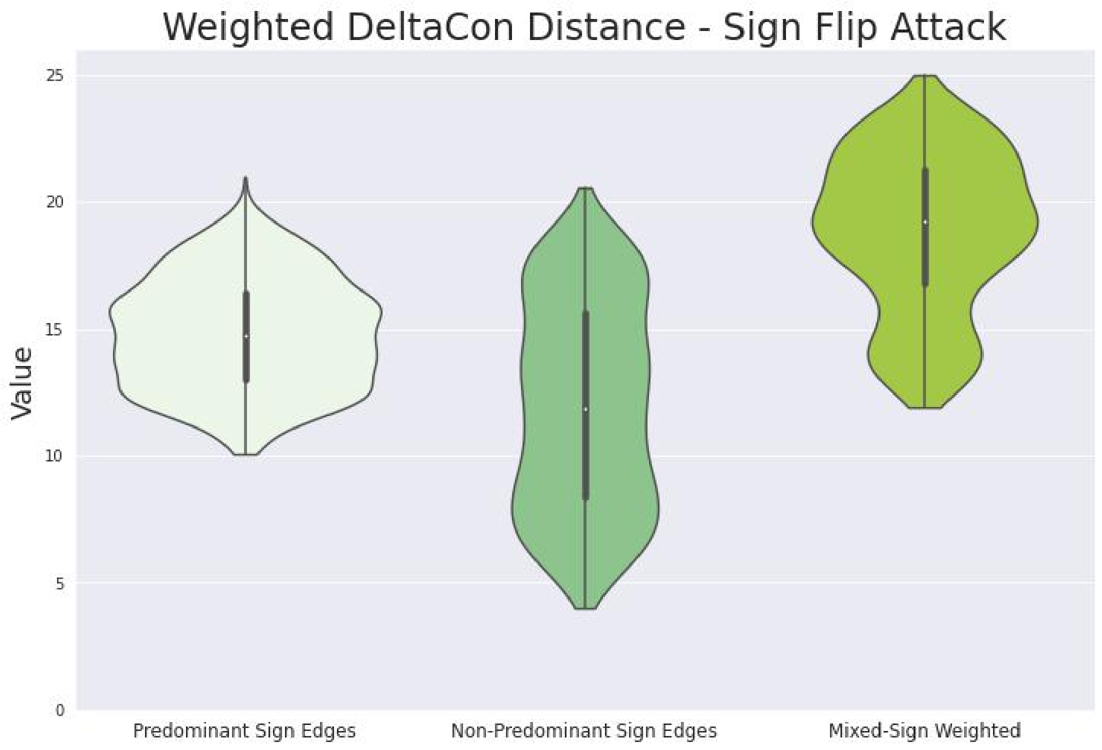

**Supplementary Figure S23**

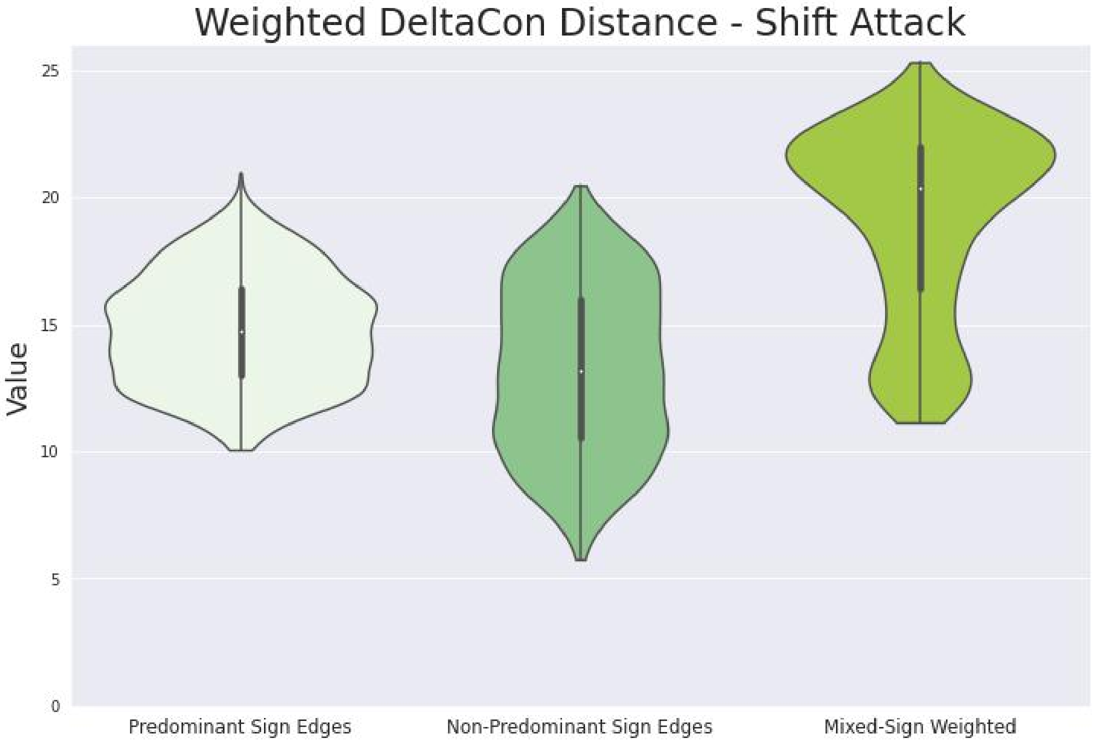

**Supplementary Figure S24**

###### *S5.4.3 Double Penalization Explains Improved Performance Compared to Subnetworks*

Figures S21 and S20 show that the mixed-sign unweighted version is more sensitive than the changes to either the predominant or non-predominant sign subnetworks alone when the adversarial attack changes sign, which is further evidence that double penalization is occurring. Similarly, figures S24, S22, S23 also show that the mixed-sign weighted DeltaCon is more sensitive to all of the adversarial attacks than either the predominant or non-predominant sign subnetworks. For both unweighted and weighted, this is not only because the support of the distributions is higher – their distributions are also skewed towards higher values, whereas the distributions of values for either subnetwork are more "evenly" distributed.

The effectiveness of the double penalization principle even for DeltaCon, suggests that the double penalization principle is still valuable even when a (dis)similarity can't be written as a convex combination of (dis)similarities for the positive and negative parts of the networks. DeltaCon has this problem because it considers "higher-order connectivities", for which in general $(A^+ + A^-)^n \neq (A^+)^n + (A^-)^n$ when $n \geq 2$.

#### S6 DISCUSSION

Section S6.1 summarizes S5. Section S6.2 discusses potential limitations to consider when interpreting the general applicability of the results. Section **??** reviews the contributions of this paper.

##### S6.1 Summary of Results

The results show that naive attempts to use pre-existing methods for comparing networks with unsigned edges can lead to useless or even actively misleading results when applied to networks with mixed-sign edge weights. They also show that considering changes in sign alone, or changes in magnitude alone, are usually insufficient. Finally, the results show how methods satisfying the double-penalization principle can avoid such misleading results, often by combining insight into changes in network structure from both positive and negative sign subnetworks.

##### S6.2 Limitations

Sections S6.2.1, S6.2.2, and S6.2.3 discuss the potential limitations of the chosen simulations. Section S6.2.1 discusses the number and variety of adversarial attacks for which these methods were tested.

Section S6.2.2 discusses the statistical properties of the random networks on which these methods were tested. Section S6.2.3 discusses the alternatives that these methods were tested against. Sections S6.2.4, S6.2.5, and S6.2.6 discuss the general limitations of the scope of this paper. Section S6.2.4 discusses the potential limited applicability of the convex combination decomposition property. Section S6.2.5 discusses which type of network comparison problems are addressed by this work. Section S6.2.6 discusses the limited applicability of this work for extending network comparison methods not discussed in this paper.

###### *S6.2.1 The Number of Investigated Adversarial Attacks Was Small*

These results only considered a limited number of adversarial attacks. The goal of this initial work was to consider adversarial attacks which did not affect the overall connectivity of the network and which affected limited aspects of the structure in obvious ways. One can think of many more changes to structure of mixed-sign networks for which we would want (dis)similarity measures to behave well. Future work can investigate whether double penalization leads to good responses for other kinds of adversarial attacks.

###### *S6.2.2 Distribution of Random Networks*

The goal was to consider a large number of networks with varying structure, such that their composite structure might be enough to encompass all "typical" networks. For example, always using a uniform distribution for the edge weights, rather than a Beta distribution with a varying hyperparameter, led to results (data not shown) that more reflected the particular nature of the uniform distribution rather than say anything about the network comparison methods themselves. For example, the spread of (dis)similarity values tended to be much smaller, and the effects of the magnitude swap attack were too predictably constrained. The chosen distribution led to mixed-sign networks with a wider range of properties.

The networks all had an expected sparsity of 50%, i.e. each possible edge was present or absent independently with probability 50%. This corresponds to an Erdos-Renyi model with parameter $p = \frac{1}{2}$. Future work might examine distributions with less uniform distributions of sparsity. Furthermore future work might want to choose magnitudes and signs more independently, and it might be worthwhile to look at magnitudes sampled from distributions which are not bounded (e.g. the exponential distribution). The Beta distributions have support limited to $(0,1)$ and we made an edge negative if its original magnitude fell in $(0, \frac{1}{2})$ and positive otherwise (see section S4.1 for reference).

Also one could argue that I had mostly generated random sparse matrices rather than true "random networks". While this concern seems unfounded given the one-to-one correspondence between networks with mixed-sign edge weights and their adjacency matrices (which could correspond to any square matrix), it is nevertheless true that networks with "realistic" connectivity have adjacency matrices whose sparsity pattern is non-generic. (In other words, matrices with no 0 entries are generic under typical random matrix distributions, which correspond to "complete" networks with every possible edge present.) I did use a distribution for which networks with many missing edges were generic, but still it could be argued that this did not correspond to "state of the art" or "typical" models for defining random network topologies.

I could have also considered Erdos-Renyi for other values of $p$, for example by treating $p$ as a random hyperparameter, or other models of random network topology. The case where $p = \frac{1}{2}$ corresponds to choosing all possible network topologies (unsigned skeletons) with equal probability, i.e. a uniform distribution, but nevertheless "most" such topologies may still be considered unrealistic compared to "typical real-world networks". The (dis)similarity methods chosen all adjust implicitly for sparsity somewhat, so it seems unlikely that any bias in the network topologies affects the validity of the overall conclusions. Nevertheless it would still be a valuable issue for future work to investigate.

###### *S6.2.3 Fairness of Comparisons*

One could argue that the shown results might make the proposed (dis)similarity measures look better unfairly because they were compared against "strawmen" alternatives. Nevertheless, with the exception of the raw spearman and the relative error, it seems most standard (dis)similarity measures that one would use for networks with less structure cannot be applied to networks with mixed-sign edge weights. Therefore to attempt to use such standard methods, it seems conceivable that many people in practice might disregard some of extra structure of networks with mixed-sign edge weights (e.g. by considering only the magnitude skeleton or the unsigned skeleton). So I believe it is fair to compare against such methods. Even to the extent such alternatives are unrealistic, comparing with them illustrates the understanding is lost when specific features of networks with mixed-sign edge weights are ignored, and thus can be justified from a "pedagogical" perspective. Future work might try to envision and make comparisons with more "realistic" or "fair" alternatives that don't obey the double penalization principle.

###### S6.2.4 Convex Combinations and Higher-Order Connectivity

A lot of emphasis was given to (dis)similarity measures that can be decomposed into a convex combination (weighted average) of their values for the positive edge subnetwork and the negative edge subnetwork. This emphasis may not be realistic, because such an approach does not seem to be generally applicable. (Dis)similarity measures which consider graph motifs involving two or more edges most likely require some reference to "higher-order adjacency matrices" $\mathbf{A}^n$ describing the number of paths between two nodes with $n$ edges. Although it is true that $\mathbf{A} = \mathbf{A}^+ - \mathbf{A}^-$, in general the "freshman's dream" is false, namely $\mathbf{A}^n = (\mathbf{A}^+ - \mathbf{A}^-)^n \neq (\mathbf{A}^+)^n - (\mathbf{A}^-)^n$ for $n \geq 2$. Cf. again section S5.4.3. Therefore I do not expect analogous properties to be applicable or available for general (dis)similarity measures considering graph motifs involving two or more edges. One could attempt to define reasonable convex combinations in terms of all $2^n$ terms of $(\mathbf{A}^+ - \mathbf{A}^-)^n$ when expanded, but such methods are unlikely to be computationally feasible (unless their computation reduces to something with far fewer terms).

Nevertheless, I do not believe that the failure of the convex combination paradigm to generalize to arbitrary (dis)similarity measures spells trouble for the Double Penalization Principle. Methods with the convex combination decomposition property are only a special case of methods satisfying the double penalization principle. The results for both unweighted and weighted DeltaCon appear to suggest that the double penalization principle remains useful even for (dis)similarity measures that consider "higher-order connectivities". Future work considering extensions to networks with mixed-sign edge weights of other (dis)similarity measures considering graph motifs with two or more edges would be worthwhile. For example, this work only considers extending the exact version of DeltaCon, but not also the approximate (and much more scalable) version of DeltaCon defined in (Koutra et al., 2016).

###### S6.2.5 Node Correspondence

Using the dichotomy between "known node correspondence" and "unknown node correspondence" network comparison methods as defined in (Tantardini et al., 2019), as mentioned before this work only considers the known node correspondence problem for networks with mixed-sign edge weights. The extent to which the double penalization principle or other ideas from this work are applicable as well to the unknown node correspondence problem for networks with mixed-sign edge weights is unclear. Future work investigating whether we need to start "from scratch" for the unknown node correspondence problem when it comes to principles for extending methods to networks with mixed-sign edge weights would be highly valuable. It seems conceivable that the ideas applicable for the known node correspondence problem might still be applicable at least indirectly to the unknown node correspondence problem, but this assumption really needs to be investigated and not taken for granted.

###### S6.2.6 General Methods

This work provides no "algorithm" or systematic procedure for extending arbitrary (dis)similarity measures to the case of networks with mixed-sign edge weights. The double penalization principle and the convex composition decomposition properties are merely "desired properties" or "design specifications" (the former "essential", the latter "optional"). I provide concrete implementations of these principles for only a handful of network comparison methods, and perhaps not even necessarily the most important ones. It seems unlikely that such a general systematic procedure could exist. For example, even amongst the three methods with the convex combination decomposition property, the formulas for the convex coefficients were different each time, following from "just so" arguments specific to each method.

### S7 APPENDIX: LEMMAS

The symbol $\mathbf{1}_I$ denotes the all ones vector $(1,\ldots,1)$ in $\mathbb{R}^I$, the notation $\hat{\mu}[\vec{\mathbf{x}}]$ denotes the arithmetic mean (empirical expectation) of $\vec{\mathbf{x}} \in \mathbb{R}^I$, $\hat{\mu}[\vec{\mathbf{x}}] := \frac{1}{I}\sum_{i=1}^{I}[\vec{\mathbf{x}}]_i$, and $\langle \vec{\mathbf{x}}, \vec{\mathbf{y}} \rangle$ denotes the standard inner product (a.k.a. the "dot product"), given $\vec{\mathbf{x}}, \vec{\mathbf{y}} \in \mathbb{R}^I$, $\langle \vec{\mathbf{x}}, \vec{\mathbf{y}} \rangle := \sum_{i=1}^{I}[\vec{\mathbf{x}}]_i \cdot [\vec{\mathbf{y}}]_i$.

#### S7.1 Mixed-Sign Spearman Correlation

**Lemma S7.1.** *Projection Lemma: when taking the (standard) inner product, "mean-centering one vector is as good as mean-centering both vectors":*

$$\begin{aligned}\langle \vec{\mathbf{x}} - \hat{\mu}[\vec{\mathbf{x}}] \cdot \mathbf{1}_I, \vec{\mathbf{y}} - \hat{\mu}[\vec{\mathbf{y}}] \cdot \mathbf{1}_I \rangle =& \langle \vec{\mathbf{x}} - \hat{\mu}[\vec{\mathbf{x}}] \cdot \mathbf{1}_I, \vec{\mathbf{y}} \rangle \\ =& \langle \vec{\mathbf{x}}, \vec{\mathbf{y}} - \hat{\mu}[\vec{\mathbf{y}}] \cdot \mathbf{1}_I \rangle .\end{aligned} \tag{S50}$$

*Here we assume $\vec{\mathbf{x}}, \vec{\mathbf{y}} \in \mathbb{R}^I$, and recall that $\mathbf{1}_I \in \mathbb{R}^I$ denotes the all 1's vector.*

**Proof:** Because inner products are symmetric,

$$\langle \vec{\mathbf{x}} - \hat{\mu}[\vec{\mathbf{x}}] \cdot \mathbf{1}_I, \vec{\mathbf{y}} - \hat{\mu}[\vec{\mathbf{y}}] \cdot \mathbf{1}_I \rangle = \langle \vec{\mathbf{y}} - \hat{\mu}[\vec{\mathbf{y}}] \cdot \mathbf{1}_I, \vec{\mathbf{x}} - \hat{\mu}[\vec{\mathbf{x}}] \cdot \mathbf{1}_I \rangle . \tag{S51}$$

Therefore it suffices to show only the first of the two claimed equalities, namely $\langle \vec{\mathbf{x}} - \hat{\mu}[\vec{\mathbf{x}}] \cdot \mathbf{1}_I, \vec{\mathbf{y}} - \hat{\mu}[\vec{\mathbf{y}}] \cdot \mathbf{1}_I \rangle = \langle \vec{\mathbf{x}} - \hat{\mu}[\vec{\mathbf{x}}] \cdot \mathbf{1}_I, \vec{\mathbf{y}} \rangle$, with the second equality following by applying symmetry (S51) before proceeding again with the steps of the proof of the first equality, and then applying symmetry (S52) once more:

$$\langle \vec{\mathbf{y}} - \hat{\mu}[\vec{\mathbf{y}}] \cdot \mathbf{1}_I, \vec{\mathbf{x}} \rangle = \langle \vec{\mathbf{x}}, \vec{\mathbf{y}} - \hat{\mu}[\vec{\mathbf{y}}] \cdot \mathbf{1}_I \rangle . \tag{S52}$$

Anyway the conclusion follows readily from the definitions and simple algebra, so providing a proof to this level of detail is probably an over-explanation making the result seem more confusing than it actually is. Nevertheless, at least for the sake of being thorough, below is a computation proving the first equality:

$$\begin{aligned}\langle \vec{\mathbf{x}} - \hat{\mu}[\vec{\mathbf{x}}] \cdot \mathbf{1}_I, \vec{\mathbf{y}} - \hat{\mu}[\vec{\mathbf{y}}] \cdot \mathbf{1}_I \rangle =& \langle \vec{\mathbf{x}} - \hat{\mu}[\vec{\mathbf{x}}] \cdot \mathbf{1}_I, \vec{\mathbf{y}} \rangle - \hat{\mu}[\vec{\mathbf{y}}] \cdot \langle \vec{\mathbf{x}} - \hat{\mu}[\vec{\mathbf{x}}] \cdot \mathbf{1}_I, \mathbf{1}_I \rangle \\ =& \langle \vec{\mathbf{x}} - \hat{\mu}[\vec{\mathbf{x}}] \cdot \mathbf{1}_I, \vec{\mathbf{y}} \rangle - \hat{\mu}[\vec{\mathbf{y}}] \cdot \left( \langle \vec{\mathbf{x}}, \mathbf{1}_I \rangle - \hat{\mu}[\vec{\mathbf{x}}] \cdot \langle \mathbf{1}_I, \mathbf{1}_I \rangle \right) \\ =& \langle \vec{\mathbf{x}} - \hat{\mu}[\vec{\mathbf{x}}] \cdot \mathbf{1}_I, \vec{\mathbf{y}} \rangle - \hat{\mu}[\vec{\mathbf{y}}] \cdot \left( \sum_{i=1}^{I} [\vec{\mathbf{x}}]_i - \hat{\mu}[\vec{\mathbf{x}}] \cdot I \right) \\ =& \langle \vec{\mathbf{x}} - \hat{\mu}[\vec{\mathbf{x}}] \cdot \mathbf{1}_I, \vec{\mathbf{y}} \rangle - \hat{\mu}[\vec{\mathbf{y}}] \cdot \left( \sum_{i=1}^{I} [\vec{\mathbf{x}}]_i - \sum_{i=1}^{I} [\vec{\mathbf{x}}]_i \right) \\ =& \langle \vec{\mathbf{x}} - \hat{\mu}[\vec{\mathbf{x}}] \cdot \mathbf{1}_I, \vec{\mathbf{y}} \rangle - 0 . \quad \square\end{aligned} \tag{S53}$$

For geometric intuition about why we should expect this lemma to be true, consider how $\hat{\mu}[\vec{\mathbf{x}}] \cdot \mathbf{1}_I$ is the projection[14] of $\vec{\mathbf{x}}$ onto the span of $\mathbf{1}_I$. Therefore mean-centering $\vec{\mathbf{x}}$ is the same as projecting[15] $\vec{\mathbf{x}}$ onto the orthogonal complement of (the span of) $\mathbf{1}_I$. Therefore the projection of $\vec{\mathbf{y}}$ onto the span of $\mathbf{1}_I$, namely $\hat{\mu}[\vec{\mathbf{y}}] \cdot \mathbf{1}_I$, is orthogonal to the mean-centered version of $\vec{\mathbf{x}}$. So in the expression $\langle \vec{\mathbf{x}} - \hat{\mu}[\vec{\mathbf{x}}] \cdot \mathbf{1}_I, \vec{\mathbf{y}} \rangle$, the contribution from $\hat{\mu}[\vec{\mathbf{y}}] \cdot \mathbf{1}_I$ cancels out:

$$\begin{aligned}\langle \vec{\mathbf{x}} - \hat{\mu}[\vec{\mathbf{x}}] \cdot \mathbf{1}_I, \vec{\mathbf{y}} \rangle =& \langle \vec{\mathbf{x}} - \hat{\mu}[\vec{\mathbf{x}}] \cdot \mathbf{1}_I, (\vec{\mathbf{y}} - \hat{\mu}[\vec{\mathbf{y}}] \cdot \mathbf{1}_I) + \hat{\mu}[\vec{\mathbf{y}}] \cdot \mathbf{1}_I \rangle \\ =& \langle \vec{\mathbf{x}} - \hat{\mu}[\vec{\mathbf{x}}] \cdot \mathbf{1}_I, \vec{\mathbf{y}} - \hat{\mu}[\vec{\mathbf{y}}] \cdot \mathbf{1}_I \rangle + \langle \vec{\mathbf{x}} - \hat{\mu}[\vec{\mathbf{x}}] \cdot \mathbf{1}_I, \hat{\mu}[\vec{\mathbf{y}}] \cdot \mathbf{1}_I \rangle \\ =& \langle \vec{\mathbf{x}} - \hat{\mu}[\vec{\mathbf{x}}] \cdot \mathbf{1}_I, \vec{\mathbf{y}} - \hat{\mu}[\vec{\mathbf{y}}] \cdot \mathbf{1}_I \rangle + 0 .\end{aligned} \tag{S54}$$

In general, if $\pi$ denotes an operator projecting onto some subspace, then for analogous reasons it is always true that $\langle \pi(\vec{\mathbf{x}}), \pi(\vec{\mathbf{y}}) \rangle = \langle \pi(\vec{\mathbf{x}}), \vec{\mathbf{y}} \rangle = \langle \vec{\mathbf{x}}, \pi(\vec{\mathbf{y}}) \rangle$.

[14] Herein, whenever I say "projection", I mean specifically "orthogonal projection".

[15] Likewise, herein "projecting" always refers to "orthogonally projecting".

**Lemma S7.2.** *As long as tied values are replaced with the mean of the tied values*[16]*, then any two rank vectors of the same length have the same mean.*

**Proof:** In the case that there are no ties, then it follows easily that all rank vectors of length $L$ have the same mean as the vector $(1,\dots,L)$. Because addition is commutative, if two vectors are the same up to a permutation of their entries, then they must have the same mean. In the absence of ties, all rank vectors by definition have entries which are permuted from the vector $(1,\dots,L)$.

In the case of tied values, we can use the general fact that replacing any subset $\mathcal{S} \subseteq [I]$ of the entries of a vector with the mean value of the entries in the subset does not change the mean of the vector. In the computation below, let $\vec{\mathbf{v}}$ denote the original vector, and let $\vec{\mathbf{v}}'$ denote the vector created by replacing the entries of $\vec{\mathbf{v}}$ corresponding to the subset $\mathcal{S}$ with their mean value:

$$\begin{aligned}
\hat{\mu}[\vec{\mathbf{v}}'] :=& \frac{1}{I}\sum_{i=1}^{I}[\vec{\mathbf{v}}']_i \\
=& \frac{1}{I}\left(\sum_{i\in\mathcal{S}}[\vec{\mathbf{v}}']_i + \sum_{i\notin\mathcal{S}}[\vec{\mathbf{v}}']_i\right) \\
=& \frac{1}{I}\left(\sum_{i\in\mathcal{S}}\left(\frac{1}{|\mathcal{S}|}\sum_{i\in\mathcal{S}}[\vec{\mathbf{v}}]_i\right) + \sum_{i\notin\mathcal{S}}[\vec{\mathbf{v}}]_i\right) \\
=& \frac{1}{I}\left(|\mathcal{S}|\cdot\left(\frac{1}{|\mathcal{S}|}\sum_{i\in\mathcal{S}}[\vec{\mathbf{v}}]_i\right) + \sum_{i\notin\mathcal{S}}[\vec{\mathbf{v}}]_i\right) \\
=& \frac{1}{I}\left(\sum_{i\in\mathcal{S}}[\vec{\mathbf{v}}]_i + \sum_{i\notin\mathcal{S}}[\vec{\mathbf{v}}]_i\right) \\
=& \frac{1}{I}\sum_{i=1}^{I}[\vec{\mathbf{v}}]_i \\
=& \hat{\mu}[\vec{\mathbf{v}}]\,.
\end{aligned} \tag{S55}$$

Applying this argument for each of the groups of tied elements, we get that the mean of a rank vector with tied values is the same as that of a rank vector without any ties, which in turn is the same as the mean of $(1,\dots,L)$. □

**Lemma S7.3.** ***Conditions for additivity of covariance for concatenated vectors:*** *Let $\vec{\mathbf{x}}_1,\vec{\mathbf{x}}_2 \in \mathbb{R}^I$, and $\vec{\mathbf{y}}_1,\vec{\mathbf{y}}_2 \in \mathbb{R}^J$. If at least one of*

- $\hat{\mu}[\vec{\mathbf{x}}_1] = \hat{\mu}[\vec{\mathbf{y}}_1]$*, or*
- $\hat{\mu}[\vec{\mathbf{x}}_2] = \hat{\mu}[\vec{\mathbf{y}}_2]$

*is true, then covariance is additive for $\vec{\mathbf{x}}_1 \oplus \vec{\mathbf{y}}_1$ and $\vec{\mathbf{x}}_2 \oplus \vec{\mathbf{y}}_2$, i.e.*

$$\widehat{\mathrm{cov}}[\vec{\mathbf{x}}_1 \oplus \vec{\mathbf{y}}_1, \vec{\mathbf{x}}_2 \oplus \vec{\mathbf{y}}_2] = \widehat{\mathrm{cov}}[\vec{\mathbf{x}}_1, \vec{\mathbf{x}}_2] + \widehat{\mathrm{cov}}[\vec{\mathbf{y}}_1, \vec{\mathbf{y}}_2]\,. \tag{S56}$$

**Proof:** Because covariance is symmetric, following reasoning analogous to that from the proof of Lemma S7.1, it suffices to give a proof only for the case that $\hat{\mu}[\vec{\mathbf{x}}_1] = \hat{\mu}[\vec{\mathbf{y}}_1]$. Note that $\hat{\mu}[\vec{\mathbf{x}}_1] = \hat{\mu}[\vec{\mathbf{y}}_1]$ implies $\hat{\mu}[\vec{\mathbf{x}}_1 \oplus \vec{\mathbf{y}}_1] = \hat{\mu}[\vec{\mathbf{x}}_1] = \hat{\mu}[\vec{\mathbf{y}}_1]$.

$$\begin{aligned}
\widehat{\mathrm{cov}}[\vec{\mathbf{x}}_1 \oplus \vec{\mathbf{y}}_1, \vec{\mathbf{x}}_2 \oplus \vec{\mathbf{y}}_2] =& \langle \vec{\mathbf{x}}_1 \oplus \vec{\mathbf{y}}_1 - \hat{\mu}[\vec{\mathbf{x}}_1 \oplus \vec{\mathbf{y}}_1]\cdot\mathbf{1}_{I+J}\ ,\ \vec{\mathbf{x}}_2 \oplus \vec{\mathbf{y}}_2 - \hat{\mu}[\vec{\mathbf{x}}_2 \oplus \vec{\mathbf{y}}_2]\cdot\mathbf{1}_{I+J}\rangle \\
=& \langle \vec{\mathbf{x}}_1 \oplus \vec{\mathbf{y}}_1 - \hat{\mu}[\vec{\mathbf{x}}_1 \oplus \vec{\mathbf{y}}_1]\cdot\mathbf{1}_{I+J}\ ,\ \vec{\mathbf{x}}_2 \oplus \vec{\mathbf{y}}_2\rangle \\
=& \langle \vec{\mathbf{x}}_1 \oplus \vec{\mathbf{y}}_1 - (\hat{\mu}[\vec{\mathbf{x}}_1]\cdot\mathbf{1}_I) \oplus (\hat{\mu}[\vec{\mathbf{y}}_1]\cdot\mathbf{1}_J)\ ,\ \vec{\mathbf{x}}_2 \oplus \vec{\mathbf{y}}_2\rangle \\
=& \langle (\vec{\mathbf{x}}_1 - \hat{\mu}[\vec{\mathbf{x}}_1]\cdot\mathbf{1}_I) \oplus (\vec{\mathbf{y}}_1 - \hat{\mu}[\vec{\mathbf{y}}_1]\cdot\mathbf{1}_J)\ ,\ \vec{\mathbf{x}}_2 \oplus \vec{\mathbf{y}}_2\rangle \\
=& \langle \vec{\mathbf{x}}_1 - \hat{\mu}[\vec{\mathbf{x}}_1]\cdot\mathbf{1}_I\ ,\ \vec{\mathbf{x}}_2\rangle + \langle \vec{\mathbf{y}}_1 - \hat{\mu}[\vec{\mathbf{y}}_1]\cdot\mathbf{1}_J\ ,\ \vec{\mathbf{y}}_2\rangle \\
=& \langle \vec{\mathbf{x}}_1 - \hat{\mu}[\vec{\mathbf{x}}_1]\cdot\mathbf{1}_I\ ,\ \vec{\mathbf{x}}_2 - \hat{\mu}[\vec{\mathbf{x}}_2]\cdot\mathbf{1}_I\rangle + \langle \vec{\mathbf{y}}_1 - \hat{\mu}[\vec{\mathbf{y}}_1]\cdot\mathbf{1}_J\ ,\ \vec{\mathbf{y}}_2 - \hat{\mu}[\vec{\mathbf{y}}_2]\cdot\mathbf{1}_J\rangle \\
=& \widehat{\mathrm{cov}}[\vec{\mathbf{x}}_1, \vec{\mathbf{x}}_2] + \widehat{\mathrm{cov}}[\vec{\mathbf{y}}_1, \vec{\mathbf{y}}_2]\,.
\end{aligned} \tag{S57}$$

[16]This is the way tied values are replaced by default in SciPy(Virtanen et al., 2020). "The average of the ranks that would have been assigned to all the tied values is assigned to each value".

The second and second-to-last equalities of (S57) use the Projection Lemma, Lemma S7.1. The third equality of (S57) uses $\hat{\mu}[\vec{\mathbf{x}}_1 \oplus \vec{\mathbf{y}}_1] = \hat{\mu}[\vec{\mathbf{x}}_1] = \hat{\mu}[\vec{\mathbf{y}}_1]$. The fourth equality of (S57) uses property (S17) from section S3.2.4. The fifth equality of (S57) uses property (S18) from section S3.2.4. The first and last equalities of (S57) are just the definition of (empirical) covariance, of course. □

**Lemma S7.4.** ***Subconvex Combination Decomposition Property:*** *For the coefficients $c_+, c_-$ from (S21) with the property that*

$$\rho_S^{\pm}(\mathcal{G}_1, \mathcal{G}_2) = c_+ \rho_S^+(\mathcal{G}_1, \mathcal{G}_2) + c_- \rho_S^-(\mathcal{G}_1, \mathcal{G}_2),$$

*we always have that $c_+ + c_- \leq 1$. (Trivially, $c_+, c_- \geq 0$.)*

**Proof:** For convenience, define the following variables:

$$\begin{aligned} a &:= \hat{\text{var}}\left[\mathcal{R}_{\mathcal{G}_1,\mathcal{G}_2}(\mathcal{G}_1^+)\right] \\ b &:= \hat{\text{var}}\left[\mathcal{R}_{\mathcal{G}_1,\mathcal{G}_2}(\mathcal{G}_1^-)\right] \\ c &:= \hat{\text{var}}\left[\mathcal{R}_{\mathcal{G}_1,\mathcal{G}_2}(\mathcal{G}_2^+)\right] \\ d &:= \hat{\text{var}}\left[\mathcal{R}_{\mathcal{G}_1,\mathcal{G}_2}(\mathcal{G}_2^-)\right]. \end{aligned} \tag{S58}$$

It follows then that the coefficients $c_+, c_-$ from (S21) equal

$$c_+ = \frac{\sqrt{ac}}{\sqrt{a+b} \cdot \sqrt{c+d}}, \quad c_- = \frac{\sqrt{bd}}{\sqrt{a+b} \cdot \sqrt{c+d}}. \tag{S59}$$

Thus the inequality $c_+ + c_- \leq 1$ is equivalent to

$$\frac{\sqrt{ac} + \sqrt{bd}}{\sqrt{a+b} \cdot \sqrt{c+d}} \leq 1 \quad \iff \quad \sqrt{ac} + \sqrt{bd} \leq \sqrt{a+b} \cdot \sqrt{c+d}. \tag{S60}$$

However this follows immediately from the Cauchy-Schwarz inequality, namely $|\langle \vec{\mathbf{u}}, \vec{\mathbf{v}} \rangle| \leq \|\vec{\mathbf{u}}\|_2 \|\vec{\mathbf{v}}\|_2$, if we define the vectors $\vec{\mathbf{u}} := (\sqrt{a}, \sqrt{b})$ and $\vec{\mathbf{v}} := (\sqrt{c}, \sqrt{d})$.

Another, equivalent way to show that the inequality (S60) is true is by using the AM-GM inequality applied to the vector $(ad, bc)$:

$$\begin{aligned} &\frac{ad+bc}{2} \geq \sqrt{abcd} \\ \iff &ad + bc + (ac + bd) \geq 2\sqrt{abcd} + (ac + bd) \\ \iff &(a+b)(c+d) \geq (\sqrt{ac} + \sqrt{bd})^2 \\ \iff &\sqrt{a+b} \cdot \sqrt{c+d} \geq \sqrt{ac} + \sqrt{bd}. \end{aligned} \tag{S61}$$

The analysis is somewhat simplified given that we can assume $a, b, c, d \geq 0$. □

#### S7.2 DeltaCon Distance

**Lemma S7.5.** *Any choice of $\varepsilon < \frac{1}{1+\|\mathbf{A}\|_\infty}$ is sufficient to guarantee that the Neumann series $(\mathbf{I} - \mathbf{W})^{-1}$ converges.*

**Proof:** To review, the $\infty$-operator (or "induced") matrix norm is the "maximum absolute row sum" of a matrix $\mathbf{M}$:

$$\|\mathbf{M}\|_\infty := \max_i \left[\sum_{j=1}^{J} |[\mathbf{M}]_{ij}|\right]. \tag{S62}$$

This operator norm is arguably one of the easiest to use to get a bound on the spectral radius because the definition of the degree matrix $\mathbf{D}$ from equation (S2) of section S2.2 involves the row sums of $\mathbf{A}$.

Recall from section S3.4 that the matrix $\mathbf{W}$ used to create the Neumann series $(\mathbf{I}-\mathbf{W})^{-1}$ is defined as

$$\mathbf{W} := \frac{\varepsilon}{1-\varepsilon^2}\mathbf{A} - \frac{\varepsilon^2}{1-\varepsilon^2}\mathbf{D}. \tag{S63}$$

Reviewing from section S3.4.1, to ensure that the Neumann series $(\mathbf{I}-\mathbf{W})^{-1}$ exists, our goal is to ensure that the spectral radius $\mathrm{sr}(\mathbf{W})$ of $\mathbf{W}$ is less than 1, $\mathrm{sr}(\mathbf{W}) < 1$, by ensuring the sufficient condition that $\|\mathbf{W}\|_\infty < 1$.

As long as $1-\varepsilon^2 > 0$, tedious algebra with the definition of $\mathbf{D}$ shows that

$$\|W\|_\infty = \frac{\varepsilon}{1-\varepsilon^2} \max_i \left[ \sum_{j\neq i} \left|[\mathbf{A}]_{ij}\right| + \left| [\mathbf{A}]_{ii} - \varepsilon \sum_{j=1}^{J} [\mathbf{A}]_{ij} \right| \right]. \tag{S64}$$

Therefore the requirement that $\|\mathbf{W}\|_\infty < 1$ is equivalent to the condition:

$$\frac{1-\varepsilon^2}{\varepsilon} \|\mathbf{W}\|_\infty = \max_i \left[ \sum_{j\neq i} \left|[\mathbf{A}]_{ij}\right| + \left| [\mathbf{A}]_{ii} - \varepsilon \sum_{j=1}^{J} [\mathbf{A}]_{ij} \right| \right] < \frac{1-\varepsilon^2}{\varepsilon}. \tag{S65}$$

Using the triangle inequality, we can bound $\frac{1-\varepsilon^2}{\varepsilon}\|\mathbf{W}\|_\infty$ by $(1+\varepsilon)\|\mathbf{A}\|_\infty$:

$$\begin{aligned} \frac{1-\varepsilon^2}{\varepsilon} \|\mathbf{W}\|_\infty &= \max_i \left[ \sum_{j\neq i} \left|[\mathbf{A}]_{ij}\right| + \left| [\mathbf{A}]_{ii} - \varepsilon \sum_{j=1}^{J} [\mathbf{A}]_{ij} \right| \right] \\ &\leq \max_i \left[ \sum_{j\neq i} \left|[\mathbf{A}]_{ij}\right| + \left|[\mathbf{A}]_{ii}\right| + \varepsilon \sum_{j=1}^{J} \left|[\mathbf{A}]_{ij}\right| \right] = (1+\varepsilon)\|\mathbf{A}\|_\infty. \end{aligned} \tag{S66}$$

Therefore it suffices to show that $(1+\varepsilon)\|\mathbf{A}\|_\infty < \frac{1-\varepsilon^2}{\varepsilon} = \frac{(1-\varepsilon)(1+\varepsilon)}{\varepsilon}$, which is equivalent to $\|\mathbf{A}\|_\infty < \frac{1-\varepsilon}{\varepsilon}$, which in turn (as can be shown via tedious algebra) is equivalent to the inequality $\varepsilon < \frac{1}{1+\|\mathbf{A}\|_\infty}$.

This satisfies $1-\varepsilon^2 > 0$ as long as $\|\mathbf{A}\|_\infty > 0$, i.e. $\mathbf{A} \neq \mathbf{0}$, where $\mathbf{0}$ denotes the all-zeros matrix (correpsonding to an empty network with no edges). However we can assume that $\mathbf{A} \neq \mathbf{0}$ without loss of generality, because when $\mathbf{A} = \mathbf{0}$, in turn $\mathbf{D} = \mathbf{0}$, implying that $\mathbf{W} = \mathbf{0}$, so that $(\mathbf{I}-\mathbf{W})^{-1} = \mathbf{I}^{-1} = \mathbf{I}$, so the Neumann series exists regardless of our choice of $\varepsilon$.

Therefore we have shown that $\varepsilon < \frac{1}{1+\|\mathbf{A}\|_\infty}$ is sufficient to guarantee that $\|\mathbf{W}\|_\infty < 1$, which in turn is sufficient to guarantee that $\mathrm{sr}(\mathbf{W}) < 1$, guaranteeing convergence of the Neumann series $(\mathbf{I}-\mathbf{W})^{-1}$. □

By seeking to upper bound the Frobenius norm by 1, and using very naive bounding techniques (mostly triangle inequality), it is also possible to prove that satisfying the much weaker bound:

$$\varepsilon < \frac{1}{1+\sqrt{S}||\mathbf{A}||_\infty} \tag{S67}$$

is sufficient to guarantee convergence of the Neumann series. It is probably possible to refine that argument to get a much better bound from the Frobenius norm, analogous to what was done in a much less general context in (Koutra et al., 2011).

It follows from the proof of Lemma S7.5 that if *at least one* of the two **strict** inequalities $\mathrm{sr}(\mathbf{W}) < \|\mathbf{W}\|_\infty$ or $\frac{1-\varepsilon^2}{\varepsilon}\|\mathbf{W}\|_\infty < (1+\varepsilon)\|\mathbf{A}\|_\infty$ is true, then we can use $\varepsilon = \frac{1}{1+\|\mathbf{A}\|_\infty}$ without any problems.

However, there do exist matrices $\mathbf{A}$ such that both $\mathrm{sr}(\mathbf{W}) = \|\mathbf{W}\|_\infty$ *and* $\frac{1-\varepsilon^2}{\varepsilon}\|\mathbf{W}\|_\infty = (1+\varepsilon)\|\mathbf{A}\|_\infty$. These should be the only matrices for which $\varepsilon = \frac{1}{1+\|\mathbf{A}\|_\infty}$ is too large. The general conditions under which $\mathrm{sr}(\mathbf{W}) = \|\mathbf{W}\|_\infty$ is true are obscure to me, but I do know that the second requirement that needs to be satisfied for a counterexample, $\frac{1-\varepsilon^2}{\varepsilon}\|\mathbf{W}\|_\infty = (1+\varepsilon)\|\mathbf{A}\|_\infty$, should be true if and only if all entries in the row of $\mathbf{A}$ with the largest sum of absolute values have the same sign (because then the absence of terms with opposite signs means there are no cancellations and thus that the triangle inequality holds with equality). Through happenstance I have found $\mathbf{A}$ that also satisfy the first requirement, that $\mathrm{sr}(\mathbf{W}) = \|\mathbf{W}\|_\infty$ when $\varepsilon = \frac{1}{1+\|\mathbf{A}\|_\infty}$.

**Lemma S7.6.** *There exist matrices* $\mathbf{A}$ *such that defining* $\varepsilon = \frac{1}{1+\|\mathbf{A}\|_\infty}$ *leads to* $\mathrm{sr}(\mathbf{W}) = 1$ *(implying that the Neumann series* $(\mathbf{I}-\mathbf{W})^{-1}$ *does not converge).*

**Proof:** Here is the matrix I found by happenstance that works

$$\mathbf{A} = \begin{bmatrix} 0 & 0 & 0 & 0 & 0 & -4.5 & 0 & 0 & 0 & 0 \\ 0 & 0 & 0 & 0 & 0 & 0 & 0 & 0 & 0 & 0 \\ 0 & 0 & 0 & 0 & 0 & 0 & 0 & 0 & 0 & -3.5 \\ 0 & 0 & 0 & 0 & 0 & 0 & 0 & 0 & 0 & 0 \\ 0 & 0 & 0 & 0 & 0 & 0 & 0 & 0 & 0 & 0 \\ -4.5 & 0 & 0 & 0 & 0 & 0 & 0 & 0 & 0 & 0 \\ 0 & 0 & 0 & 0 & 0 & 0 & 0 & 0 & 0 & 0 \\ 0 & 0 & 0 & 0 & 0 & 0 & 0 & 0 & 0 & 0 \\ 0 & 0 & 0 & 0 & 0 & 0 & 0 & 0 & 0 & 0 \\ 0 & 0 & -4.5 & 0 & 0 & 0 & 0 & 0 & 0 & 0 \end{bmatrix}. \tag{S68}$$

Rescaling a counterexample matrix by a constant does not appear to affect its status as a counterexample, nor does deleting (and/or adding) rows and/or columns consisting entirely of 0's. This leads to the simpler counterexample:

$$\mathbf{A} = \begin{bmatrix} 0 & 0 & 9 & 0 \\ 0 & 0 & 0 & 7 \\ 9 & 0 & 0 & 0 \\ 0 & 9 & 0 & 0 \end{bmatrix}, \tag{S69}$$

for which $\varepsilon = \frac{1}{1+\|\mathbf{A}\|_\infty} = \frac{1}{10}$, and thus

$$\mathbf{W} = \begin{bmatrix} -\frac{9}{99} & 0 & \frac{90}{99} & 0 \\ 0 & -\frac{7}{99} & 0 & \frac{70}{99} \\ \frac{90}{99} & 0 & -\frac{9}{99} & 0 \\ 0 & \frac{90}{99} & 0 & -\frac{9}{99} \end{bmatrix}. \tag{S70}$$

Very tedious calculations reveal that indeed the spectral radius of the matrix from (S70) is 1. Related counterexamples appear to also include

$$\mathbf{A} = \begin{bmatrix} 0 & \pm 9 \\ \pm 9 & 0 \end{bmatrix}, \quad \mathbf{A} = \begin{bmatrix} 0 & \pm 9 & 0 \\ \pm 9 & 0 & 0 \\ 0 & 0 & \pm 9 \end{bmatrix}, \quad \mathbf{A} = \begin{bmatrix} 0 & \pm 9 & 0 \\ \pm 9 & 0 & 0 \\ 0 & 0 & \pm x \end{bmatrix} \; (\forall x \text{ s.t. } |x| < 9). \tag{S71}$$

Discerning which operations will preserve the property $\mathrm{sr}(\mathbf{W}) = \|\mathbf{W}\|_\infty$ is subtle, since e.g. the following matrices appear to ***not*** be counterexamples:

$$\mathbf{A} = \begin{bmatrix} 0 & \pm x \\ \pm 9 & 0 \end{bmatrix}, \quad \mathbf{A} = \begin{bmatrix} 0 & \pm 9 \\ \pm x & 0 \end{bmatrix}, \quad \mathbf{A} = \begin{bmatrix} 0 & \pm x & 0 \\ \pm 9 & 0 & 0 \\ 0 & 0 & \pm 9 \end{bmatrix} \; (\forall x \text{ s.t. } |x| < 9). \tag{S72}$$

In any case, counterexamples clearly exist, but also appear to not be generic. □
